# Multi-omics Analyses Provide Insight into the Biosynthesis Pathways of Fucoxanthin in *Isochrysis galbana*

**DOI:** 10.1101/2021.11.28.470214

**Authors:** Duo Chen, Xue Yuan, XueHai Zheng, Jingping Fang, Gang Lin, Rongmao Li, Jiannan Chen, Wenjin He, Zhen Huang, Wenfang Fan, Limin Liang, Chentao Lin, Jinmao Zhu, Youqiang Chen, Ting Xue

**Affiliations:** The Public Service Platform for Industrialization Development Technology of Marine Biological Medicine and Products of the State Oceanic Administration, Center of Engineering Technology Research for Microalga Germplasm Improvement of Fujian, Fujian Key Laboratory of Special Marine Bioresource Sustainable Utilization, Key Laboratory of Developmental and Neural Biology, Southern Institute of Oceanography, College of Life Sciences, Fujian Normal University, Fuzhou 350117, China; Fujian Fishery Resources Monitoring Center, Fuzhou 350003, China

**Author notes:** Equal contribution (Duo Chen, Xue Yuan, XueHai Zheng, Jingping Fang). Corresponding author. E-mail address (Xue T).

**Keywords:** *Isochrysis galbana*, Fucoxanthin, Whole-genome duplication, Metabolome, Transcriptome

## Abstract

*Isochrysis galbana* is considered an ideal bait for functional foods and nutraceuticals in humans because of its high fucoxanthin (Fx) content. However, multi-omics analysis of the regulation networks for Fx biosynthesis in *I. galbana* has not been reported. In this study, we report a high-quality genome sequence of *I. galbana* LG007, which has a 92.73 Mb genome size, with a contig N50 of 6.99 Mb and 14,900 protein-coding genes. Phylogenomic inferences confirmed the monophyly of Haptophyta, with *I. galbana* sister to *Emiliania huxleyi* and *Chrysochromulina tobinii*. Evolutionary analysis revealed an estimated divergence time between *I. galbana* and *E. huxleyi* of ~ 133 million years ago (Mya). Gene family analysis indicated that lipid metabolism-related genes exhibited significant expansion, including *IgPLMT, IgOAR1* and Δ-4 desaturase. Metabolome analysis showed that the content of carotenoid in *I. galbana* cultured under green light for 7 days was higher than that of white light, and β-carotene was the main carotenoids, accounting for 79.09% of the total carotenoids. Comprehensive analysis of multi-omics analysis revealed that β-carotene, antheraxanthin, zeaxanthin, and Fx content was increased by green light induction, which was significantly correlated with the expression of *IgMYB98, IgZDS, IgPDS, IgLHCX2, IgZEP, IgLCYb,* and *IgNSY*. These findings contribute to understanding Fx biosynthesis and its regulation, providing a valuable reference for food and pharmaceutical applications.

## Introduction

Fucoxanthin (Fx) is widely distributed in algae and some invertebrate cells, including *Phaeodactylum tricornutum, Ectocarpus siliculosus, Thalassiosira pseudonana, Isochrysis galbana,* and *Nannochloropsis gaditana* [1,2]. Fx can be assembled with chlorophyll into Fx-chlorophyll protein (FCP) with certain proteins, and FCP exists in thylakoid membranes of algae and acts as a light-capturing antenna [3]. Fx has excellent blue-green light harvesting and photoprotection capabilities, which could help algae make full use of solar energy in different bands at different depths of seawater [3,4]. Fx, an oxygenated carotenoid, exhibits potential advantages with various pharmacological activities, including anti-inflammatory, anti-tumor, anti-obesity, anti-oxidant, anti-diabetic, anti-malarial, and anti-lipid effects [5]. However, the utilization of Fx as a nutraceutical in food and nutrient supplements is limited because of its low production level and poor stability. Microalgae are regarded as the most promising alternative Fx production algae with multiple biotechnological advantages, such as short growth cycle, easy handling, and large-scale artificial cultivation [6]. Compared with other microalgae, such as *Phaeodactylum tricornutum* and *Nannochloropsis gaditana*, *I. galbana* lacks cell walls and is much easier to be digested and handled, making it a good initial food source for the larvae of aquatic animals [7,8]. Moreover, *I. galbana* is a marine single-cell microalgae with rich in Fx (more than 10% of dry weight biomass) and lipid (7.0%-20% dry weight biomass), which is considered ideal material for the development of functional foods for humans [9]. Additionally, we found that the Fx content of *I. galbana* LG007 was the highest in different strains or species, which can be used as an ideal material for follow-up research (Figure S1).

Although the process of Fx synthesis has not been fully elucidated, several studies have attempted to reveal genes or proteins involved in Fx biosynthesis [10,11]. Genes involved in Fx biosynthesis have been identified, including β-carotene, phytoene synthase (*PYS*), phytoene desaturase (*PDS*), 15-cis-ζ-carotene isomerase (*ZISO),* ζ-carotene desaturase (*ZDS*), carotenoid isomerase (*CRTISO*), and lycopene β-cyclase (*LCYb*) [10,11]. However, some enzymes participating in the final step of Fx biosynthesis have not been discovered [11]. There are two generally accepted hypotheses for the final step of Fx biosynthesis from violaxanthin to Fx in Fx-producing algae: (1) violaxanthin is a precursor of Fx, which is converted by phycoxanthin, (2) neoxanthin is the precursor of Fx, which is formed by the ketonization of the neoxanthin and acetylation of an intermediate [12,13]. Studies have focused on the key genes related to the Fx synthesis pathway, mainly involved in the expression of some genes in the Fx synthesis pathway by external inducing factors (light intensity, methyl jasmonate, and arachidonic acid). Zhang et al. revealed the change in Fx content and gene expression pattern of the Fx synthesis pathway under conditions of high irradiance stress in the diatom *P. tricornutum,* showing an evident linear relationship between Fx content and the expression levels of *PYS* and zeaxanthin epoxidase (*ZEP*) [14]. Yu et al. reported that the expression level of *LCYb* could be significantly increased by treatment with methyl jasmonate and arachidonic acid, and the content of Fx in *P. tricornutum* was significantly higher than that in the control group, which showed that *LCYb* played an important role in the synthesis of Fx [15]. However, the aforementioned studies are mainly reflected in *P. tricornutum*, and few studies are on enzyme genes related to the Fx synthesis pathway of *I. galbana*. Because of limited genome information, how *I. galbana* regulates Fx biosynthesis at the DNA and RNA levels remains unclear. Draft genome sequences of *I. galbana* was generated in 2014 based on next-generation sequencing. However, incomplete genome assemblies produced short contigs and scaffolds, causing problems for the follow-up research of *I. galbana* [16]. Additionally, high-quality genomic resources can enable breeding novel *I. galbana* strains with higher Fx content in industrial practice for commercial use. But up to now, a systematic analysis of the regulatory networks for Fx biosynthesis in *I. galbana* has not been performed using genome, transcriptome and metabolome data according to our review of the literature.

In this study, we generated a high-quality genome assembly and annotation of *I. galbana* LG007 by using the third-generation sequencing (PacBio SEQUEL platform). The high-quality *I. galbana* LG007 genome provides a valuable resource about evolutionary events and genomic characteristics of aquatic algae. A study revealed the influence of spectral intensity and quality of blue-green light on Fx content in Chlorophyceae, but it did not involve the role of green light as a single source for Fx biosynthesis [17]. Our previous results suggested that Fx content could be increased under green light conditions, which is a special simulating factor that occurs during the cultivation of *I. galbana* (Figure S2). Transcriptomic and metabolomic analyses were performed on algae cells at different stages of cultivation (3, 5, 7, 9 days) under different light quality conditions (white and green) to reveal key genes or metabolic products that are potentially related to the accumulation and regulation of Fx biosynthesis.

## Results

### Genome sequencing and assembly

~ 15.5 Gb of PacBio long reads and 8.92 Gb of Illumina clean reads were generated (Table S1). The total length of all reads assembled from the *I. galbana* LG007 genome contained 353 contigs was 92.59 Mb, with a contig N50 of 666.66 kb, GC content of 58.44% and the longest contig length of 2.93 Mb (**Figure 1A, B**, Table S2, and Figure S3). The size of the assembled genome was close to that estimated by flow cytometry and 17-Kmer (Figures S4-S5). BUSCO analysis of our present assembly showed that ~ 83.8% of the plant orthologs were included in the assembled sequences (Table S3). Likewise, ~ 98.4% of Illumina clean reads and ~ 99.78% of PacBio long reads could be mapped to the genome, respectively (Tables S4-S5). These metrics implied that the assembled genome is credible and can be used for subsequent analysis. Using the 3D-DNA and LACHESIS workflow, 98.22% (90.95 Mb) of the genome was successfully anchored onto 15 superscaffolds (Figures S6-S7 and Table S6). Scaffold N50 of the *I. galbana* LG007 genome after high-throughput chromatin conformation capture (Hi-C) assisted assembly reached 6.99 Mb (Table S7), generating a high-quality genome assembly for *I. galbana*.

**Figure 1.**
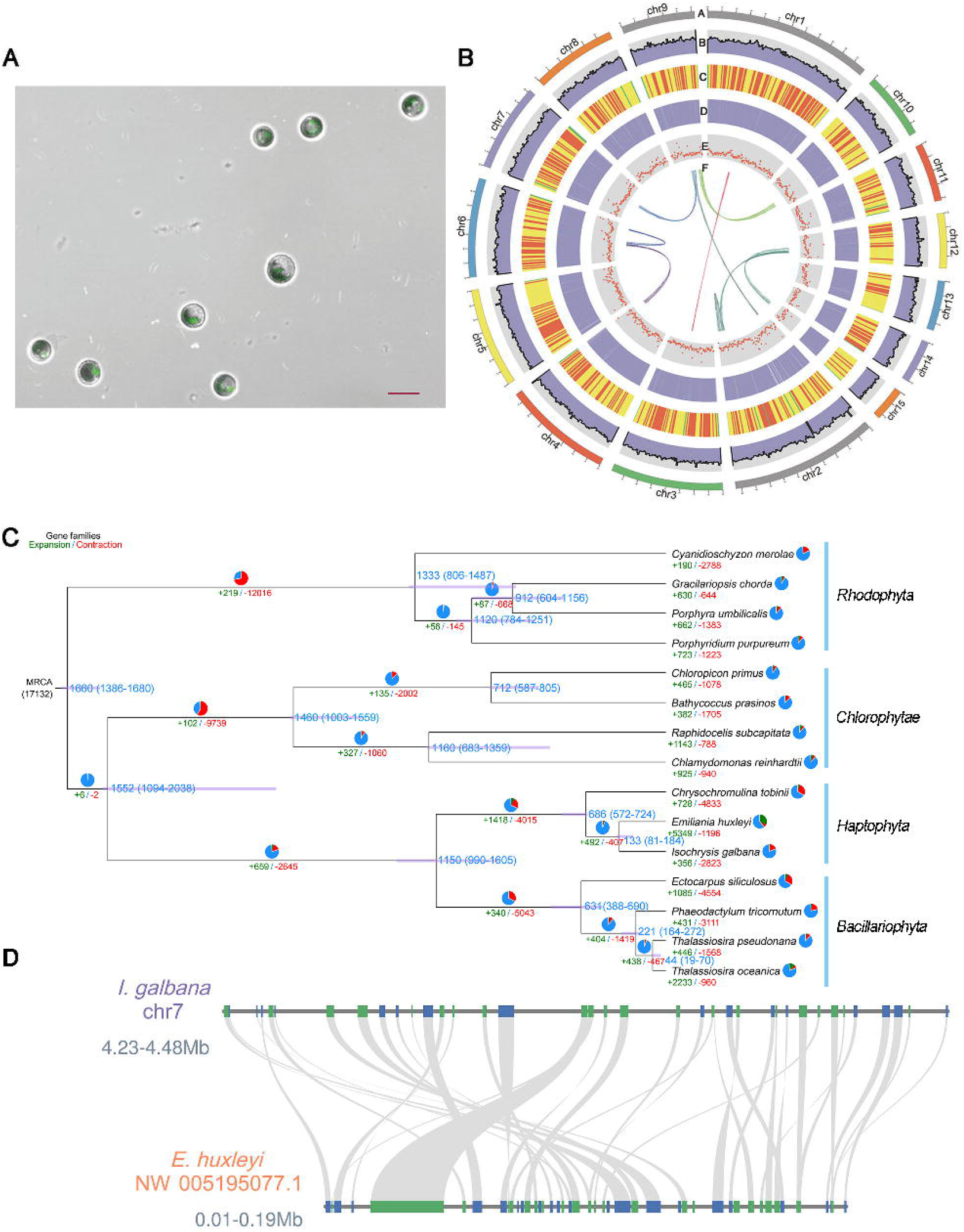
Genomic characteristics of *I. galbana*. **A.** Confocal laser scanning microscopic images (bottom; bar represents 20 μm) of *I. galbana* cells. **B.** Images of *I. galbana* assembly (a assembled superscaffolds. b distribution of GC content. c density of gene. d expression values. e percent coverage of TEs in nonoverlapping windows. f syntenic blocks within the genome). **C** Phylogenetic tree of *I. galbana* and 14 species. Green and red numbers represent expanded and contracted gene families, respectively. The estimated divergence time (Mya) is denoted at each node with blue font. **D** Microsynteny analysis of *I. galbana* superscaffolds and *E. huxleyi* scaffolds.

### Gene prediction and annotation

A total of 14,900 protein-coding genes were predicted in *I. galbana* LG007 genome by combining the homology search results, *de novo* prediction, and transcriptome evidence. The protein-coding genes had an average gene length of 1789 bp, and an average coding sequence (CDS) length of 1428 bp (Table S8). We functionally annotated 9161, 12,469, 3773, 4977, and 9161 genes to EggNOG, Non-Redundant (NR), Kyoto Encyclopedia of Genes and Genomes (KEGG), Gene Ontology (GO), and Clusters of Orthologous Groups (COG), leading to ~ 83.89% (12,500 genes) of the total genes with at least one match to the known public database (Table S9). A total of 439 transcription factors (TFs) distributed in 20 families, including 198 protein kinase families, 55 *HSF,* 49 *ZFWD,* and 44 *MYB* superfamily proteins (Table S10). In addition, we also identified 95 tRNAs, 58 rRNAs, and 4 snRNAs in an *I. galbana* LG007 genome (Table S11). ~ 46.82% of the assembled *I. galbana* LG007 genome comprised repetitive sequences. Long terminal repeat (LTR) retrotransposons spanning 15.36% of the assembled genome with 1.08% Ty1/Copia and 4.53% Ty3/Gypsy. Non-LTR elements accounted for 12.54% of the genome, including 11.62% long interspersed nuclear elements (LINEs) and 0.91% short interspersed nuclear elements (SINEs). Tandem Repeats Finder identified over 43,633 tandem repeats, spanning 4.3% of the *I. galbana* LG007 genome (Table S12).

### Phylogenetic evolution of the *I. galbana* genome

A total of 179 single-copy homologous genes were identified among 15 genomes by using OrthoFinder (version 2.3.12) and were used to reconstruct a phylogenetic tree (Figure 1C). Phylogenetic analysis showed that *I. galbana* diverged into the Haptophyta branch ~ 133 Mya after the divergence of the Rhodophyta (1333 Mya), Chlorophytae (1460 Mya), and Bacillariophyta (1150 Mya). These results support the view that *E. huxleyi, I. galbana* and *C. tobinii* as monophyletic groups share a common Haptophyta ancestor [18–20]. Although *E. huxleyi* and *I. galbana* genomes appear to share only limited collinearity, this observation may stem from the lower quality of the *E. huxleyi* genome assembly (Figure 1D). The results showed that *I. galbana* and *E. huxleyi* may have a close relationship and are sisters in coccolithophores, which is consistent with the findings of phylogenetic analysis. Comparative genomic analysis showed that gene family expansions outnumbered contractions in *Raphidocelis subcapitata, Thalassiosira oceanica,* and *E. huxleyi*. We also discovered 356 expanded and 2823 contracted gene families in *I. galbana* LG007. KEGG pathway analysis showed that the expanded gene families were specifically enriched in signal transduction, purine metabolism, lipid metabolism, and ABC transporters were enriched in the expanded genes (Table S13). GO analysis showed that these expanded genes were related to signaling, metabolic processes, stimulus response, and catalytic activity (Table S14). Interestingly, lipid metabolism-related genes (*IgPLMT, IgOAR1,* and Δ-4 desaturase) exhibit significant expansion, indicating that the expansion of these genes in *I. galbana* could enhance the regulation and biosynthesis of carotenoid, resulting in a high content of Fx in *I. galbana*. A total of 2823 contracted gene families highlighted the functions pertaining to signal transduction, starch metabolism, and biosynthesis of other secondary metabolites (Table S15). GO terms of the contracted genes were associated with binding, catalytic activity, transporter activity, transcription regulator activity, stimulus-response, signaling, and biosynthesis of secondary metabolites (Table S16). Thus, it is likely that the contractions of secondary metabolites biosynthesis and stimulus-related genes could affect the accumulation of other secondary metabolites and resistance in *I. galbana,* resulting in a relatively good photoprotection capabilities only in the blue-green light of seawater. A comparison of *E. huxleyi, C. tobinii, P. tricornutum, C. reinhardtii,* and *I. galbana* LG007 revealed that 2027 (31.37%) of the 12,387 *I. galbana* LG007 gene families were common to other four species, whereas 3135 gene families were specific to *I. galbana* LG007 (**Figure 2A**). GO enrichment analysis showed that the functions of these specific genes mainly included metabolic process, catalytic activity, biological regulation, stimulus response, developmental process, binding, pigmentation, molecular transducer activity, and transcription regulator activity (Table S17). KEGG enrichment analysis showed enrichment of the calcium signaling pathway, the cGMP-PKG signaling pathway, fatty acid biosynthesis, signal transduction, energy metabolism, and terpenoid metabolism (Table S18).

**Figure 2.**
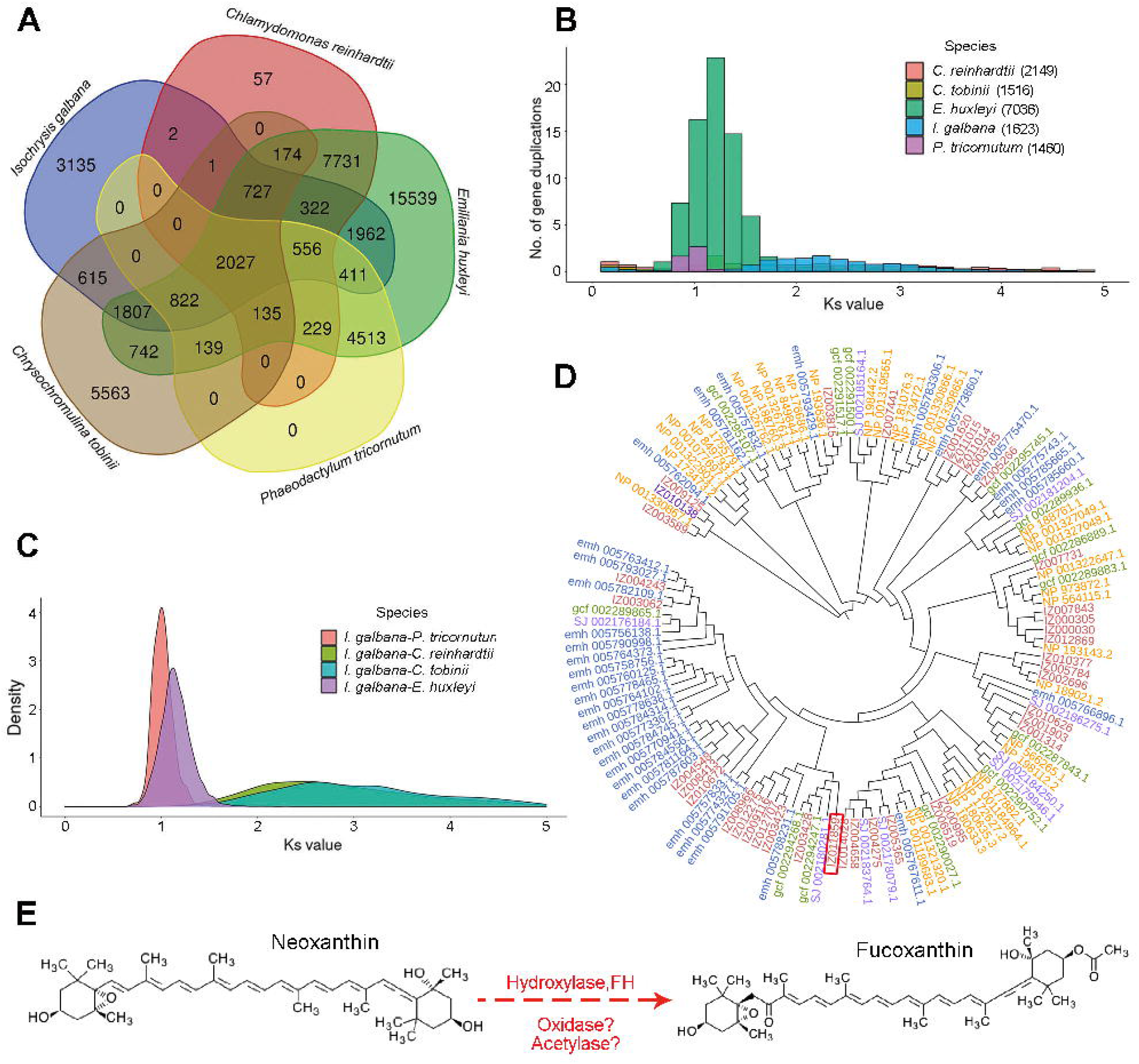
Evolution of the *I. galbana* LG007 genome. A. Shared and unique gene families among five species. **B.** *K*s distributions for duplicated gene pairs in *I. galbana, P. tricornutum, C. reinhardtii, E. huxleyi,* and *C. tobinii*. **C.** Distribution of the *K*s between the *I. galbana, P. tricornutum, C. reinhardtii, E. huxleyi,* and *C. tobinii*. **D.** Evolutionary tree of hydroxylase genes in *I. galbana* (IZ), *P. tricornutum* (SJ), *E. huxleyi* (emh), *C. tobinii* (gcf), and *A. thaliana* (NP). Evolutionary tree of hydroxylase gene family was constructed by RAxML software. The red, purple, blue, green, and yellow fonts represent the hydroxylase genes from *I. galbana, P. tricornutum, E. huxleyi, C. tobinii,* and *A. thaliana,* respectively. Fucoxanthin hydroxylase gene (*IgFH,* IZ011859) is marked with red box. **E.** Chemical reactions possibility in the process of synthesis according to the structure of diadinoxanthin and Fx.

To infer the whole-genome duplication (WGD) event in *I. galbana*, we calculated the synonymous substitution rate (*K*s) by a mixture model implemented in the R package. 2149, 1516, 7038, 1460, and 1623 genes were used to calculate the *Ks* value for *C. reinhardtii, C. tobinii, E. huxleyi, P. tricornutum,* and *I. galbana,* respectively. The sharp peak of distribution in *I. galbana* has a median *K*s of approximately 2.53, which is higher the ortholog divergences of *I. galbana* and *P. tricornutum* (*K*s, ~0.95), and the divergences of *I. galbana* and *E. huxleyi* (*K*s, ~1.38) (Figure 2B, C). Comparative analysis of *I. galbana* and other four species genomes provided evidence of the two WGD events according to the sharp peaks in synonymous substitution rate (approximately 0.95 and 2.53) values (Figure 2C). The distribution of *K*s values suggested that divergence between *I. galbana* and *E. huxleyi* occurred at ~ 133 Mya (*K*s, ~ 1.38), later than the ancient WGD in *I. galbana* (*K*s, ~ 2.53), which was dated at ~ 245 Mya. This finding indicated that *I. galbana*, *P. tricornutum*, and *E. huxleyi* experienced long-term divergence and speciation.

### Analysis of gene family related to Fx pathway in *I. galbana*

The metabolic processes of violaxanthin, neoxanthin, diadinoxanthin and Fx may be very complicated, which involved in oxidation, isomerization, acetylation, deepoxidation, hydrogenation, and hydroxylation chemical reactions according to the structure of substrates or products. To efficiently identify candidate genes involved in Fx biosynthesis, we identified 39 hydroxylase genes by a combination of direct screening of the genome assembly annotations and conserved domain BLAST searches. One of the 39 hydroxylase genes catalyzed the hydroxylation of hydrophobic substrates, which has a similar chemical reaction according to the structure of diadinoxanthin, neoxanthin, and Fx, suggesting that the fucoxanthin hydroxylase gene (*IgFH*, IZ011859) might have a function similar to that of the characterized neoxanthin-Fx or diadinoxanthin-Fx as candidate genes (Figure 2D, E and Figure S8). We could not detect Fx or new products after incubating recombinant IgFH proteins with diadinoxanthin by enzyme activity assay. However, overexpression and enzyme activity assays confirmed that neoxanthin could be further catalyzed by the IgFH proteins, implying that *IgFH* could play a key role in Fx biosynthesis (Figure S9).

Two distinct interconversion cycles of zeaxanthin to violaxanthin (*VDE, ZEP*) and diatoxanthin to diadinoxanthin (*DDE*, *DEP*) containing epoxidase and de-epoxidase were involved in the same type of catalytic reaction. We identified 6, 16, 4, and 1 epoxidase family proteins in the *I. galbana*, *A. thaliana*, *P. tricornutum,* and *C. reinhardtii,* respectively. The number of epoxidase gene proteins from *I. galbana* (6) was lower than that of *A. thaliana* (16), but was close to that of *P. tricornutum* (4) (Figures. S10-S12). Additionally, we constructed a phylogenetic tree based on the identified de-epoxidase proteins from 15 amino acid sequences of *I. galbana* (5), *A. thaliana* (4), *P. tricornutum* (5), and *C. reinhardtii* (1). The number of de-epoxidase gene families in *I. galbana* (5) was significantly higher than that in *C. reinhardtii* (1). We speculated that the presence of most epoxidase and de-epoxidase gene proteins could play a role in the Fx metabolic synthesis of *I. galbana* (Figures S13-S15).

### Identification of differential expressed genes related to Fx pathway in *I. galbana*

Among the annotated genes, 12,093 (96.74%) genes were expressed in 24 samples. Some genes were highly expressed on day 7 (7d) with green light (Figure S16), which has a similar trend to the phenotype of Fx content (Figure S17). To explore the differentially expressed genes (DEGs) involved in the Fx biosynthesis, DEGs of pairwise comparisons between control and treatment were analyzed (e.g., control 3d vs treated 3d, control 5d vs treated 5d). The number of stage-specific genes (fragments per kilobase of exon per million fragments mapped, FPKM ≥ 8000) varied from 646 to 807 for the control group and 448 to 802 for the treated group (Figure S18). The number of stage-specific genes in the treatment group changed little in the 7d, but decreased significantly at 9d. The number of stage-specific genes in the control group fluctuated in the 7d and increased sharply at 9d (Figure S18). These results indicate that 7d would be an important period for Fx biosynthesis. GO enrichment analyses of these stage-specific genes between the control and treated groups showed a representation of genes associated with various biological regulation, carboxylic acid biosynthetic process, fatty acid metabolic process, stimulus response, and catalytic process (Figure S19).

In total, 3730 genes exhibited significantly higher expression, and 4089 genes exhibited significantly lower expression at different stages in the treated group than in the control group (**Figure 3A**). Among these comparisons, the up-regulated and down-regulated expression between the control and treatment groups was greatest at 5d and 7d, indicating a difference in the transcription levels at the 5d and 7d stage with green light irradiation. Some TFs also exhibited a significant difference between the control and treatment groups, for example, the members of the *MYB, HB,* and *HD-ZIF* families involved in pigment accumulation and resistance stress showed significantly higher expression (Figure 3B). Fatty acid elongation, steroid biosynthesis, signaling, stimulus response, and cell development were enriched in the DEGs, particularly at the 7d stage with green light irradiation (Figure 3C). To explore the metabolic pathways responsible for the differences between the control and treated groups, we analyzed the expression profiles of DEGs using the MapMan tool. We found that the genes involved in lipid metabolism, light reactions, and pyruvate oxidation were more active in the treated group at the 7d stage, suggesting higher energy and more synthetic substrates for the metabolism of terpenoids and the β-carotene pathway (Figure 3D). With regard to MYB proteins, 52 and 146 MYB proteins were identified in *I. galbana* and *A. thaliana,* respectively (**Figure 4A** and Figure S20). R2R3-MYB TFs are related to the biosynthesis of pigment, suggesting a close relationship with the accumulation of Fx in *I. galbana* under light-induced conditions. Among genes with a relatively high expression of 114 TFs, we found that *IgMYB98* (IZ007092) is an R2R3-MYB transcription factor, which is significantly down-regulated (*p* = 1.99E-09) in the synthesis of Fx and may be a key gene for negative regulation of Fx biosynthesis in *I. galbana* under light-induced conditions (Figure 4B and Table S19).

**Figure 3.**
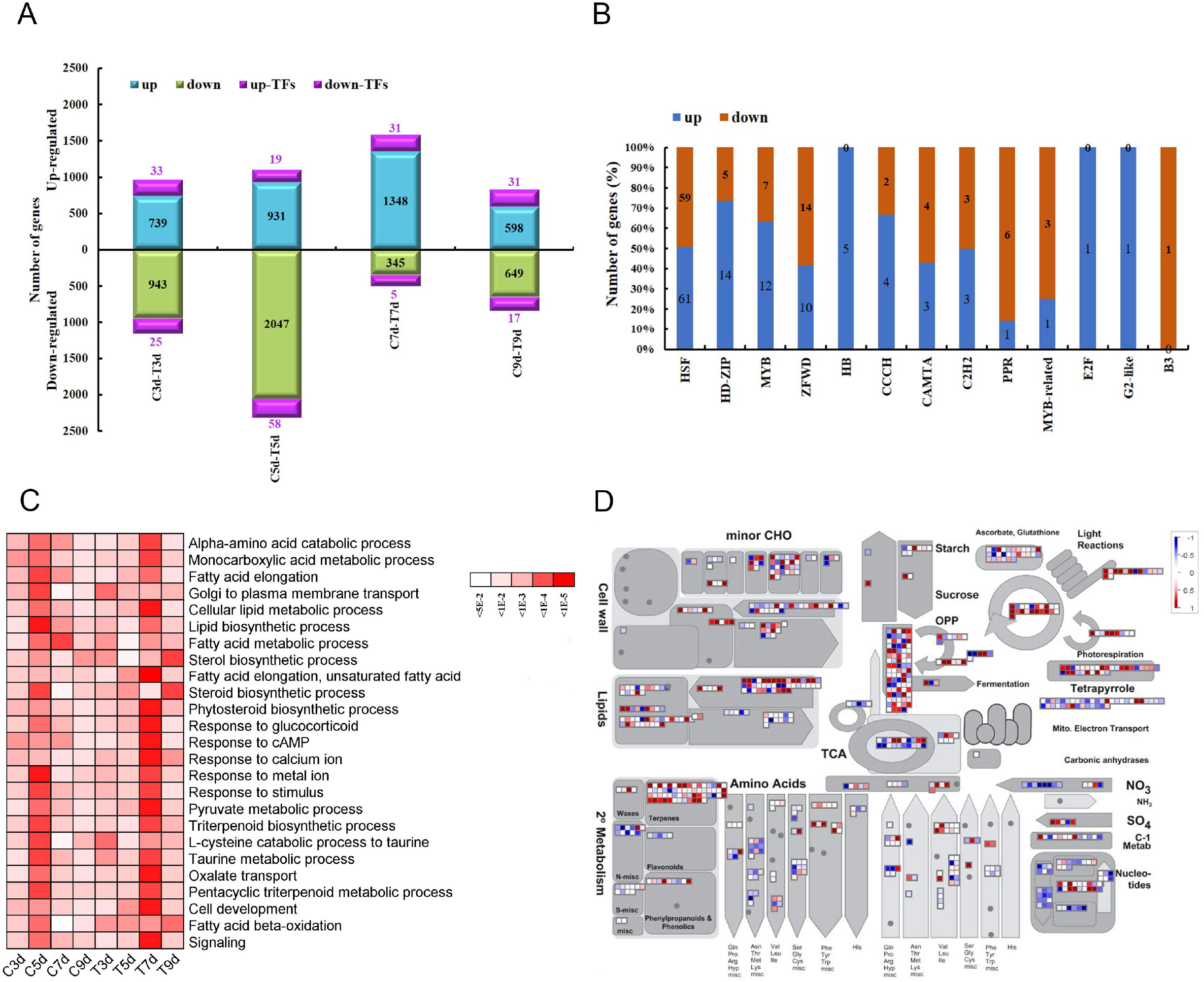
Differential gene expression in the treated group as compared with control group at different stages. **A.** Number of up- and down-regulated genes at different cultivation times in the treated group (green light) and control group (white light). The number of up- or down-regulated TFs at different cultivation times in the treated group is also given. **B.** Number of different TF families showing up- or down-regulation in the treated group. **C.** GO terms analysis of DEGs (biological process) at different cultivation times in the control and treated groups. The color scale on the right indicates significance (corrected *p-value*). **D.** Metabolic pathways with differential expression profile in treated group as compared with control group at 7d. DEGs between the treated group and control group at 7d were loaded into the MapMan software. Red and blue colors indicate high and low expression, respectively.

**Figure 4.**
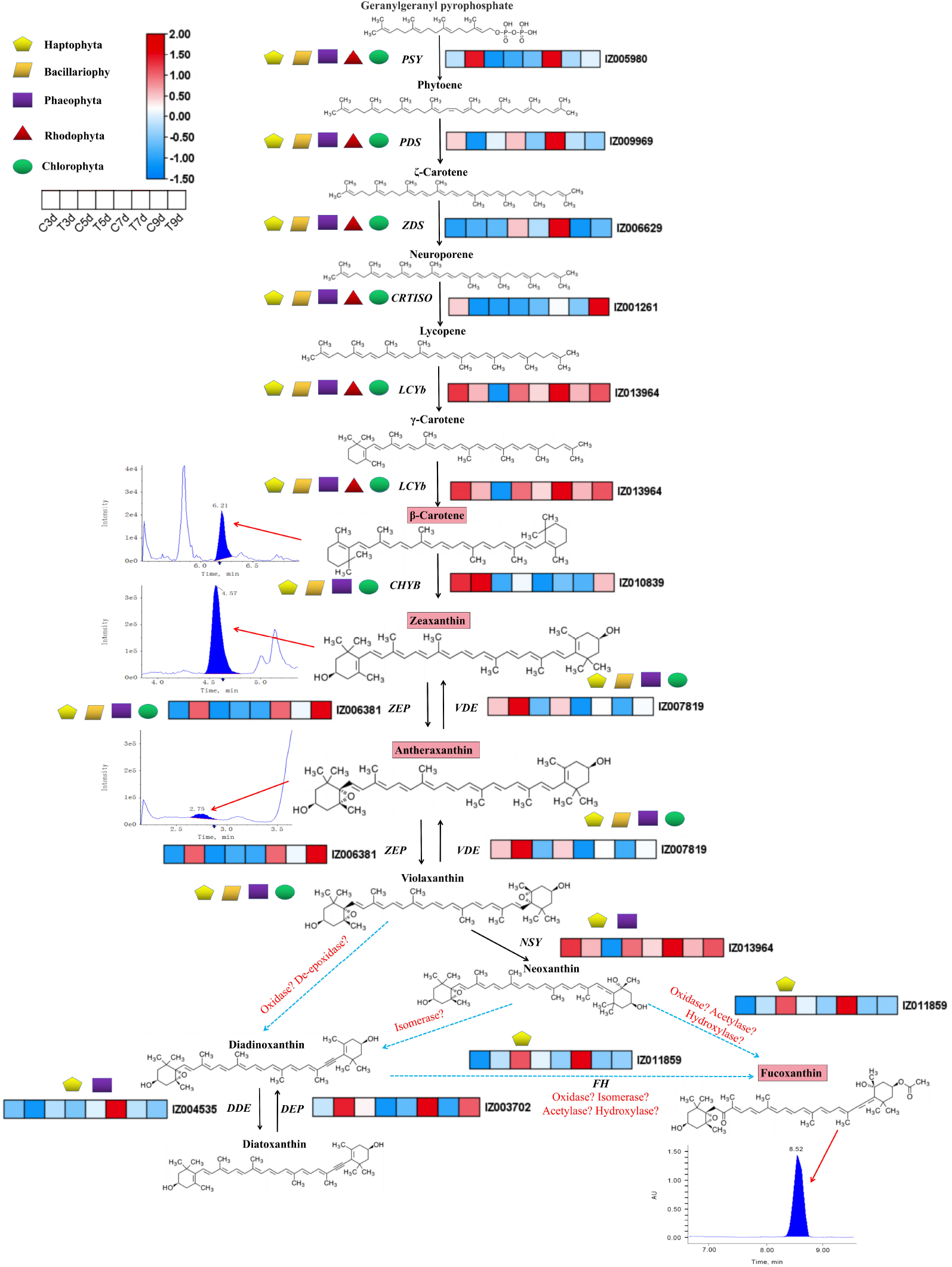
Transcriptomic and metabolomic analysis of fucoxanthin accumulation in *I. galbana*. **A.** Evolutionary tree of MYB box genes in *I. galbana* (IZ) and *A. thaliana* (At). **B.** Heat map of the DEGs in the treated group and control groups at different stages. Each box represents an individual gene, and the red and blue colors indicate high and low expression of gene, respectively. **C.** Heat map of the changes of carotenoid in *I. galbana* both in the white and in the green light. **D.** Content of major carotenoid in *I. galbana* under different light qualities. Error bars indicate SDs from three replicates. Capital letters and small letters indicate that significance is at the *0.01* or *0.05* level, respectively.

### Metabolic differences under different light qualities

In order to understand the biosynthesis pathways of the Fx accumulation under different light qualities (white and green light), two samples (7d-G, cultivate for 7d with green light; 7d-W, cultivate for 7d with white light) were collected at 7d. Fifteen carotenoids were identified in the comparison of 7d-G vs.7d-W groups, including three types of carotenes, carotenoid esters, and xanthophylls (Figure 4C and Table S20). Phytoene, ζ-carotene, neurosporene, lycopene, γ-carotene, violaxanthin, and neoxanthin involved in carotenoid biosynthesis were not detected between the 7d-G and 7d-W groups, indicating that these types of carotenoids may be prone to degradation or rapid conversion in *I. galbana*. The main carotenoids that accumulated in the 7d-G group were β-carotene, echinenone, violaxanthin-myristate, zeaxanthin, and β-cryptoxanthin, among which β-carotene was the main carotenoid, accounting for 79.09% of the total carotenoid. Echinenone had the second-highest content in the 7d-G and 7d-W groups, accounting for 8.65% and 7.17% of the total carotenoid content, respectively. Heat map analysis revealed that the content of carotenoid in the 7d-G group was significantly higher than that in the 7d-W group, including β-carotene (1.64-fold increase), lutein-myristate (1.56-fold increase), β-cryptoxanthin (1.29-fold increase), capsanthin (2.43-fold increase), and zeaxanthin (2.67-fold increase) (Figure 4D and Figure S17B). We identified eight differential accumulated carotenoids (DACs) in 7d-W vs. 7d-G, including seven up-regulated DACs and one down-regulated DACs. The number of up-regulated DACs was much higher than that of down-regulated DACs in the comparison of 7d-G vs.7d-W, suggesting the abundant diversity of carotenoid present under green light. Notable increases in carotenoid from the 7d-W to 7d-G samples included those in ε-carotene (2.28-fold increase, *p* = 0.008), violaxanthin-laurate (3.50-fold increase, *p* = 0.003), violaxanthin-myristate (2.58-fold increase, *p* = 0.008), antheraxanthin (2.86-fold increase, *p* = 0.02), capsanthin (2.56-fold increase, *p* = 0.001), zeaxanthin (2.67-fold increase, *p* = 0.002), and Fx (2.14-fold increase, *p* = 0.009) (Table S21). These results showed that green light had a significant effect on the metabolism of carotenoid in *I. galbana*.

### Gene co-expression network involved in Fx accumulation

To identify the hub genes, we performed weighted gene co-expression network analysis (WGCNA) for the control and treated groups separately. Twenty-five modules (comprising 31-2830 genes) were identified in the control group, and 24 modules (comprising 30-3333 genes) were recognized in the treated group (**Figure 5A, D** and Figure S21). Notably, the red co-expression module of the control group and turquoise co-expression module of the treated group showed a relatively high correlation (*r* ≥ 0.60) with Fx content (Figure 5B, E). GO and KEGG pathway enrichment analysis of relatively higher correlation modules highlighted key DEGs and biological processes with Fx content (Figure 5C, F). For example, the GO and KEGG analyses showed that the red module of the control group included most of the genes involved in metabolic processes, stimulus response, biological regulation, biosynthetic process, biosynthesis of secondary metabolites, fatty acid biosynthesis, and carotenoid biosynthesis (Figures S22-S23). The turquoise module associated with green light irradiation for Fx content showed enrichment of GO terms and KEGG pathways related to biological process, metabolic process, biosynthetic processes, catabolism processes, metabolic pathways, biosynthesis of secondary metabolites, and carbon metabolism (Figures S24-S25). Next, we studied the preservation of co-expression modules between the control and treated groups (Figure S26). We identified a midnight-blue module (35 genes) between the control and treated groups, and the harbored genes of module was enriched in metabolic processes, negative regulation of biological process, and stimulus response (Figure S26 C, D). Taken together, hub gene analysis identified ζ-carotene desaturase (*IgZDS,* IZ006629), phytoene desaturase (*IgPDS,* IZ009969), and Fx-chlorophyll a (*IgLHCX2,* IZ013244) in the red, turquoise, and midnight-blue modules, which are involved in the biosynthetic pathway of β-carotene (Fx synthesis).

**Figure 5.**
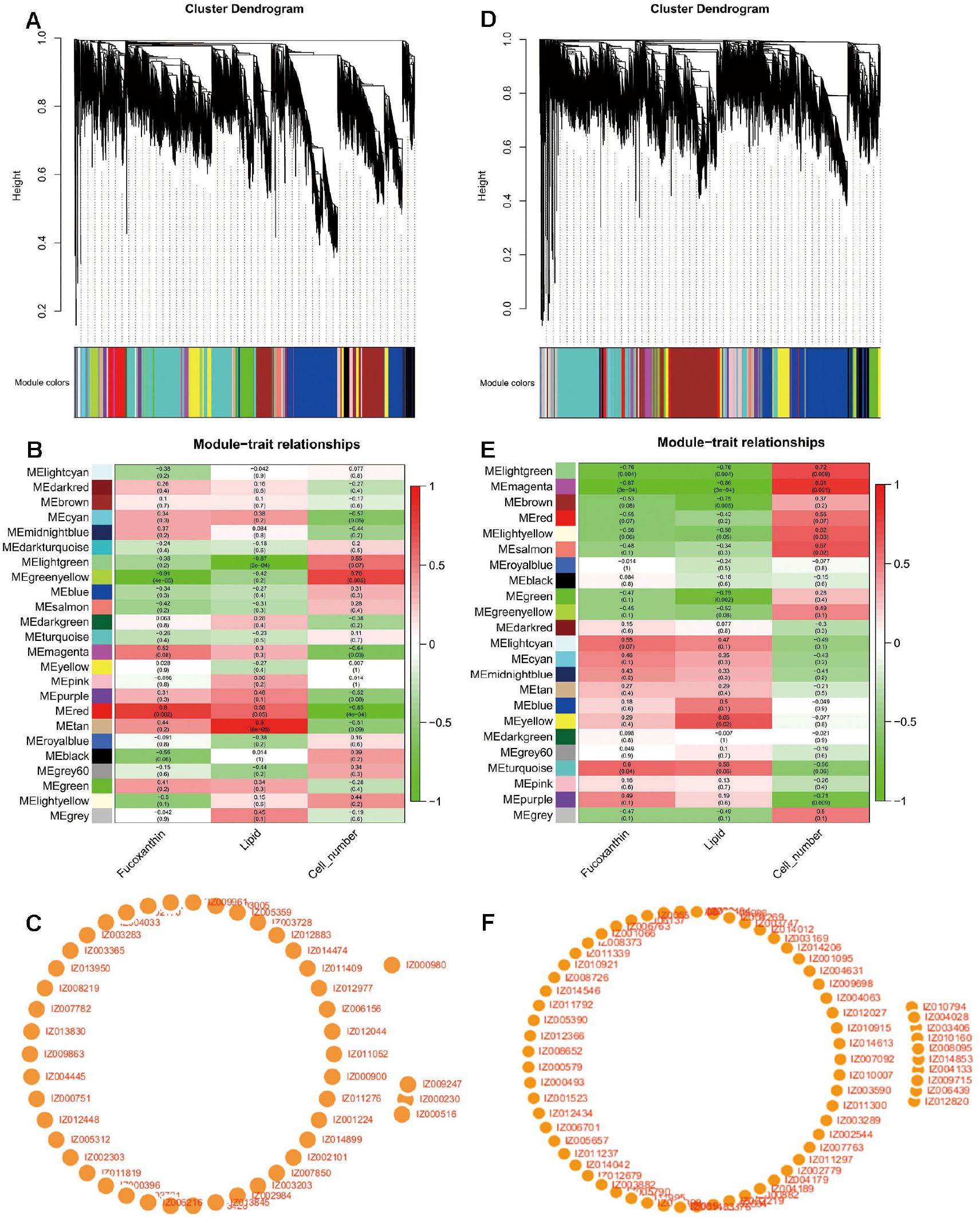
Co-expression network during Fx accumulation under different light qualities. **A.** Hierarchical clustering from WGCNA in control group. **B.** Heatmap plot of the correlation of modules. **C.** Transcriptional regulatory network between genes and module membership in control group. **D.** Hierarchical clustering from WGCNA in treated group. **E.** Heatmap plot of the correlation of modules. **F.** Transcriptional regulatory network between genes and module membership in treated group. Red represents positive correlation, and green represents negative correlation.

### Integrated transcriptomic and metabolomic analyses

To explore the relationship between genes and metabolites involved in Fx synthesis under different light qualities (white and green lights), the pathway of DEGs and DACs related to Fx was constructed (Figure 6). Genes involved in the Fx biosynthetic pathway exhibited a very high expression in the treated group at the 7d stage according to the transcriptome data. Of these, nine DEGs were up-regulated in the comparison of C7d vs. T7d, such as *IgPSY* (IZ005980), *IgPDS* (IZ009969), *IgZDS* (IZ006629), *IgLCYb* (IZ013964), *IgZEP* (IZ006381), *IgNSY* (IZ013964), *IgDDE* (IZ004535), *IgDEP* (IZ003702), and *IgFH* (IZ011859); *IgCRTISO* (IZ001261), *IgCHYB* (IZ010839) and *IgVDE* (IZ007819) were down-regulated. There were both β- and ε-branches of carotenoid biosynthesis in the comparison of 7d-W vs. 7d-G, and an abundance of β-carotene as well as small amounts of antheraxanthin, zeaxanthin, and Fx. The content of one carotene (ε-carotene), two carotenoid esters (violaxanthin-laurate and violaxanthin-myristate), and four xanthophylls (antheraxanthin, capsanthin, zeaxanthin, and Fx) increased, and the content of zeaxanthin-palmitate decreased with green light induction. Taken together, beta-carotene, antheraxanthin, zeaxanthin, and Fx involved in Fx biosynthesis were found to be accumulated and up-regulated by green light induction, which showed a trend similar to that of *IgPSY, IgLCYb, IgNSY, IgDDE, IgDEP, IgFH, IgMYB98, IgZDS, IgPDS,* and *IgLHCX2* (Figure 6). The results showed that the up-regulation of these genes in the 7d-G group led to enhance the biosynthetic pathway of Fx. Therefore, we hypothesized that green light can enhance Fx and beta-carotene in the carotenoid pathway. Four unigenes (*IgMYB98*, *IgZDS*, *IgPDS*, and *IgLHCX2*) were selected for expression analysis (Figure S27).

**Figure 6.**
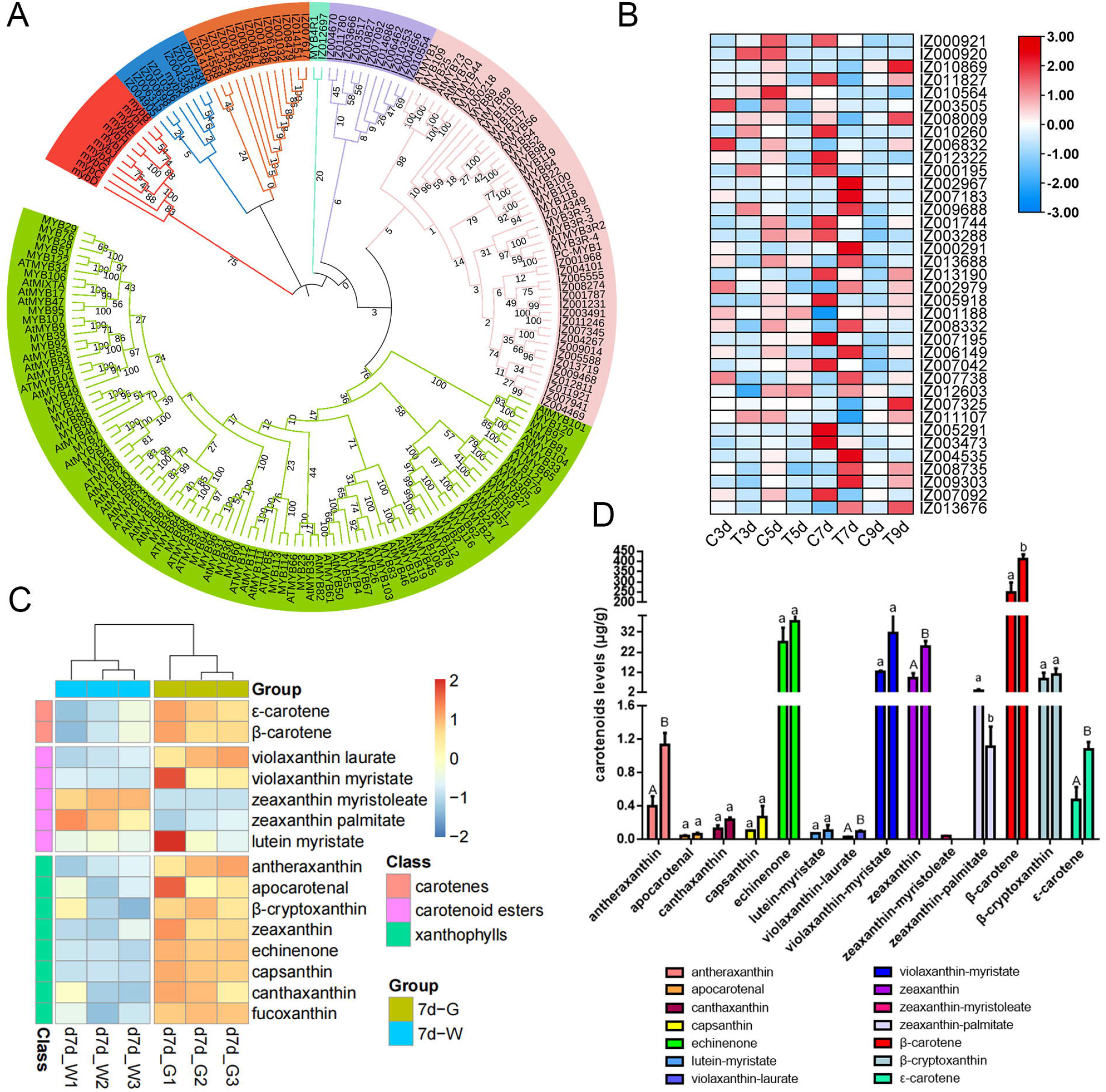
Diagram of the Fx metabolic pathway according to the results of transcriptional regulation and carotenoid changes in *I. galbana*. DEGs are shown in red (up-regulated) and blue (down-regulated). Heat map showing log2 values of transcripts. Chemical reactions possibilities in the Fx pathway are shown in red font. Blue dotted arrows indicate predicted or unknown reactions. Black boxes refer to the content of carotenoid was detected and increased by green light induction at 7 days in the Fx pathway. Red arrows indicate chromatogram of the corresponding carotenoid.

## Discussion

We reported a 92.59 Mb high-quality genome assembly of *I. galbana* with N50 scaffold of 6.99 Mb, and the contig N50 6.99 (Mb) of the assembled genome was 55.04-fold higher than that prior short-read assemblies (N50 contig size of 419 kb) [16]. These results provide the foundation for regulating Fx accumulation in Haptophyta and potential applications for other Fx-producing algae. Phylogenetic and collinearity analysis showed that *E. huxleyi, I. galbana* and *C. tobinii* as monophyletic groups share a common Haptophyta ancestor. Comparative analysis provided evidence of the WGD event, which was dated at ~ 245 Mya and earlier than the divergence time of *E. huxleyi* and *I. galbana* (~ 133 Mya). In *I. galbana,* most genes involved in metabolic regulation have a relatively conserved structure or function, and a few TF gene families have even expanded a subset of duplicates. For example, overexpression of *MYB7* could activate the promoter of the *AdLCY-β* (lycopene-β-cyclase) gene in the carotenoid synthesis pathway of kiwifruit, altering the content of carotenoid and chlorophyll [21]. The MYB gene family (*IgMYB98*) maintained a higher number or expansion in Haptophyta and Bacillariophyta, and exhibited higher transcriptional activity, indicating that the latest algae with rich in carotenoid linked to responses to water stress environmental stimuli exhibit lineage-specific gene expansions in environmental adaptation and metabolic regulation. Notably, lipid metabolism-related genes exhibit significant expansion, including *IgPLMT, IgOAR1,* and Δ-4 desaturase. These results indicated that the expansion of lipid metabolism-related genes in *I. galbana* could enhance the regulation and biosynthesis of Fx.

Although Fx plays an important role as a photosensor of blue-green light and an effector of carotenoid-dependent, the metabolic pathway of Fx remains unclear, and many unknown steps in the process of violaxanthin to Fx [3,22]. We identified one domain from *I. galbana* genome by comparing with the hydroxylase function domain and predicted the *IgFH* gene, which is closely related to the chemical reaction according to the structure of diadinoxanthin and Fx, suggesting that it might function similarly to the characterized diadinoxanthin-fucoxanthin as a candidate gene. The increasing in Fx content between the control and treated groups indicated that 7d with green light would be an important period for Fx biosynthesis. GO enrichment showed that these stage-specific genes in the control and treated groups were related to various biological regulation and fatty acid metabolic processes. Several TFs have been implicated in carotenoid accumulation; however, the members of MYB, bHLH and HB families involved in pigment accumulation and resistance stress showed differential regulation response to the different light qualities [23–25]. We performed WGCNA analysis to identify gene modules. DEGs and TFs were significantly correlated with Fx synthesis. These results suggested that the identified TFs may be related to the accumulation and regulation of Fx production in *I. galbana* by the induction of green light. Transcrptiomics data suggested that transcriptional profiling and phenotypic data methods can be beneficial to identify the most promising candidate genes involved in the Fx biosynthesis.

Studies have showed that multiple key genes are related to the Fx synthesis pathway, including *PYS*, *PDS*, *ZISO*, *ZDS*, *CRTISO*, and *LCYb* [10–11,14]. Comprehensive analysis of multi-omics helps reveal the underlying accumulation of carotenoid [26–29]. For example, Jia et al. revealed the molecular mechanism of white petal color in *Brassica napus* by metabolomic and transcriptomic analyses, mining several candidate genes involved in carotenoid biosynthesis (*BnWRKY22, BnNCED4b*) [27]. Xia et al. found that DEGs and DACs involved in carotenoid biosynthesis were significantly up-regulated and accumulated more in yellow flower petals than in the green bud petals and white flower petals, indicating a predominantly promotion function for color transition in *Lonicera japonica* [29]. Thirteen genes (*PSY1*, *PSY2, PDS1, PDS2, ZDS, CYCB, LCYB1, LCYB2, LCYE, CHYB, LUT1, VDE,* and *ZEP*) were related to the carotenoid biosynthesis, which was strongly correlated with the changes in lycopene, β-carotene, and β-cryptoxanthin, providing an insight into controlling fruit color in papaya fruit [28]. Up- or down-regulated DEGs involved in the carotenoid biosynthetic pathway greatly affect the content of trans-β-carotene trans-β-cryptoxanthin and 5, 8-epoxy-β-carotene, resulting in a striking difference between peel and flesh tissue during on-tree loquat development [26]. Although most studies on the carotenoid biosynthesis have focused on the color transition of fruits and flower petals, combined metabolome and transcriptome analysis of carotenoid biosynthesis in *I. galbana* have not been reported yet. In this study, we identified 12 DEGs and 4 DACs involved in Fx biosynthesis by metabolomic and transcriptomic analyses. Notable increases in carotenoids involved in Fx biosynthesis from the 7d-W to 7d-G samples included ε-carotene, antheraxanthin, zeaxanthin and Fx, suggesting the abundant diversity of carotenoids present under green light. *LCYb* catalyzes the formation of β-carotene and its oxides from lycopene, which is a key step in the synthesis of β-carotene [28]. *ZEP* plays a key function in the xanthophyll cycle of plants, catalyzing the conversion of zeaxanthin to anthraxanthin and violaxanthin [30]. *NXS* catalyzes the conversion of the double-epoxidation precursor violaxanthin into lutein with equilibrated double bonds, representing the classic end of the formation of plant xanthophyll [31]. Therefore, we hypothesized that green light can accumulate β-carotene, zeaxanthin, and Fx by activating the xanthophyll cycle process in the Fx pathway. The results of genome and transcrptiomone indicate that how the genome of *I. galbana* provides a useful model for studying the evolution of Fx-producing algae and the mechanism of Fx biosynthesis.

## Conclusion

In summary, we report a high-quality genome of *I. galbana* LG007 by using the PacBio SEQUEL platform and Hi-C technology. Domain identification of a novel gene that encodes neoxanthin-Fx hydroxylase was analyzed. Fx content could be increased under green light condition, which is a special simulating factor that occurs during the cultivation of *I. galbana*. Metabolome analysis indicated that 7d-G accumulated a higher content of carotenoids than that of 7d-W, and β-carotene was the main carotenoids, accounting for 79.09% of the total carotenoids. Multi-omics analysis revealed that DEGs or TFs significantly correlated with the accumulation and regulation of Fx synthesis, including *IgMYB98*, *IgZDS*, *IgPDS*, and *IgLHCX2*. Therefore, our findings advance the understanding of Fx biosynthesis and its regulation, providing an important resource for food and pharmaceutical applications.

## Materials and methods

### Sample materials and genome sequencing

*I. galbana* LG007 was separated from the near sea area of Chuanshi Island in Fujian and deposited with the Southern Institute of Oceanography, Fujian Normal University, China. The seawater used for culture was collected from the near sea area of Chuanshi Island in Fujian, with a salinity of 28 ‰. The algae were cultured in 100 mL f/2 medium and incubated at 23 ± 1 °C with shaking the bottle manually 6 times per day under continuous light of 100 μmol photons m ^-2^ s ^-1^ with fluorescent lamps [32]. Genomic DNA was prepared using the CTAB method to construct the Pacbio and Illumina libraries. Concentrated DNA were applied to select size with BluePippin; they were repaired, tailed, adaptor-ligated, and used for library construction in accordance with the released protocol released by PacBio. Next, ~ 15.5 Gb of clean data were obtained from the PacBio sequencing, and used to estimated genome size.

### Genomic size estimation

The BD FACSCalibur cytometer (Becton Dickinson, San Jose, CA) was used to estimate the genome size of *I. galbana* LG007, which was calculated as a ratio of the average fluorescence. We further used Illumina short reads and a K-mer-based method to estimate the genome size, heterozygosity, and repeat content of *I. galbana* LG007. Approximately 8.92 Gb of Illumina data were generated and used to calculate the abundance of 17-K-mers by Genomescop2 software (version 2.0) (Table S1).

### Genome assembly and completeness assessment

After removing the low-quality short reads and sequencing adaptors, the clean data were corrected, trimmed and assembled by using CANU software with default parameters [33]. For improving the accuracy of base-pair correction, preliminary assembled contigs were polished by the BWA and Pilon software using ~8.92 Gb Illumina data [34]. Summary statistics of the assembled genome are presented in the Table S1. Assessment of the completeness of *I. galbana* LG007 genome was evaluated through BUSCO using eukaryotic models [35]. Illumina short reads and PacBio long reads were properly mapped to the genome via Bowtie2 [36] and Minimap2 [37], respectively.

### Superscaffold construction using Hi-C technology

The nuclear integrity of samples were examined by 6-diamidino-2-phenylindole (DAPI) staining to guarantee the quality of the Hi-C procedure [38,39]. By filtering adapter sequences and low-quality pair-end reads, ~ 12.35 Gb of clean data were generated (Table S1). The Hi-C clean data were properly mapped to the *I. galbana* LG007 by BWA (version 0.7), and then erroneous mappings and duplicates were filtered by the Juicer pipeline [40]. The output of the Juicer pipeline was used for 3D-DNA analysis with default parameters, including misjoin correction, ordering, and orientation [41]. To ensure the accuracy of assembly, the assembled contigs combined with Hi-C data were ordered and clustered into the superscaffolds by using LACHESIS based on the relationships among valid reads [39], then filtered the invalid read pairs by HiC-Pro (version 2.7.8) [42].

### Gene and repetitive sequence annotation

LTR_FINDER, Tandem Repeats Finder, and RepeatMasker were used to identify the repeat sequences in the *I. galbana* LG007 genome, as previously described [43,44]. We then performed annotation of the *I. galbana* LG007 genome assembly by combining the homology search results, *de novo* prediction and transcriptome evidence. *E. huxleyi*, *C. tobinii*, *P. tricornutum*, *E. siliculosus* and *C. reinhardtii* were selected to perform the homology annotation. We predicted the coding genes with MAKER pipeline (version 2.31.9) by using transcript sequences from RNA-Seq [45]. The protein-coding genes were compared to the content of eggNOG, GO, COG, and KEGG by using BLASTP with an E-value cutoff of 1E^−^5 [46–48]. ncRNAs and small RNAs were identified by searching from the Rfam and miRNA databases, respectively [49]. In addition, other types of non-coding RNA, including miRNA and snRNA, were predicted by alignment to the Pfam database using INFERNAL software.

### Genome evolution analysis

Single-copy genes were identified among 15 genomes by using OrthoFinder and downloaded from the NCBI database, including *E. huxleyi*, *C. tobinii*, *P. tricornutum*, *E. siliculosus*, *C. reinhardtii*, *Pennisetum purpureum*, *Chloropicon primus*, *Bathycoccus prasinos*, *Porphyra umbilicalis*, *Gracilariopsis chorda, Cyanidioschyzon merolae, R. subcapitata, T. pseudonana,* and *T. oceanica* [50]. Based on the identified single-copy protein sequences, a phylogenetic tree was constructed by using RAxML software with *P. purpureum, P. umbilicalis, G. chorda* and *C. merolae* as the outgroup [50]. The divergence time of each tree node was calculated using the TimeTree database and the MCMCtree software. We used CAFÉ software (version 3.1) to identify the expansion and contraction of gene families with the criterion of a *p-value* < 0.05 [51]. GO terms for gene were obtained from the corresponding InterPro or Pfam entries. KEGG terms were assigned at the KEGG pathway database (http://www.genome.jp/kegg). Enrichment analyses of KEGG pathway and GO term were performed using the OmicShare tools (https://www.omicshare.com/tools).

### Identification of candidate gene related to Fx pathway in *I. galbana*

The identification of orthologs of previously known functional genes in the Fx pathway were performed by combining the results of the genome assembly annotations, transcriptional expression level and conserved domain BLAST searches. The orthologs and previously known functional genes in the Fx pathway exhibited identity scores >85%, suggesting that these genes are functionally similar and can be used as candidate genes involved in Fx biosynthesis. For gene family analysis related to Fx pathway in *I. galbana,* BLASTP and HMMER were used to search for homologous proteins of related gene families in *I. galbana* LG007 (E-value < 1E^−^10), and then further confirmed using both the NCBI conserved domain database tool [52]. The final deduced homologous proteins sequence were aligned by using the ClustalW software [53]. RAxML software was used to construct a phylogenetic tree via the maximum likelihood method with 1000 bootstrap iterations [54].

### Analysis of WGD and gene synteny

For detecting the polyploidization events in the *I. galbana* LG007genome, the protein sequences from *I. galbana* LG007 were intercompared to identify conserved paralogs by using BLASTP with an E-value ≤ 1E^−^5. *E. huxleyi, C. tobinii, P. tricornutum,* and *C. reinhardtii* were also analyzed and used for comparison. We identified the collinear blocks by using MCScanX and calculated the non-synonymous (*K*a), *K*s, and *K*a/*K*s values for syntenic gene pairs by using *K*a*K*s_Calculator software (version 2.0) [55,56]. Syntenic blocks between *I. galbana* LG007, *E. huxleyi* and *C. tobinii* were identified by using MCScanX [55].

### Transcriptome sequencing

Our previous results suggested that the green light could promote Fx synthesis in the 7d stage (*p* < 0.05, 14.06% higher) (Figure S2). To investigate the transcriptome dynamics and response of Fx accumulation under different light qualities in *I.galbana* LG007, we performed transcriptomic analysis of the simulated cells under white and green light at different stages of cultivation. *I. galbana* LG007 was cultured in 100 mL f/2 medium and incubated at 23 ± 1 °C with shaking the bottle manually 4-6 times per day under continuous light of 100 μmol photons m ^-2^ s ^-1^ with a 12 h:12 h light:dark cycle [33]. The culture (10^6^ cells/mL) was evenly divided into eight groups and cultured in a spectrum-adjustable plant growth box (Catalog No. AKF-KYG04-600DZ, Anhui Ancorgreen Photoelectric Technology Co. Ltd., Hefei, China) at 3d, 5d, 7d, and 9 d, respectively (Figure S16A). Four treated groups (T3d, T5d, T7d, and T9d) were treated with green light irradiation 100 μmol photons m ^-2^ s ^-1^ (green light source: LED circular lamp beads [Catalog No. SZG05A0A, Seoul Semiconductor Co. Ltd., Siheung-si, Korea]; spectrum: 525 nm), and four control groups (C3d, C5d, C7d, and C9d) with a white light of 100 μmol photons m ^-2^ s^-1^ (white light source: LED circular lamp beads [Catalog No. LH351H-D, Samsung LED Co. Ltd., Tianjin, China]; spectral range: 400-700 nm) were used as controls. Total RNA from each sample was extracted using a TransZol Up Plus RNA Kit (Catalog No. ER501-01, Transgen Biotech, Beijing, China), and the corresponding cDNA library was constructed for RNA sequencing.

### Gene expression analysis

Approximately 307.77 Gb of high-quality transcript data were produced and processed by Trimmomatic (version 0.36). The high-quality filtered reads were mapped onto the genome by using HISAT2 with the default parameters [57,58]. FPKM values were calculated using Stringtie and Ballgown [59,60]. DEGs between the control and treated groups were analyzed using DESeq2 based on a criteria of a fold change ≥ 1 and false discovery rate ≤ 0.05 [61], then performed by KEGG and GO enrichment analysis, respectively. The ratio of each sample to the genome is more than 90%, and the number of reads per sample was estimated to range from 32,792,958 to 59,111,858 (Tables S22-S23).

### Metabolite profiling and statistical analysis

To explore the metabolites of *I. galbana* LG007 under white and green light, we collected 7d-W and 7d-G samples with three biological replicates. Stock solution of Fx was prepared by dissolving 0.5 mg Fx in 50 mL methanol solution. Stock solutions of Fx standard was gradient diluted as follows: 5 μg/mL, 10 μg/mL, 20 μg/mL, 50 μg/mL, and 100 μg/mL. Fx production was detected according to the linear relationship between the peak areas of the samples and a standard curve (R^2^ = 0.999). According to the above method of *I.galbana* LG007 fermentation, 10 mL of the fermentation mixture was centrifuged at 8,000 rpm for 20 min, followed by removal of the supernatant and washing with distilled water for three times. After vacuum freeze-drying, 1 mL of acetone was added to the freeze-dried algae to extract the total carotenoids. The supernatant was harvested by centrifugation (8,000 rpm, 15 min) and filtered (0.25 μm filter membrane), respectively. The supernatant was analyzed by HPLC using a Waters e2695 Liquid Chromatograph equipped with a Waters 2998 PDA detector and separated on a SunFire C18 HPLC column (250×4.6mm; 5μm). The mobile phase consisting of a ternary solvents of water (A)/methanol (B)/acetonitrile (C) (15:30:55, v/v/v) and the flow rate of the mobile phase was 1 mL/min. The Fx content was detected using a PDA detector at 447 nm. Except for Fx, other carotenoids are obtained using MetWare (http://www.metware.cn/) according to the following method: (1) the vacuum freeze-dried algae were crushed using a grinding mill (MM 400, Retsch) at 30 Hz for 1.5 min. Powder (100 mg) was dissolved in 1.2 mL of 70% methanol, vortexed for 30 s, and stored at 4°C overnight. (2) the mixture was centrifuged at 12,000 rpm for 10 min, and then filtered through a 0.22 μm membrane to obtain the supernatant for subsuquent analysis [62]. (3) carotenoid content was detected using a UPLC system (Shim-pack UFLC SHIMADZU CBM30A system) and an MS/MS system (AB Sciex 6500 QTRAP), which was equipped with a APCI + and controlled by Analyst (version 1.6.3) software. DACs were determined by a fold change ≥ 1 and *p-value* < 0.05. Identified metabolites were annotated and mapped by using the KEGG compound and pathway databases, respectively.

### Co-expression network analysis

Based on log2 (1 + FPKM) values, WGCNA was performed by using a minimum module size of 30; a soft power of 11 (control group), 8 (treated group), and 14 (control-treated group); and a merge cut height of 0.25 (Figure S28). Eigengene values of WGCNA module were calculated and associated with lipid and Fx content at different culture stages [63]. Each module gene was analyzed by GO enrichment and visualized using Cytoscape [64].

### qRT-PCR validation and *in vitro* experiments

Quantitative real-time PCR (qRT-PCR) experiment was performed using SYBR Green PCR Master Mix (TaKaRa, China) in an Applied Biosystems 7300 real-time PCR System (Framingham, MA, USA) [65]. The *IgHF* gene was amplified from *I. galbana* LG007, and ligated into the pTrc99a vector (Figure S29). After transformation into *E. coli* K-12 MG1655 cells (Invitrogen, Carlsbad, CA), recombinant protein expression was induced by 0.2 mM isopropyl-thio-β-D-galactopyranoside with vigorous shaking 220 rpm for 24 h at 37 °C. 30 OD cells were harvested by centrifugation at 8,000 rpm for 10 min, then induced with 10 mL Tris-HCl lysis buffer (50 mM, pH 7.5), 10% (v/v) glycerol, and 1.67 μM neoxanthin (Sigma-Aldrich, Louis, MO) with vigorous shaking at 220 rpm at 37 °C for 12 h, respectively. 2 mL cultures were centrifuged at 8,000 rpm for 5 min, and suspended in 2 mL methanol (chromatographic grade) by ultrasonic crushing at 60 Hz for 20 min. The supernatant was obtained by centrifugation (8,000 rpm, 5 min) and filtration (0.22 μm filter membrane) and further used for HPLC analysis.

### Data availability

The assembled genome sequences have been deposited at the National Center for Biotechnology Information (NCBI) (BioProject: PRJNA669236), and are publicly accessible at https://www.ncbi.nlm.nih.gov/bioproject. Raw sequencing data for RNA-Seq were used for annotation and biological analyses and have been deposited in the Genome Sequence Archive in National Genomics Data Center, China National Center for Bioinformation/Beijing Institute of Genomics, Chinese Academy of Sciences (GSA: CRA003291), and are publicly accessible at https://ngdc.cncb.ac.cn/gsa [66]. The whole genome sequence data reported in this paper have been deposited in the Genome Warehouse in National Genomics Data Center, Beijing Institute of Genomics, Chinese Academy of Sciences (GWH: GWHAZHV00000000), and are publicly accessible at https://bigd.big.ac.cn/gwh [67].

## CRediT author statement

**Duo Chen:** Methodology, Formal analysis, Writing-Original Draft. **Xue Yuan:** Formal analysis, Data Curation. **XueHai Zheng:** Methodology, Validation, Data Curation. **Jingping Fang:** Formal analysis, Resources, Data Curation. **Gang Lin:** Resources. **Rongmao Li:** Resources. **Jiannan Chen:** Resources. **Wenjin He:** Investigation. **Zhen Huang:** Investigation. **Wenfang Fan:** Formal analysis. **Limin Liang:** Validation, Data Curation. **Chentao Lin:** Resources. **Jinmao Zhu:** Writing - review & editing. **Youqiang Chen:** Writing - review & editing. **Ting Xue:** Conceptualization, Writing-Review & Editing, Supervision, Project administration, Funding acquisition. All authors read and approved the final manuscript.

## Acknowledgments

We thank Feng Qi (Fujian Normal University) for sharing plasmid and method that were used for the the catalytic reaction of *IgFH* gene in *E. coli*. This work was supported by the National Natural Science Foundation of China (Grant no. 42006087), and the Sugar Crop Research System, China (Grant no. CARS-170501).

## Supplementary materials

**Figure S1 Determination the content of Fx in different strains or species by HPLC**

All experiments were performed in triplicate. Each value presents the mean ± SD. “*I*” represents error bars for the various determinations (n = 3). *I. galbana* GY-D66 and *I. zhangjiangensis* GY-H2 were purchased from Shanghai Guangyu Biological Technology Co., Ltd. *I. galbana* FACHB-861 and *P. tricornutum* FACHB-863 were purchased from Freshwater Algae Culture Collection at the Institute of Hydrobiology. *I. zhangjiangensis* YB1Z2 is a mutant strain of *I. zhangjiangensis* GY-H2 mutagenized by atmospheric and room temperature plasmas. *I. galbana* LG007, *Thalassiosira weissflogii* ND-8, and *Chaetoceros muelleri* LJ1 were isolated and identified by the Southern Institute of Oceanography, Fujian Normal University, China.

**Figure S2 Determination the Fx content of *I. galbana* by HPLC under white and green light**

**A.** Ordinary algae culture chamber. **B.** Content of fucoxanthin in 10^7^ cell density at different cultivation times with green light irradiation 100 μmol photons m ^-2^ s^-1^ and white light of 100 μmol photons m ^-2^ s ^-1^, respectively. All experiments were performed in triplicate. Each value presents the mean ± SD. “*I*” represents error bars for the various determinations (n = 3). “**” and “*” indicate that significance are at *0.01* and *0.05* level, respectively.

**Figure S3 GC content and sequencing depth correlation analysis**

The abscissa represents GC content, the ordinate represents sequencing depth, the right is the sequencing depth distribution, and the top is the GC content distribution.

**Figure S4 The size estimation of the *I. galbana* genome by flow cytometry**

The genome size of *I. galbana* was estimated to be approximately 83.75 ± 0.107% of *C. reinhardtii* (~ 120 Mb) as internal reference. We validated the result by flow cytometry, with the *I. galbana* genome size identified as100.5 ± 12.86 Mb.

**Figure S5 17-K-mer count distribution for the *I. galbana* genome size estimation**

**Figure S6 *I. galbana* genome-wide all-by-all Hi-C interaction heat map**

The map shows high-resolution individual superscaffolds, which were scaffolded and assembled independently.

**Figure S7 Syntenic analysis of *I. galbana* superscaffolds**

**A.** line. **B.** bar. **C.** Syntenic analysis of *I. galbana* superscaffolds and *E. huxleyi* scaffolds.

**Figure S8 Figure S8 Structure of hydroxylase gene family in *I. galbana***

**Figure S9 Analysis of the activity of *IgFH in vitro***

**Figure S10 The evolutionary tree of epoxidase gene families in *I. galbana* (IZ), *A. thaliana* (NP), *P. tricornutum* (XP) and *C. reinhardtii* (ChrXP)**

**Figure S11 Motif composition of whole amino acid sequences for epoxidase gene family in *I. galbana***

**Figure S12 Structure of epoxidase gene family in *I. galbana***

**Figure S13 The evolutionary tree of de-epoxidase gene families in *I. galbana* (IZ), *A. thaliana* (NP), *P. tricornutum* (XP) and *C. reinhardtii* (ChrXP)**

**Figure S14 Motif composition of whole amino acid sequences for de-epoxidase gene family in *I. galbana***

**Figure S15 Structure of de-epoxidase gene family in *I. galbana***

**Figure S16 Heat map showing the control- and treated-specific genes (FPKM≥8000)**

Each box represents an individual gene, and the red and blue boxes represent relatively high levels and low levels of gene expression, respectively.

**Figure S17 Determination the Fx content of *I. galbana* by HPLC under white and green light**

**A.** Spectrum adjustable plant growth box. **B.** Content of fucoxanthin at different cultivation times with green light irradiation 100 μmol photons m ^-2^ s ^-1^ and white light of 100 μmol photons m ^-2^ s ^-1^, respectively. All experiments were performed in triplicate. Each value presents the mean ± SD. “*I*” represents error bars for the various determinations (n = 3). “**” indicate that significance is at *0.01* level.

**Figure S18 Bar graph showing the number of preferentially expressed genes specifically and commonly in control and treated groups at different cultivation times**

**Figure S19 GO enrichment map of preferentially expressed genes at all the stages in in control and treated groups**

**Figure S20 Motif composition of whole amino acid sequences for MYB gene family in *I. galbana***

**Figure S21 Distribution of number of genes in different modules**

**A.** Number of genes in different modules in control group. **B.** Number of genes in different modules in treated group.

**Figure S22 GO enrichment of the specific expressed genes in the red module of *I. galbana***

**Figure S23 KEGG enrichment of the specific expressed genes in the red module of *I. galbana***

**Figure S24 GO enrichment of the specific expressed genes in the turquoise module of *I. galbana***

**Figure S25 KEGG enrichment of the specific expressed genes in turquoise module of *I. galbana***

**Figure S26 Co-expression network analysis**

**A.** Cluster dendrogram and module assignment from WGCNA in control and treated groups. Each color represents a certain gene module. **B.** Heat map of the correlation of WGCNA modules with traits. Red is the correlation between module and trait, and blue is a negative correlation. The module highlighted with dark color represents a significant module associated with traits. **C.** The heat map for modules with Fx. **D.** GO enrichment of the specific expressed genes in the midnightblue module of *I. galbana*.

**Figure S27** Expression analysis for qRT-PCR validation according to the RNA-seq data

**A.** *IgLHCX2*. **B.** *IgZDS*. **C.** *IgPDS*. **D.** *IgMYB98*.

**Figure S28 A soft power for WGCNA analysis**

**A.** A soft of control group (11). **B.** A soft of treated group (8). **C.** A soft of control-treated group (14).

**Figure S29 Construction of the pTrc99a-IgFH over-expression vector**

**Table S1 Sequencing data used for *I. galbana* LG007 genome construction**

**Table S2 Assembly statistics for nuclear genome**

**Table S3 Assessment of the completeness of the *I. galbana* LG007 genome assembly by BUSCO**

**Table S4 Genomic resequencing alignment rate and coverage assessment**

**Table S5 Pacbio data alignment rate**

**Table S6 Pseudomolecule length statistics after Hi-C assisted assembly**

**Table S7 Quality assembly statistics of the *I. galbana* after the Hi-C data based pseudo-chromosome assembly**

**Table S8 Statistics for gene, exon, CDS and introne in *I. galbana* LG007 genome**

**Table S9 Annotation statistics for the *I. galbana* LG007 genome**

**Table S10 Transcription factors**

**Table S11 Statistical analysis of non-coding RNAs in *I. galbana* LG007 genome**

**Table S12 Repetitive element annotations in the *I. galbana* LG007**

**Table S13 KEGG enrichment of the expanded families genes identified in *I. galbana***

**Table S14 GO enrichment of the expanded families genes identified in *I. galbana***

**Table S15 KEGG enrichment of the contracted families genes identified in *I. galbana***

**Table S16 GO enrichment of the contracted families genes identified in *I. galbana***

**Table S17 GO enrichment of the specific families genes identified in *I. galbana***

**Table S18 KEGG enrichment of the specific families genes identified in *I. galbana***

**Table S19 Differentially expressed genes in the comparison of C7d vs. T7d by transcriptome data**

**Table S20 Raw data of carotenoids compound content detected by HPLC**

**Table S21 Differentially accumulated carotenoids compounds in the comparison of 7d-W vs. 7d-G by targeted metabolomics (n=3)**

**Table S22 Statistical analysis of transcriptome data**

**Table S23 Mapping data of each transcriptome sample to the generated genome assembly**

