## Supplementary for "Multi-omics Analyses Provide Insight into the Biosynthesis Pathways of Fucoxanthin in *Isochrysis galbana*": editing certification of previous manuscript.pdf

AJE®

### Editing Certificate

This document certifies that the manuscript

prepared by the authors

**Ting Xue**

was edited for proper English language, grammar, punctuation, spelling, and overall style by one or more of the highly qualified native English speaking editors at AJE.

This certificate was issued on **October 28, 2020** and may be verified on the AJE website using the verification code **3896-1A98-B9CC-40C4-D768**.

AJE® EDITING CERTIFIED

Neither the research content nor the authors' intentions were altered in any way during the editing process. Documents receiving this certification should be English-ready for publication; however, the author has the ability to accept or reject our suggestions and changes. To verify the final AJE edited version, please visit our verification page at aje.com/certificate. If you have any questions or concerns about this edited document, please contact AJE at.

AJE provides a range of editing, translation, and manuscript services for researchers and publishers around the world. For more information about our company, services, and partner discounts, please visit aje.com.
