## Supplementary for "Multi-omics Analyses Provide Insight into the Biosynthesis Pathways of Fucoxanthin in *Isochrysis galbana*": editing certification of revised version.pdf

ELSEVIER

### Certificate of Elsevier Language Editing Services

The following article was edited by Elsevier Language Editing Services:

Authored by:

**Ting Xue**

Date: 22-Nov-2021

Serial number: LEEX-16197-4A8C1B0C7CF0
