## Supplementary for "Multi-omics Analyses Provide Insight into the Biosynthesis Pathways of Fucoxanthin in *Isochrysis galbana*": Supplementary tables S1-12,19-23.docx

**Table S1** Sequencing data used for *I. galbana* LG007 genome construction

| Library  resource | Sequencing platform | Insert size (bp) | Clean data (Gb) | Sequence coverage (X) | Use of the data |
| --- | --- | --- | --- | --- | --- |
| Genome | Illumina HiSeq X Ten | 250 bp | 8.92 | 96 | Genome estimation and polishing |
| Genome | PacBio SEQUEL | 20 kb | 15.53 | 166 | Genome assembly |
| Hi-C | Illumina HiSeq X Ten | 250 bp | 12.35 | 137 | Chromosome construction |
| Transcriptome | Illumina HiSeq X Ten | 0.6-3 kb | 307.77 | - | Difference analysis and annotation |

**Table S2** Assembly statistics for nuclear genome

| Items | Statistics |
| --- | --- |
| Estimated genome size | 92.59 Mb |
| Genomic G+C content | 58.44% |
| Number of assembled scaffolds | 353 |
| Number of scaffolds > 2 kb | 353 |
| min length (bp) | 5,065 |
| max length (bp) | 2,925,745 |
| Average length (bp) | 262,298 |
| Scaffolds N50 | 666.7 kb |
| Predicted gene models | 14,900 |
| Average coding sequence length (bp) | 1,428 |
| Average transcript length (bp) | 1,764 |
| Average gene length (bp) | 1,789 |
| Chromosomal assembled N50 | 6.99 Mb |

**Table S3** Assessment of the completeness of the *I. galbana* LG007 genome assembly by BUSCO

| Type | Number | Percent (%) |
| --- | --- | --- |
| Complete BUSCOs (C) | 254 | 83.8 |
| Complete and single-copy BUSCOs (S) | 248 | 81.8 |
| Complete and duplicated BUSCOs (D) | 6 | 2.0 |
| Fragmented BUSCOs (F) | 13 | 4.3 |
| Missing BUSCOs (M) | 36 | 11.9 |
| Total BUSCO groups searched | 303 | 100 |

**Table S4** Genomic resequencing alignment rate and coverage assessment

| Items | Statistics |
| --- | --- |
| Unmapped bases (Mb) | 144.6 |
| Mapped bases (Mb) | 887.5 |
| Map rate (%) | 98.4 |
| Genome Length (Mb) | 93 |
| Mean Depth | 89 |
| Coverage Rate (%) | 97.6 |

**Table S5** Pacbio data alignment rate

| Items | Statistics |
| --- | --- |
| QC-passed reads | 288211 |
| Duplicates | 0 |
| Mapped reads | 287594 |
| Mapping rate (%) | 99.78 |

**Table S6** Pseudomolecule length statistics after Hi-C assisted assembly

| Pseudomolecule | Length/bp |
| --- | --- |
| Chr1 | 11908665 |
| Chr2 | 10401026 |
| Chr3 | 7919487 |
| Chr4 | 7664558 |
| Chr5 | 7036706 |
| Chr6 | 6994209 |
| Chr7 | 6185998 |
| Chr8 | 5984543 |
| Chr9 | 5214086 |
| Chr10 | 4726453 |
| Chr11 | 4149466 |
| Chr12 | 3870612 |
| Chr13 | 3512008 |
| Chr14 | 3046428 |
| Chr15 | 2341788 |
| Total anchored | 90956033 |
| Unanchored | 1777877 |

**Table S7** Quality assembly statistics of the *I. galbana* after the Hi-C data based pseudo-chromosome assembly

| Items | Canu | | Hi-C | |
| --- | --- | --- | --- | --- |
|  | Contig_len (Mb) | Contig_number | Contig_len (Mb) | Contig_number |
| Total | 92.59 | 353 | 92.64 | 142 |
| Max | 2.92 | - | 11.91 | - |
| Number>=2 kb | - | 353 | - | 78 |
| N50 | 0.66 | - | 6.99 | - |

**Table S8** Statistics for gene, exon, CDS and introne in *I. galbana* LG007 genome

|  | Gene stat | Exons stat | CDS stat | Introne stat |
| --- | --- | --- | --- | --- |
| Total length | 26,666,990 | 21,283,343 | 21,283,343 | 5,383,647 |
| Total number | 14,900 | 43,481 | 14,900 | 28,581 |
| Average length | 1,789.73 | 489.49 | 1,428.41 | 188.36 |

**Table S9** Annotation statistics for the *I. galbana* LG007 genome

| **Annotation statistics for nuclear genome** | | Number | Percent (%) |
| --- | --- | --- | --- |
|  | Total protein | 14,900 |  |
|  | NR | 12,469 | 83.68 |
|  | eggNOG | 9,161 | 61.48 |
|  | GO | 4,977 | 33.40 |
|  | COG | 9,161 | 61.48 |
|  | KEGG | 3,773 | 25.32 |
|  | At least in one database | 12,500 | 83.89 |

**Table S10** Transcription factors

| Number | Classify |
| --- | --- |
| 198 | protein kinase family protein |
| 55 | heat shock protein |
| 49 | WD-40 repeat family protein / zfwd4 protein (ZFWD4) |
| 44 | myb domain protein 3r-3 |
| 23 | pentatricopeptide (PPR) repeat-containing protein |
| 13 | Calmodulin-binding transcription activator protein with CG-1 and Ankyrin domains |
| 11 | CCCH-type zinc finger protein with ARM repeat domain |
| 7 | DnaJ domain ;Myb-like DNA-binding domain |
| 7 | Zinc finger (C2H2 type) family protein / transcription factor jumonji (jmj) family protein |
| 6 | Homeodomain-like superfamily protein |
| 6 | ethylene induced calmodulin binding protein |
| 6 | pathogenesis related homeodomain protein A |
| 3 | GATA transcription factor 15 |
| 3 | E2F transcription factor |
| 2 | winged-helix DNA-binding transcription factor family protein |
| 2 | Plant-specific transcription factor YABBY family protein |
| 1 | AP2/B3-like transcriptional factor family protein |
| 1 | HD-ZIP IV family of homeobox-leucine zipper protein with lipid-binding START domain |
| 1 | Homeobox-leucine zipper protein 4 (HB-4) / HD-ZIP protein |
| 1 | K-box region and MADS-box transcription factor family protein |

**Table S11** Statistical analysis of non-coding RNAs in *I. galbana* LG007 genome

| ncRNA Items | Number |
| --- | --- |
| tRNA | 95 |
| rRNA | 58 |
| snRNA | 4 |

**Table S12** Repetitive element annotations in the *I. galbana* LG007

|  | No. of TEs | Length (bp) | % of TEs | % of genome |
| --- | --- | --- | --- | --- |
| Total repeat fraction | 161907 | 43349564 | 100 | 46.82 |
| Class I: Retroelement | 73047 | 31195044 | 71.96 | 33.69 |
| LTR Retrotransposon | 27946 | 14219065 | 32.8 | 15.36 |
| Ty1/Copia | 1992 | 998417 | 2.3 | 1.08 |
| Ty3/Gypsy | 4491 | 4191464 | 9.67 | 4.53 |
| Other | 21463 | 9029184 | 20.83 | 9.75 |
| non-LTR Retrotransposon | 26756 | 11607249 | 26.78 | 12.54 |
| LINE | 21408 | 10762090 | 24.83 | 11.62 |
| SINE | 5348 | 845159 | 1.95 | 0.91 |
| unclassified retroelement | 18345 | 5368730 | 12.38 | 5.8 |
| Class II: DNA Transposon | 33472 | 7828746 | 18.06 | 8.46 |
| CMC | 1176 | 203681 | 0.47 | 0.22 |
| hAT | 4039 | 1494270 | 3.45 | 1.61 |
| Mutator | 629 | 121730 | 0.28 | 0.13 |
| Tc1/Mariner | 180 | 71173 | 0.16 | 0.08 |
| PIF/Harbinger | 242 | 98683 | 0.23 | 0.11 |
| Other | 27026 | 5768036 | 13.31 | 6.23 |
| Helitron | 69 | 12880 | 0.03 | 0.01 |
| Tandem Repeats | 43633 | 3981448 | 9.18 | 4.3 |
| Unkown | 2891 | 1025862 | 2.37 | 1.11 |

**Table S19** Differentially expressed genes in the comparison of C7d vs. T7d by transcriptome data

| gene_id | log2FoldChange | lfcSE | stat | pvalue | padj |
| --- | --- | --- | --- | --- | --- |
| IZ000921 | -0.618228045 | 0.28330285 | -2.182216116 | 0.029093586 | 0.128349004 |
| IZ000920 | -0.862878253 | 0.395637297 | -2.180983083 | 0.029184668 | 0.128651816 |
| IZ010869 | -0.019082564 | 0.513613945 | -0.037153517 | 0.970362601 | 0.987349924 |
| IZ011827 | -0.011840673 | 0.34486052 | -0.034334672 | 0.972610277 | 0.98799762 |
| IZ010564 | -0.772164862 | 0.382123114 | -2.02072273 | 0.043308475 | 0.165837741 |
| IZ003505 | -0.300845842 | 0.657648064 | -0.4574572 | 0.647342459 | 0.817137655 |
| IZ008009 | -0.46717575 | 0.49530133 | -0.943215215 | 0.345570833 | 0.581678454 |
| IZ010260 | -0.654490375 | 0.30151432 | -2.170677585 | 0.029955553 | 0.130988806 |
| IZ006832 | 0.358276546 | 0.41590108 | 0.861446541 | 0.388992153 | 0.620149672 |
| IZ012322 | -1.041756606 | 0.374929781 | -2.7785379 | 0.005460414 | 0.042540685 |
| IZ000195 | -1.812689526 | 1.357545286 | -1.335270023 | 0.181787986 | 0.402025948 |
| IZ002967 | 0.095797365 | 0.520894459 | 0.183909357 | 0.854084564 | 0.935664555 |
| IZ007183 | 0.129312115 | 0.50400017 | 0.256571569 | 0.797509525 | 0.907113004 |
| IZ009688 | 1.78319659 | 0.298305616 | 5.97775065 | 2.26E-09 | 3.38E-07 |
| IZ001744 | -1.692288335 | 0.47018672 | -3.599183607 | 0.000319218 | 0.005373199 |
| IZ003288 | -0.726128197 | 0.452481446 | -1.604768998 | 0.108544651 | 0.294645988 |
| IZ000291 | 0.254309903 | 0.288428783 | 0.881707784 | 0.377934851 | 0.610612329 |
| IZ013688 | 1.393393923 | 0.378958137 | 3.67690725 | 0.000236079 | 0.004268348 |
| IZ013190 | -0.215916559 | 0.283158015 | -0.762530274 | 0.445743582 | 0.667805133 |
| IZ002979 | 2.110430225 | 0.414719263 | 5.088816496 | 3.60E-07 | 2.74E-05 |
| IZ005918 | -2.151611413 | 0.423190094 | -5.084266964 | 3.69E-07 | 2.76E-05 |
| IZ001188 | 0.163039038 | 0.373436404 | 0.436591174 | 0.662407864 | 0.827300281 |
| IZ008332 | 1.281168966 | 0.461675097 | 2.775044558 | 0.005519415 | 0.0427694 |
| IZ007195 | -1.15478264 | 0.575781124 | -2.005593084 | 0.04489968 | 0.169885644 |
| IZ006149 | 1.375698459 | 0.302553454 | 4.546960017 | 5.44E-06 | 0.000239136 |
| IZ007042 | -0.637016402 | 0.29061089 | -2.191990816 | 0.02838017 | 0.126337578 |
| IZ007738 | 0.905071091 | 0.322767895 | 2.804092678 | 0.00504584 | 0.040526097 |
| IZ012603 | 1.494212381 | 0.284789023 | 5.246734463 | 1.55E-07 | 1.38E-05 |
| IZ007325 | -0.076755196 | 0.329380162 | -0.233029202 | 0.81573873 | 0.916852587 |
| IZ011107 | -0.545539022 | 0.307710975 | -1.772894262 | 0.07624623 | 0.237586114 |
| IZ005291 | -0.3532401 | 0.314010222 | -1.124931851 | 0.260617914 | 0.493797145 |
| IZ003473 | -0.090417255 | 0.376020193 | -0.240458509 | 0.809974825 | 0.913979253 |
| IZ004535 | 1.205586609 | 0.407258941 | 2.960245895 | 0.003073936 | 0.028450653 |
| IZ008735 | 0.616690163 | 0.304361504 | 2.026176621 | 0.042746689 | 0.16426549 |
| IZ009303 | 0.99199501 | 0.317468966 | 3.124699155 | 0.00177987 | 0.019397025 |
| IZ007092 | -1.657513516 | 0.276271983 | -5.999571508 | 1.99E-09 | 1.32E-07 |
| IZ013676 | 1.152823461 | 0.319632713 | 3.606713001 | 0.0003101 | 0.005271705 |

**Table S20 Raw data of carotenoids compound content detected by HPLC**

| Compound | 7d-W1 | 7d-W2 | 7d-W3 | 7d-G1 | 7d-G2 | 7d-G3 | Pvalue | FoldChange | Log2FC |
| --- | --- | --- | --- | --- | --- | --- | --- | --- | --- |
| β-Carotene | 202.74 | 251.97 | 296.51 | 439.63 | 393.78 | 397.05 | 0.0129 | 1.6379 | 0.7119 |
| ε-Carotene | 0.32 | 0.45 | 0.64 | 1.18 | 1.07 | 0.99 | 0.0088 | 2.2854 | 1.1925 |
| Lutein myristate | 0.07 | 0.08 | 0.07 | 0.19 | 0.08 | 0.07 | 0.3977 | 1.5650 | 0.6461 |
| Violaxanthin laurate | 0.02 | 0.03 | 0.04 | 0.08 | 0.10 | 0.11 | 0.0031 | 3.5034 | 1.8087 |
| Violaxanthin myristate | 13.05 | 11.83 | 11.55 | 43.48 | 24.11 | 26.54 | 0.0865 | 2.5839 | 1.3695 |
| Zeaxanthin myristoleate | 0.04 | 0.05 | 0.05 | N/A | N/A | N/A | N/A | N/A | N/A |
| Zeaxanthin palmitate | 3.50 | 3.11 | 2.34 | 0.85 | 1.18 | 1.31 | 0.0201 | 0.3740 | -1.4189 |
| Antheraxanthin | 0.26 | 0.43 | 0.50 | 0.98 | 1.19 | 1.24 | 0.0026 | 2.8554 | 1.5137 |
| Apocarotenal | 0.05 | 0.03 | 0.04 | 0.08 | 0.06 | 0.06 | 0.0641 | 1.5979 | 0.6762 |
| Canthaxanthin | 0.18 | 0.11 | 0.11 | 0.26 | 0.25 | 0.21 | 0.0254 | 1.8273 | 0.8697 |
| Capsanthin | 0.10 | 0.11 | 0.10 | 0.43 | 0.19 | 0.18 | 0.1822 | 2.5528 | 1.3521 |
| Echinenone | 35.18 | 24.02 | 21.73 | 45.79 | 32.51 | 33.32 | 0.1622 | 1.3792 | 0.4639 |
| Zeaxanthin | 8.35 | 7.30 | 12.04 | 28.16 | 22.62 | 23.14 | 0.0029 | 2.6692 | 1.4164 |
| β-Cryptoxanthin | 12.38 | 7.28 | 5.99 | 14.47 | 9.33 | 9.52 | 0.3791 | 1.2983 | 0.3766 |
| Fucoxanthin | 2.83 | 2.12 | 2.75 | 5.28 | 5.49 | 6.01 | 0.0091 | 2.1476 | 1.1028 |

**Table S21** Differentially accumulated carotenoids compounds in the comparison of 7d-W vs. 7d-G by targeted metabolomics (n=3)

| Compound | Class | Pvalue | FoldChange | Log2FoldChange |  |
| --- | --- | --- | --- | --- | --- |
| ε-Carotene | carotenes | 0.0088 | 2.2854 | 1.1925 | up |
| violaxanthin laurate | carotenoid esters | 0.0031 | 3.5034 | 1.8087 | up |
| violaxanthin myristate | carotenoid esters | 0.0087 | 2.5839 | 1.3695 | up |
| zeaxanthin palmitate | carotenoid esters | 0.0201 | 0.3740 | -1.4189 | down |
| antheraxanthin | xanthophylls | 0.0026 | 2.8554 | 1.5137 | up |
| capsanthin | xanthophylls | 0.0018 | 2.5528 | 1.3521 | up |
| zeaxanthin | xanthophylls | 0.0029 | 2.6692 | 1.4164 | up |
| fucoxanthin | xanthophylls | 0.0091 | 2.1476 | 1.1028 | up |

**Table S22** Statistical analysis of transcriptome data

| Sample | Repeat | Total_reads | Total_bases | GC_content | Q20 | Q30 |
| --- | --- | --- | --- | --- | --- | --- |
| C9d | C_9_r1 | 67,821,102 | 10,173,165,300 | 60.12% | 96.11% | 90.83% |
|  | C_9_r2 | 74,208,330 | 11,131,249,500 | 60.33% | 96.10% | 90.82% |
|  | C_9_r3 | 83,696,476 | 12,554,471,400 | 60.12% | 96.14% | 90.85% |
| C7d | C_7_r1 | 108,882,342 | 16,332,351,300 | 60.70% | 95.50% | 89.52% |
|  | C_7_r2 | 90,029,112 | 13,504,366,800 | 60.14% | 95.83% | 90.21% |
|  | C_7_r3 | 75,430,952 | 11,314,642,800 | 60.14% | 95.97% | 90.52% |
| C5d | C_5_r1 | 109,665,534 | 16,449,830,100 | 60.49% | 96.67% | 92.29% |
|  | C_5_r2 | 93,155,890 | 13,973,383,500 | 60.26% | 95.61% | 89.76% |
|  | C_5_r3 | 83,308,448 | 12,496,267,200 | 59.98% | 95.96% | 90.48% |
| C3d | C_3_r1 | 82,271,478 | 12,340,721,700 | 60.11% | 96.11% | 90.82% |
|  | C_3_r2 | 85,417,812 | 12,812,671,800 | 59.83% | 96.66% | 92.01% |
|  | C_3_r3 | 92,750,238 | 13,912,535,700 | 60.13% | 97.01% | 93.02% |
| T9d | T-9_r1 | 82,128,488 | 12,319,273,200 | 60.07% | 96.26% | 91.01% |
|  | T-9_r2 | 76,520,248 | 11,478,037,200 | 60.30% | 94.51% | 87.56% |
|  | T-9_r3 | 91,676,842 | 13,751,526,300 | 56.83% | 96.18% | 90.83% |
| T7d | T-7_r1 | 75,864,106 | 11,379,615,900 | 60.06% | 96.38% | 91.33% |
|  | T-7_r2 | 77,804,744 | 11,670,711,600 | 60.01% | 96.19% | 90.93% |
|  | T-7_r3 | 66,379,212 | 9,956,881,800 | 60.42% | 96.82% | 92.32% |
| T5d | T-5_r1 | 82,685,654 | 12,402,848,100 | 60.07% | 96.27% | 91.14% |
|  | T-5_r2 | 83,617,610 | 12,542,641,500 | 60.47% | 97.05% | 92.83% |
|  | T-5_r3 | 65,585,916 | 9,837,887,400 | 60.15% | 95.98% | 90.44% |
| T3d | T-3_r1 | 118,223,716 | 17,733,557,400 | 59.75% | 96.96% | 92.55% |
|  | T-3_r2 | 109,416,042 | 16,412,406,300 | 60.52% | 96.38% | 91.34% |
|  | T-3_r3 | 75,262,158 | 11,289,323,700 | 60.02% | 95.86% | 90.21% |

**Table S23** Mapping data of each transcriptome sample to the generated genome assembly

| Sample | Repeat | Total_reads | Unique_mapped | Multiple_mapped | Unmapped |
| --- | --- | --- | --- | --- | --- |
| C9d | C_9_r1 | 33910551 | 93.62 | 1.54 | 4.84 |
|  | C_9_r2 | 37104165 | 93.45 | 1.52 | 5.03 |
|  | C_9_r3 | 41848238 | 93.41 | 1.5 | 5.09 |
| C7d | C_7_r1 | 54441171 | 92.25 | 1.79 | 5.96 |
|  | C_7_r2 | 45014556 | 92.68 | 1.5 | 5.82 |
|  | C_7_r3 | 37715476 | 93.38 | 1.77 | 4.85 |
| C5d | C_5_r1 | 54832767 | 92.76 | 1.7 | 5.54 |
|  | C_5_r2 | 46577945 | 93.34 | 1.68 | 4.98 |
|  | C_5_r3 | 41654224 | 92.6 | 1.93 | 5.47 |
| C3d | C_3_r1 | 41135739 | 93.56 | 1.42 | 5.02 |
|  | C_3_r2 | 42708906 | 93.48 | 1.45 | 5.07 |
|  | C_3_r3 | 46375119 | 93.63 | 1.61 | 4.76 |
| T9d | T_9_r1 | 41064244 | 93.68 | 1.64 | 4.68 |
|  | T_9_r2 | 38260124 | 92.53 | 1.65 | 5.82 |
|  | T_9_r3 | 45838421 | 91.11 | 1.84 | 7.05 |
| T7d | T_7_r1 | 37932053 | 93.96 | 1.61 | 4.43 |
|  | T_7_r2 | 38902372 | 93.37 | 1.59 | 5.04 |
|  | T_7_r3 | 33189606 | 93.24 | 1.81 | 4.95 |
| T5d | T_5_r1 | 41342827 | 93.83 | 1.52 | 4.65 |
|  | T_5_r2 | 41808805 | 93.61 | 1.68 | 4.71 |
|  | T_5_r3 | 32792958 | 93.35 | 1.6 | 5.05 |
| T3d | T_3_r1 | 59111858 | 93.65 | 1.72 | 4.63 |
|  | T_3_r2 | 54708021 | 93.16 | 1.65 | 5.19 |
|  | T_3_r3 | 37631079 | 93.47 | 1.45 | 5.08 |
