## Supplementary for "Multi-omics Analyses Provide Insight into the Biosynthesis Pathways of Fucoxanthin in *Isochrysis galbana*": Supplementary tables.docx

**Table S1** Sequencing data used for *I. galbana* LG007 genome construction

| Library  resource | Sequencing platform | Insert size (bp) | Clean data (Gb) | Sequence coverage (X) | Use of the data |
| --- | --- | --- | --- | --- | --- |
| Genome | Illumina HiSeq X Ten | 250 bp | 8.92 | 96 | Genome estimation and polishing |
| Genome | PacBio SEQUEL | 20 kb | 15.53 | 166 | Genome assembly |
| Hi-C | Illumina HiSeq X Ten | 250 bp | 12.35 | 137 | Chromosome construction |
| Transcriptome | Illumina HiSeq X Ten | 0.6-3 kb | 307.77 | - | Difference analysis and annotation |

**Table S2** Assembly statistics for nuclear genome

| Items | Statistics |
| --- | --- |
| Estimated genome size | 92.59 Mb |
| Genomic G+C content | 58.44% |
| Number of assembled scaffolds | 353 |
| Number of scaffolds > 2 kb | 353 |
| min length (bp) | 5,065 |
| max length (bp) | 2,925,745 |
| Average length (bp) | 262,298 |
| Scaffolds N50 | 666.7 kb |
| Predicted gene models | 14,900 |
| Average coding sequence length (bp) | 1,428 |
| Average transcript length (bp) | 1,764 |
| Average gene length (bp) | 1,789 |
| Chromosomal assembled N50 | 6.99 Mb |

**Table S3** Assessment of the completeness of the *I. galbana* LG007 genome assembly by BUSCO

| Type | Number | Percent (%) |
| --- | --- | --- |
| Complete BUSCOs (C) | 254 | 83.8 |
| Complete and single-copy BUSCOs (S) | 248 | 81.8 |
| Complete and duplicated BUSCOs (D) | 6 | 2.0 |
| Fragmented BUSCOs (F) | 13 | 4.3 |
| Missing BUSCOs (M) | 36 | 11.9 |
| Total BUSCO groups searched | 303 | 100 |

**Table S4** Genomic resequencing alignment rate and coverage assessment

| Items | Statistics |
| --- | --- |
| Unmapped bases (Mb) | 144.6 |
| Mapped bases (Mb) | 887.5 |
| Map rate (%) | 98.4 |
| Genome Length (Mb) | 93 |
| Mean Depth | 89 |
| Coverage Rate (%) | 97.6 |

**Table S5** Pacbio data alignment rate

| Items | Statistics |
| --- | --- |
| QC-passed reads | 288211 |
| Duplicates | 0 |
| Mapped reads | 287594 |
| Mapping rate (%) | 99.78 |

**Table S6** Pseudomolecule length statistics after Hi-C assisted assembly

| Pseudomolecule | Length/bp |
| --- | --- |
| Chr1 | 11908665 |
| Chr2 | 10401026 |
| Chr3 | 7919487 |
| Chr4 | 7664558 |
| Chr5 | 7036706 |
| Chr6 | 6994209 |
| Chr7 | 6185998 |
| Chr8 | 5984543 |
| Chr9 | 5214086 |
| Chr10 | 4726453 |
| Chr11 | 4149466 |
| Chr12 | 3870612 |
| Chr13 | 3512008 |
| Chr14 | 3046428 |
| Chr15 | 2341788 |
| Total anchored | 90956033 |
| Unanchored | 1777877 |

**Table S7** Quality assembly statistics of the *I. galbana* after the Hi-C data based pseudo-chromosome assembly

| Items | Canu | | Hi-C | |
| --- | --- | --- | --- | --- |
|  | Contig_len (Mb) | Contig_number | Contig_len (Mb) | Contig_number |
| Total | 92.59 | 353 | 92.64 | 142 |
| Max | 2.92 | - | 11.91 | - |
| Number>=2 kb | - | 353 | - | 78 |
| N50 | 0.66 | - | 6.99 | - |

**Table S8** Statistics for gene, exon, CDS and introne in *I. galbana* LG007 genome

|  | Gene stat | Exons stat | CDS stat | Introne stat |
| --- | --- | --- | --- | --- |
| Total length | 26,666,990 | 21,283,343 | 21,283,343 | 5,383,647 |
| Total number | 14,900 | 43,481 | 14,900 | 28,581 |
| Average length | 1,789.73 | 489.49 | 1,428.41 | 188.36 |

**Table S9** Annotation statistics for the *I. galbana* LG007 genome

| **Annotation statistics for nuclear genome** | | Number | Percent (%) |
| --- | --- | --- | --- |
|  | Total protein | 14,900 |  |
|  | NR | 12,469 | 83.68 |
|  | eggNOG | 9,161 | 61.48 |
|  | GO | 4,977 | 33.40 |
|  | COG | 9,161 | 61.48 |
|  | KEGG | 3,773 | 25.32 |
|  | At least in one database | 12,500 | 83.89 |

**Table S10** Transcription factors

| Number | Classify |
| --- | --- |
| 198 | protein kinase family protein |
| 55 | heat shock protein |
| 49 | WD-40 repeat family protein / zfwd4 protein (ZFWD4) |
| 44 | myb domain protein 3r-3 |
| 23 | pentatricopeptide (PPR) repeat-containing protein |
| 13 | Calmodulin-binding transcription activator protein with CG-1 and Ankyrin domains |
| 11 | CCCH-type zinc finger protein with ARM repeat domain |
| 7 | DnaJ domain ;Myb-like DNA-binding domain |
| 7 | Zinc finger (C2H2 type) family protein / transcription factor jumonji (jmj) family protein |
| 6 | Homeodomain-like superfamily protein |
| 6 | ethylene induced calmodulin binding protein |
| 6 | pathogenesis related homeodomain protein A |
| 3 | GATA transcription factor 15 |
| 3 | E2F transcription factor |
| 2 | winged-helix DNA-binding transcription factor family protein |
| 2 | Plant-specific transcription factor YABBY family protein |
| 1 | AP2/B3-like transcriptional factor family protein |
| 1 | HD-ZIP IV family of homeobox-leucine zipper protein with lipid-binding START domain |
| 1 | Homeobox-leucine zipper protein 4 (HB-4) / HD-ZIP protein |
| 1 | K-box region and MADS-box transcription factor family protein |

**Table S11** Statistical analysis of non-coding RNAs in *I. galbana* LG007 genome

| ncRNA Items | Number |
| --- | --- |
| tRNA | 95 |
| rRNA | 58 |
| snRNA | 4 |

**Table S12** Repetitive element annotations in the *I. galbana* LG007

|  | No. of TEs | Length (bp) | % of TEs | % of genome |
| --- | --- | --- | --- | --- |
| Total repeat fraction | 161907 | 43349564 | 100 | 46.82 |
| Class I: Retroelement | 73047 | 31195044 | 71.96 | 33.69 |
| LTR Retrotransposon | 27946 | 14219065 | 32.8 | 15.36 |
| Ty1/Copia | 1992 | 998417 | 2.3 | 1.08 |
| Ty3/Gypsy | 4491 | 4191464 | 9.67 | 4.53 |
| Other | 21463 | 9029184 | 20.83 | 9.75 |
| non-LTR Retrotransposon | 26756 | 11607249 | 26.78 | 12.54 |
| LINE | 21408 | 10762090 | 24.83 | 11.62 |
| SINE | 5348 | 845159 | 1.95 | 0.91 |
| unclassified retroelement | 18345 | 5368730 | 12.38 | 5.8 |
| Class II: DNA Transposon | 33472 | 7828746 | 18.06 | 8.46 |
| CMC | 1176 | 203681 | 0.47 | 0.22 |
| hAT | 4039 | 1494270 | 3.45 | 1.61 |
| Mutator | 629 | 121730 | 0.28 | 0.13 |
| Tc1/Mariner | 180 | 71173 | 0.16 | 0.08 |
| PIF/Harbinger | 242 | 98683 | 0.23 | 0.11 |
| Other | 27026 | 5768036 | 13.31 | 6.23 |
| Helitron | 69 | 12880 | 0.03 | 0.01 |
| Tandem Repeats | 43633 | 3981448 | 9.18 | 4.3 |
| Unkown | 2891 | 1025862 | 2.37 | 1.11 |

**Table S13** KEGG enrichment of the expanded families genes identified in *I. galbana*

| KEGG_A_class | KEGG_B_class | Pathway | number | p-value | q-value | Genes |
| --- | --- | --- | --- | --- | --- | --- |
| Environmental Information Processing | Membrane transport | ABC transporters | 9 | 4.35E-11 | 1.57E-09 | IZ004963;IZ005238;IZ010481;IZ011657;IZ001888;IZ004017;  IZ007255;IZ008011;IZ011853 |
| Environmental Information Processing | Signal transduction | Two-component system | 9 | 1.16E-10 | 2.09E-09 | IZ004963;IZ005238;IZ010481;IZ011657;IZ001888;IZ004017;  IZ007255;IZ008011;IZ011853 |
| Metabolism | Nucleotide metabolism | Purine metabolism | 5 | 0.000982026 | 7.07E-03 | IZ003969;IZ012145;IZ012146;IZ014649;IZ002615 |
| Metabolism | Global and overview maps | Microbial metabolism in diverse environments | 3 | 0.3034138 | 4.57E-01 | IZ001240;IZ007901;IZ010956 |
| Metabolism | Global and overview maps | Metabolic pathways | 12 | 0.3603783 | 4.99E-01 | IZ001240;IZ003969;IZ004051;IZ006575;IZ007901;IZ010956;  IZ012145;IZ012146;IZ012985;IZ014649;IZ014829;IZ002615 |
| Metabolism | Global and overview maps | Biosynthesis of secondary metabolites | 3 | 0.8444505 | 8.44E-01 | IZ001240;IZ004051;IZ014829 |
| Metabolism | Carbohydrate metabolism | Glyoxylate and dicarboxylate metabolism | 2 | 0.04869659 | 1.59E-01 | IZ007901;IZ010956 |
| Metabolism | Carbohydrate metabolism | Pentose phosphate pathway | 2 | 0.0564664 | 1.69E-01 | IZ007901;IZ010956 |
| Metabolism | Amino acid metabolism | Lysine degradation | 2 | 0.07030375 | 1.88E-01 | IZ006575;IZ012985 |
| Metabolism | Lipid metabolism | Glycerophospholipid metabolism | 2 | 0.07319447 | 1.88E-01 | IZ004051;IZ014829 |
| Metabolism | Global and overview maps | Carbon metabolism | 2 | 0.4720631 | 5.48E-01 | IZ007901;IZ010956 |
| Cellular Processes | Cellular community - eukaryotes | Focal adhesion | 1 | 0.2812106 | 4.57E-01 | IZ006370 |
| Metabolism | Metabolism of cofactors and vitamins | Porphyrin and chlorophyll metabolism | 1 | 0.3104506 | 4.57E-01 | IZ001240 |
| Cellular Processes | Cell motility | Regulation of actin cytoskeleton | 1 | 0.3175789 | 4.57E-01 | IZ006370 |
| Environmental Information Processing | Signal transduction | Calcium signaling pathway | 1 | 0.4343465 | 5.21E-01 | IZ006370 |
| Environmental Information Processing | Signal transduction | cGMP - PKG signaling pathway | 1 | 0.4343465 | 5.21E-01 | IZ006370 |
| Environmental Information Processing | Signal transduction | PI3K-Akt signaling pathway | 1 | 0.5115662 | 5.69E-01 | IZ000809 |
| Environmental Information Processing | Signal transduction | Apelin signaling pathway | 1 | 0.5217293 | 5.69E-01 | IZ006370 |

**Table S14** GO enrichment of the expanded families genes identified in *I. galbana*

| GO Term | number | p-value | p-adjust | Genes_id |
| --- | --- | --- | --- | --- |
| response to stimulus | 22 | 0.243762872 | 0.566807701 | IZ000514,IZ000648,IZ001888,IZ002346,IZ002462,IZ003775,IZ003799,IZ004017,IZ004500,IZ004747,  IZ004761,IZ004963,IZ006187,IZ008011,IZ010376,IZ011657,IZ012145,IZ012146,IZ012297,IZ013171,  IZ013806,IZ014649 |
| cellular process | 40 | 0.270129153 | 0.5980009 | IZ000359,IZ000514,IZ000648,IZ001888,IZ002145,IZ002346,IZ002462,IZ003225,IZ003288,IZ003775,  IZ003799,IZ004017,IZ004051,IZ004500,IZ004747,IZ004761,IZ004963,IZ005121,IZ006575,IZ006892,  IZ007042,IZ007195,IZ007901,IZ007933,IZ008011,IZ009320,IZ010260,IZ010376,IZ010634,IZ010956,  IZ011442,IZ011657,IZ012145,IZ012146,IZ012297,IZ012328,IZ013171,IZ013806,IZ014649,IZ014829 |
| signaling | 9 | 0.337365388 | 0.661549398 | IZ002145,IZ003775,IZ004500,IZ004761,IZ010376,IZ012145,IZ012146,IZ013806,IZ014649 |
| single-organism process | 37 | 0.341909372 | 0.666198336 | IZ000359,IZ000514,IZ000648,IZ001888,IZ002145,IZ002346,IZ002462,IZ003288,IZ003775,IZ003799,  IZ004017,IZ004051,IZ004500,IZ004747,IZ004761,IZ004963,IZ005121,IZ006575,IZ006644,IZ006892,  IZ007042,IZ007195,IZ007901,IZ008011,IZ009320,IZ010260,IZ010376,IZ010634,IZ010956,IZ011657,  IZ012145,IZ012146,IZ012328,IZ013171,IZ013806,IZ014649,IZ014829 |
| immune system process | 4 | 0.350905609 | 0.675927737 | IZ004500,IZ004761,IZ010376,IZ013806 |
| biological adhesion | 2 | 0.441001746 | 0.762653691 | IZ004761,IZ010376 |
| localization | 12 | 0.740589615 | 0.962573382 | IZ000648,IZ001888,IZ002346,IZ003799,IZ004017,IZ004500,IZ004963,IZ008011,IZ010376,IZ011657,  IZ013171,IZ013806 |
| negative regulation of biological process | 6 | 0.830490294 | 1 | IZ004747,IZ004761,IZ010376,IZ012145,IZ012146,IZ014649 |
| metabolic process | 31 | 0.832199171 | 1 | IZ000359,IZ000514,IZ001888,IZ003225,IZ003288,IZ003775,IZ003875,IZ004017,IZ004051,IZ004747,  IZ004761,IZ004963,IZ005121,IZ006575,IZ007042,IZ007195,IZ007901,IZ007933,IZ008011,IZ010260,  IZ010376,IZ010634,IZ010956,IZ011442,IZ011657,IZ012145,IZ012146,IZ012297,IZ012328,IZ014649,IZ014829 |
| developmental process | 14 | 0.905193048 | 1 | IZ000514,IZ000648,IZ002145,IZ002462,IZ004500,IZ006644,IZ006892,IZ009320,IZ010376,IZ010634,  IZ012145,IZ012146,IZ013806,IZ014649 |
| regulation of biological process | 13 | 0.964923709 | 1 | IZ000359,IZ002462,IZ003225,IZ003775,IZ004500,IZ004747,IZ004761,IZ010376,IZ012145,IZ012146,  IZ012297,IZ013806,IZ014649 |
| reproductive process | 4 | 0.965536855 | 1 | IZ000514,IZ002462,IZ005121,IZ010376 |
| multi-organism process | 3 | 0.990061135 | 1 | IZ004500,IZ005121,IZ013806 |
| cellular component organization or biogenesis | 9 | 0.996109803 | 1 | IZ000359,IZ002462,IZ004500,IZ006575,IZ006892,IZ009320,IZ010376,IZ012328,IZ013806 |
| growth | 2 | 0.996700725 | 1 | IZ002462,IZ010376 |
| multicellular organismal process | 9 | 0.997883195 | 1 | IZ000514,IZ000648,IZ002462,IZ004500,IZ005121,IZ006644,IZ010376,IZ010634,IZ013806 |
| transporter activity | 9 | 0.008793802 | 0.044820026 | IZ000648,IZ001888,IZ002346,IZ003799,IZ004017,IZ004963,IZ008011,IZ011657,IZ013171 |
| structural molecule activity | 3 | 0.224648199 | 0.394382393 | IZ004500,IZ010376,IZ013806 |
| catalytic activity | 23 | 0.293882517 | 0.465459075 | IZ000514,IZ001888,IZ003288,IZ003775,IZ003875,IZ004017,IZ004051,IZ004747,IZ004963,IZ005121,  IZ006575,IZ007042,IZ007195,IZ007901,IZ007933,IZ008011,IZ010260,IZ010634,IZ010956,IZ011657,  IZ012297,IZ012328,IZ014829 |
| binding | 8 | 0.996797129 | 1 | IZ002462,IZ003288,IZ003775,IZ007042,IZ007195,IZ010260,IZ010376,IZ011506 |

**Table S15** KEGG enrichment of the contracted families genes identified in *I. galbana*

| KEGG_A_class | KEGG_B_class | Pathway | number | p-value | q-value | Genes |
| --- | --- | --- | --- | --- | --- | --- |
| Metabolism | Global and overview maps | Metabolic pathways | 48 | 0.2857671 | 0.76937296 | IZ000038;IZ005085;IZ007554;IZ008968;IZ010315;IZ013625;IZ014320;IZ000679;IZ000715;IZ001340;IZ001871;IZ002279;IZ002348;IZ002423;IZ003695;IZ003733;IZ004326;IZ006003;IZ006156;IZ006366;IZ007805;IZ008796;IZ009611;IZ010613;IZ011172;IZ011447;IZ011461;IZ012261;IZ013277;IZ013320;IZ013355;IZ013965;IZ014766;IZ000798;IZ001487;IZ001634;IZ002116;IZ002532;IZ002557;IZ002560;IZ003022;IZ003431;IZ003450;IZ003519;IZ003770;IZ005551;IZ006679;IZ007055 |
| Metabolism | Global and overview maps | Biosynthesis of secondary metabolites | 13 | 0.9494193 | 0.96678484 | IZ005085;IZ007554;IZ008968;IZ006003;IZ010613;IZ011172;IZ013355;IZ000798;IZ001634;IZ002116;IZ003022;IZ003431;IZ007055 |
| Cellular Processes | Cellular community - eukaryotes | Signaling pathways regulating pluripotency of stem cells | 12 | 0.001895077 | 0.09285877 | IZ000281;IZ006649;IZ008132;IZ008236;IZ008481;IZ009625;IZ009635;IZ010858;IZ013166;IZ001478;IZ001996;IZ007056 |
| Cellular Processes | Cellular community - prokaryotes | Quorum sensing | 7 | 0.000666482 | 0.07805154 | IZ000284;IZ000585;IZ002727;IZ007996;IZ010843;IZ012082;IZ003431 |
| Genetic Information Processing | Folding, sorting and degradation | RNA degradation | 7 | 0.03066147 | 0.36880217 | IZ002757;IZ003347;IZ006578;IZ006823;IZ011122;IZ013774;IZ001933 |
| Metabolism | Carbohydrate metabolism | Inositol phosphate metabolism | 6 | 0.000955733 | 0.07805154 | IZ010315;IZ013625;IZ014320;IZ011461;IZ003431;IZ005551 |
| Metabolism | Carbohydrate metabolism | Starch and sucrose metabolism | 6 | 0.01021047 | 0.23664661 | IZ005085;IZ006003;IZ006156;IZ007805;IZ002116;IZ002557 |
| Metabolism | Nucleotide metabolism | Purine metabolism | 5 | 0.2411962 | 0.70482811 | IZ007554;IZ003733;IZ013277;IZ001634;IZ002532 |
| Genetic Information Processing | Translation | RNA transport | 5 | 0.3425649 | 0.77735109 | IZ000543;IZ003347;IZ006359;IZ007863;IZ001470 |
| Metabolism | Global and overview maps | Microbial metabolism in diverse environments | 5 | 0.9181361 | 0.96129634 | IZ002279;IZ009611;IZ003450;IZ006679;IZ007055 |
| Environmental Information Processing | Signal transduction | VEGF signaling pathway | 4 | 0.003928481 | 0.13749684 | IZ002694;IZ004459;IZ006649;IZ011461 |
| Metabolism | Nucleotide metabolism | Pyrimidine metabolism | 4 | 0.1101915 | 0.53924113 | IZ000038;IZ007554;IZ001487;IZ001634 |
| Metabolism | Lipid metabolism | Glycerophospholipid metabolism | 4 | 0.1173991 | 0.53924113 | IZ013355;IZ003022;IZ003431;IZ003770 |
| Environmental Information Processing | Signal transduction | Phosphatidylinositol signaling system | 4 | 0.1173991 | 0.53924113 | IZ013625;IZ014320;IZ011461;IZ005551 |
| Environmental Information Processing | Signal transduction | Calcium signaling pathway | 4 | 0.2083574 | 0.67903652 | IZ014320;IZ004459;IZ011461;IZ005551 |
| Metabolism | Lipid metabolism | Ether lipid metabolism | 3 | 0.002375428 | 0.09699664 | IZ003022;IZ003431;IZ003770 |
| Environmental Information Processing | Signal transduction | ErbB signaling pathway | 3 | 0.03922449 | 0.36880217 | IZ006649;IZ011461;IZ001478 |
| Environmental Information Processing | Membrane transport | ABC transporters | 3 | 0.2214357 | 0.69553521 | IZ000284;IZ010843;IZ012082 |
| Environmental Information Processing | Signal transduction | Ras signaling pathway | 3 | 0.2214357 | 0.69553521 | IZ006649;IZ010072;IZ011461 |
| Environmental Information Processing | Signal transduction | MAPK signaling pathway | 3 | 0.2329303 | 0.70482811 | IZ004459;IZ006649;IZ002944 |
| Genetic Information Processing | Replication and repair | DNA replication | 3 | 0.2445322 | 0.70482811 | IZ002742;IZ010707;IZ001607 |
| Genetic Information Processing | Folding, sorting and degradation | Protein export | 3 | 0.2445322 | 0.70482811 | IZ000585;IZ002727;IZ007996 |
| Environmental Information Processing | Signal transduction | Two-component system | 3 | 0.2679983 | 0.74613163 | IZ009087;IZ006156;IZ010613 |
| Environmental Information Processing | Signal transduction | mTOR signaling pathway | 3 | 0.433926 | 0.79308479 | IZ003677;IZ006649;IZ001478 |
| Environmental Information Processing | Signal transduction | PI3K-Akt signaling pathway | 3 | 0.5749061 | 0.83821506 | IZ003677;IZ006649;IZ001478 |

**Table S16** GO enrichment of the contracted families genes identified in *I. galbana*

| GO Term | number | p-value | p-adjust | Genes_id |
| --- | --- | --- | --- | --- |
| localization | 60 | 0.025172212 | 0.443808946 | IZ000165,IZ000398,IZ000456,IZ000461,IZ000543,IZ000676,IZ000789,IZ000899,IZ001470,IZ001634,IZ002742,IZ003002,IZ003194,IZ003292,IZ003347,IZ003677,IZ003697,IZ003723,IZ004425,IZ004450,IZ004483,IZ004518,IZ004604,IZ005010,IZ005345,IZ005471,IZ005707,IZ006011,IZ006147,IZ006366,IZ006511,IZ006767,IZ006904,IZ007080,IZ007874,IZ007954,IZ007996,IZ008656,IZ008689,IZ008969,IZ009076,IZ009179,IZ009423,IZ009843,IZ010530,IZ010629,IZ011568,IZ011808,IZ012570,IZ012575,IZ013208,IZ013327,IZ013506,IZ013662,IZ013838,IZ013900,IZ014197,IZ014636,IZ014701,IZ014862 |
| negative regulation of biological process | 38 | 0.029538555 | 0.443808946 | IZ000029,IZ000398,IZ000456,IZ000461,IZ001229,IZ001396,IZ001634,IZ001933,IZ002560,IZ002742,IZ002991,IZ003347,IZ004475,IZ004972,IZ005471,IZ006147,IZ006366,IZ006511,IZ006649,IZ006962,IZ007554,IZ007781,IZ007789,IZ007863,IZ007874,IZ008743,IZ008969,IZ009179,IZ009773,IZ009809,IZ011479,IZ011568,IZ012018,IZ012276,IZ013542,IZ014636,IZ014766,IZ014852 |
| immune system process | 17 | 0.037146286 | 0.443808946 | IZ000029,IZ000456,IZ000461,IZ000789,IZ001229,IZ001470,IZ001634,IZ002991,IZ004604,IZ005471,IZ009179,IZ009809,IZ012018,IZ012276,IZ014320,IZ014605,IZ014636 |
| biological adhesion | 10 | 0.038756261 | 0.443808946 | IZ000461,IZ001229,IZ001634,IZ002991,IZ005471,IZ005888,IZ009809,IZ012018,IZ014320,IZ014636 |
| biological regulation | 78 | 0.116204806 | 0.526019172 | IZ000029,IZ000165,IZ000398,IZ000456,IZ000461,IZ000543,IZ000789,IZ000899,IZ001229,IZ001396,IZ001634,IZ001918,IZ001933,IZ001953,IZ002074,IZ002560,IZ002742,IZ002991,IZ003347,IZ003677,IZ003723,IZ004213,IZ004450,IZ004459,IZ004475,IZ004604,IZ004972,IZ005010,IZ005345,IZ005471,IZ005707,IZ006011,IZ006147,IZ006366,IZ006367,IZ006511,IZ006649,IZ006904,IZ006962,IZ007554,IZ007781,IZ007789,IZ007863,IZ007874,IZ007954,IZ008689,IZ008743,IZ008969,IZ008987,IZ009076,IZ009179,IZ009773,IZ009809,IZ009843,IZ010218,IZ010629,IZ010707,IZ011479,IZ011568,IZ011676,IZ012018,IZ012276,IZ012570,IZ012575,IZ013208,IZ013506,IZ013542,IZ013555,IZ013662,IZ013838,IZ013900,IZ013972,IZ014320,IZ014605,IZ014636,IZ014766,IZ014852,IZ014862 |
| regulation of biological process | 73 | 0.118227569 | 0.534312338 | IZ000029,IZ000165,IZ000398,IZ000456,IZ000461,IZ000543,IZ000789,IZ000899,IZ001229,IZ001396,IZ001634,IZ001918,IZ001933,IZ001953,IZ002074,IZ002560,IZ002742,IZ002991,IZ003347,IZ003677,IZ003723,IZ004213,IZ004450,IZ004475,IZ004604,IZ004972,IZ005010,IZ005345,IZ005471,IZ005707,IZ006011,IZ006147,IZ006366,IZ006367,IZ006511,IZ006649,IZ006962,IZ007554,IZ007781,IZ007789,IZ007863,IZ007874,IZ008689,IZ008743,IZ008969,IZ008987,IZ009076,IZ009179,IZ009773,IZ009809,IZ009843,IZ010218,IZ010629,IZ010707,IZ011479,IZ011568,IZ011676,IZ012018,IZ012276,IZ012575,IZ013208,IZ013506,IZ013542,IZ013555,IZ013662,IZ013838,IZ013972,IZ014320,IZ014605,IZ014636,IZ014766,IZ014852,IZ014862 |
| multi-organism process | 33 | 0.16060266 | 0.576487193 | IZ000029,IZ000165,IZ000398,IZ000456,IZ000899,IZ001470,IZ001478,IZ001634,IZ001921,IZ002560,IZ002742,IZ004475,IZ005010,IZ006649,IZ007080,IZ007554,IZ007781,IZ007789,IZ007874,IZ007954,IZ008969,IZ009076,IZ009179,IZ009373,IZ009423,IZ012276,IZ012570,IZ013542,IZ013838,IZ014605,IZ014636,IZ014766,IZ014852 |
| positive regulation of biological process | 37 | 0.195233832 | 0.586825374 | IZ000029,IZ000398,IZ000461,IZ000543,IZ000789,IZ000899,IZ001396,IZ001634,IZ001918,IZ001953,IZ002074,IZ003347,IZ004604,IZ005010,IZ005345,IZ005471,IZ005707,IZ006147,IZ006366,IZ006367,IZ007781,IZ008969,IZ009076,IZ009179,IZ009773,IZ009843,IZ010629,IZ011568,IZ011676,IZ013208,IZ013542,IZ013555,IZ013972,IZ014320,IZ014605,IZ014636,IZ014852 |
| signaling | 29 | 0.368317898 | 0.662309811 | IZ000029,IZ000165,IZ000789,IZ000899,IZ001229,IZ001396,IZ001918,IZ002991,IZ003677,IZ004450,IZ004604,IZ005010,IZ006649,IZ006962,IZ007874,IZ007954,IZ008743,IZ008969,IZ009076,IZ009179,IZ009773,IZ009809,IZ010218,IZ010629,IZ012018,IZ013900,IZ014320,IZ014605,IZ014636 |
| reproductive process | 32 | 0.504252436 | 0.731781883 | IZ000165,IZ000398,IZ000461,IZ000899,IZ001478,IZ001634,IZ001921,IZ002560,IZ002742,IZ005010,IZ005345,IZ006366,IZ006511,IZ006649,IZ007080,IZ007132,IZ007554,IZ007579,IZ007874,IZ007954,IZ008969,IZ009076,IZ009179,IZ009373,IZ009423,IZ011568,IZ012570,IZ013208,IZ013838,IZ014636,IZ014766,IZ014852 |
| response to stimulus | 69 | 0.52931886 | 0.745470279 | IZ000029,IZ000165,IZ000398,IZ000456,IZ000461,IZ000789,IZ000899,IZ001229,IZ001396,IZ001470,IZ001593,IZ001634,IZ001918,IZ001921,IZ001933,IZ002074,IZ002560,IZ002731,IZ002742,IZ002991,IZ003292,IZ003677,IZ004425,IZ004450,IZ004459,IZ004475,IZ004604,IZ005010,IZ005345,IZ005471,IZ005707,IZ006649,IZ006962,IZ007080,IZ007554,IZ007781,IZ007789,IZ007863,IZ007874,IZ008743,IZ008969,IZ009076,IZ009179,IZ009373,IZ009423,IZ009611,IZ009773,IZ009809,IZ010218,IZ010629,IZ010707,IZ011315,IZ011479,IZ011568,IZ011808,IZ012018,IZ012276,IZ012793,IZ013208,IZ013327,IZ013506,IZ013555,IZ013838,IZ014197,IZ014320,IZ014605,IZ014636,IZ014852,IZ014862 |
| reproduction | 36 | 0.653690399 | 0.812978473 | IZ000165,IZ000398,IZ000461,IZ000899,IZ001478,IZ001634,IZ001921,IZ002116,IZ002560,IZ002742,IZ005010,IZ005345,IZ006366,IZ006367,IZ006511,IZ006649,IZ007080,IZ007132,IZ007554,IZ007579,IZ007781,IZ007874,IZ007954,IZ008969,IZ009076,IZ009179,IZ009373,IZ009423,IZ011568,IZ012570,IZ012575,IZ013208,IZ013838,IZ014636,IZ014766,IZ014852 |
| developmental process | 61 | 0.680271406 | 0.824803323 | IZ000165,IZ000398,IZ000461,IZ000543,IZ000899,IZ001478,IZ001593,IZ001634,IZ001836,IZ001918,IZ001921,IZ002116,IZ002560,IZ002742,IZ003347,IZ003723,IZ004992,IZ005004,IZ005010,IZ005345,IZ005471,IZ005888,IZ006147,IZ006366,IZ006367,IZ006962,IZ007080,IZ007132,IZ007554,IZ007579,IZ007781,IZ007789,IZ007874,IZ007925,IZ007954,IZ008969,IZ008987,IZ009076,IZ009120,IZ009179,IZ009373,IZ009423,IZ009534,IZ009592,IZ009773,IZ009843,IZ010629,IZ011315,IZ011568,IZ011663,IZ012570,IZ012575,IZ012675,IZ013186,IZ013208,IZ013313,IZ013520,IZ013662,IZ014320,IZ014636,IZ014766 |
| growth | 23 | 0.776575228 | 0.883027307 | IZ000165,IZ000398,IZ000461,IZ000543,IZ000899,IZ001634,IZ001918,IZ002116,IZ002572,IZ005345,IZ006366,IZ006367,IZ006649,IZ006687,IZ008969,IZ009076,IZ009179,IZ009843,IZ011568,IZ012575,IZ013186,IZ013208,IZ014368 |
| metabolic process | 113 | 0.88428477 | 0.943551381 | IZ000029,IZ000038,IZ000398,IZ000456,IZ000461,IZ000543,IZ000676,IZ000679,IZ000782,IZ000789,IZ000798,IZ000899,IZ001229,IZ001396,IZ001478,IZ001593,IZ001607,IZ001634,IZ001918,IZ001933,IZ001953,IZ002061,IZ002074,IZ002116,IZ002348,IZ002423,IZ002560,IZ002572,IZ002742,IZ002991,IZ003002,IZ003194,IZ003347,IZ003450,IZ003677,IZ003697,IZ003723,IZ003764,IZ003959,IZ004213,IZ004326,IZ004459,IZ004475,IZ004518,IZ004604,IZ004983,IZ005010,IZ005045,IZ005471,IZ005534,IZ005614,IZ006011,IZ006147,IZ006359,IZ006366,IZ006649,IZ006687,IZ006962,IZ007080,IZ007554,IZ007579,IZ007781,IZ007789,IZ007863,IZ007874,IZ007954,IZ007996,IZ008394,IZ008630,IZ008689,IZ008743,IZ008969,IZ009076,IZ009120,IZ009179,IZ009423,IZ009611,IZ009773,IZ009809,IZ010120,IZ010218,IZ010315,IZ010613,IZ010629,IZ010707,IZ011122,IZ011265,IZ011315,IZ011479,IZ011568,IZ011663,IZ011676,IZ012018,IZ012276,IZ012675,IZ013203,IZ013506,IZ013520,IZ013542,IZ013555,IZ013625,IZ013662,IZ013701,IZ013838,IZ013900,IZ013965,IZ013972,IZ014320,IZ014605,IZ014636,IZ014742,IZ014766,IZ014852 |
| cellular process | 131 | 0.940906453 | 0.981906846 | IZ000029,IZ000038,IZ000165,IZ000398,IZ000456,IZ000461,IZ000543,IZ000676,IZ000782,IZ000789,IZ000798,IZ000899,IZ001229,IZ001396,IZ001470,IZ001478,IZ001593,IZ001607,IZ001634,IZ001918,IZ001921,IZ001933,IZ001953,IZ002061,IZ002074,IZ002116,IZ002560,IZ002572,IZ002731,IZ002742,IZ002991,IZ003002,IZ003194,IZ003292,IZ003347,IZ003450,IZ003677,IZ003697,IZ003723,IZ003764,IZ004213,IZ004326,IZ004450,IZ004459,IZ004475,IZ004483,IZ004518,IZ004604,IZ004972,IZ004983,IZ005010,IZ005045,IZ005345,IZ005471,IZ005614,IZ005707,IZ005888,IZ006011,IZ006147,IZ006359,IZ006366,IZ006511,IZ006649,IZ006687,IZ006962,IZ007080,IZ007132,IZ007554,IZ007579,IZ007781,IZ007789,IZ007863,IZ007874,IZ007954,IZ007996,IZ008394,IZ008656,IZ008743,IZ008969,IZ008987,IZ009076,IZ009120,IZ009179,IZ009373,IZ009423,IZ009534,IZ009592,IZ009611,IZ009773,IZ009809,IZ009843,IZ010120,IZ010218,IZ010315,IZ010530,IZ010613,IZ010629,IZ010707,IZ011122,IZ011315,IZ011479,IZ011568,IZ011663,IZ011676,IZ011808,IZ012018,IZ012276,IZ012570,IZ012575,IZ012675,IZ013186,IZ013203,IZ013208,IZ013327,IZ013506,IZ013520,IZ013542,IZ013555,IZ013625,IZ013662,IZ013701,IZ013838,IZ013900,IZ013972,IZ014197,IZ014320,IZ014605,IZ014636,IZ014766,IZ014852,IZ014862 |
| transporter activity | 20 | 0.079917261 | 0.320298314 | IZ000165,IZ000789,IZ003002,IZ003194,IZ003292,IZ004425,IZ004450,IZ005345,IZ005707,IZ006767,IZ008656,IZ009843,IZ011808,IZ012575,IZ013208,IZ013327,IZ014197,IZ014636,IZ014701,IZ014862 |
| signal transducer activity | 4 | 0.242032917 | 0.472010555 | IZ004450,IZ006649,IZ010218,IZ014605 |
| molecular transducer activity | 4 | 0.324639032 | 0.547156515 | IZ004450,IZ006649,IZ010218,IZ014605 |
| binding | 59 | 0.458688945 | 0.626720387 | IZ000398,IZ000456,IZ000543,IZ000676,IZ001634,IZ001918,IZ001921,IZ001953,IZ002061,IZ002074,IZ002116,IZ002560,IZ002572,IZ002742,IZ003347,IZ003697,IZ004459,IZ004483,IZ005010,IZ005471,IZ005614,IZ005888,IZ006147,IZ006511,IZ006687,IZ007080,IZ007132,IZ007781,IZ007874,IZ007932,IZ007996,IZ008665,IZ008969,IZ009120,IZ009179,IZ009373,IZ009423,IZ009843,IZ010120,IZ010613,IZ010629,IZ011315,IZ011479,IZ011568,IZ011676,IZ012276,IZ013186,IZ013506,IZ013542,IZ013555,IZ013662,IZ013701,IZ013838,IZ013900,IZ013972,IZ014636,IZ014701,IZ014852,IZ014862 |
| catalytic activity | 72 | 0.970978715 | 0.984518259 | IZ000038,IZ000398,IZ000461,IZ000543,IZ000676,IZ000679,IZ000782,IZ000798,IZ000899,IZ001478,IZ001593,IZ001607,IZ001634,IZ001933,IZ002061,IZ002074,IZ002116,IZ002423,IZ002560,IZ002572,IZ002742,IZ003450,IZ003677,IZ003697,IZ003723,IZ003764,IZ003959,IZ004326,IZ004459,IZ004604,IZ004983,IZ005010,IZ005534,IZ005614,IZ006359,IZ006366,IZ006649,IZ007554,IZ007579,IZ007789,IZ007874,IZ007954,IZ007996,IZ008394,IZ008630,IZ009076,IZ009120,IZ009179,IZ009611,IZ009773,IZ010218,IZ010315,IZ010629,IZ010707,IZ011315,IZ011479,IZ011663,IZ011676,IZ012675,IZ013203,IZ013506,IZ013520,IZ013625,IZ013701,IZ013838,IZ013900,IZ013965,IZ014320,IZ014605,IZ014742,IZ014766,IZ014852 |

**Table S17** GO enrichment of the specific families genes identified in *I. galbana*

| GO Term | number | p-value | p-adjust | Genes_id |
| --- | --- | --- | --- | --- |
| signaling | 62 | 0.029881894 | 0.497376798 | IZ000131,IZ000616,IZ000832,IZ001275,IZ001762,IZ002145,IZ002439,IZ002585,IZ003737,IZ004491,IZ004500,IZ004558,IZ004755,IZ004758,IZ005641,IZ005716,IZ005748,IZ005782,IZ006009,IZ006524,IZ006718,IZ006924,IZ007013,IZ007081,IZ007094,IZ007424,IZ007569,IZ008124,IZ008982,IZ009360,IZ009581,IZ009744,IZ009927,IZ010039,IZ010158,IZ010218,IZ010221,IZ010629,IZ010983,IZ011738,IZ011929,IZ011981,IZ012000,IZ012018,IZ012171,IZ012220,IZ013362,IZ013426,IZ013466,IZ013537,IZ013600,IZ013806,IZ013946,IZ014053,IZ014202,IZ014344,IZ014404,IZ014447,IZ014509,IZ014575,IZ014670,IZ014843 |
| biological adhesion | 16 | 0.032914926 | 0.497376798 | IZ002353,IZ003631,IZ005471,IZ005716,IZ007013,IZ007424,IZ009927,IZ011415,IZ011981,IZ012018,IZ013362,IZ013466,IZ013719,IZ014344,IZ014447,IZ014843 |
| transcription factor activity, protein binding | 7 | 0.062512865 | 0.475406631 | IZ000171,IZ003902,IZ004601,IZ004898,IZ006718,IZ008403,IZ009140 |
| regulation of biological process | 131 | 0.098102922 | 0.54141515 | IZ000131,IZ000171,IZ000218,IZ000452,IZ000596,IZ000616,IZ000632,IZ000638,IZ000832,IZ001275,IZ001762,IZ001784,IZ001899,IZ002233,IZ002289,IZ002328,IZ002353,IZ002439,IZ002585,IZ002637,IZ003472,IZ003631,IZ003737,IZ003858,IZ003902,IZ004194,IZ004491,IZ004500,IZ004558,IZ004601,IZ004643,IZ004678,IZ004730,IZ004755,IZ004758,IZ004887,IZ004898,IZ005046,IZ005353,IZ005458,IZ005471,IZ005641,IZ005700,IZ005716,IZ005748,IZ005782,IZ006009,IZ006524,IZ006619,IZ006709,IZ006718,IZ006773,IZ006814,IZ006918,IZ006924,IZ007013,IZ007081,IZ007092,IZ007094,IZ007098,IZ007210,IZ007424,IZ007569,IZ007617,IZ007880,IZ008124,IZ008403,IZ008515,IZ008555,IZ008600,IZ008982,IZ009005,IZ009102,IZ009140,IZ009265,IZ009360,IZ009384,IZ009413,IZ009420,IZ009522,IZ009581,IZ009594,IZ009744,IZ009774,IZ009811,IZ009927,IZ009936,IZ010039,IZ010158,IZ010186,IZ010191,IZ010218,IZ010256,IZ010361,IZ010629,IZ010813,IZ010983,IZ011112,IZ011415,IZ011738,IZ011929,IZ011981,IZ012000,IZ012018,IZ012166,IZ012171,IZ012220,IZ012953,IZ013169,IZ013362,IZ013426,IZ013466,IZ013537,IZ013600,IZ013662,IZ013670,IZ013719,IZ013806,IZ013946,IZ014053,IZ014171,IZ014202,IZ014220,IZ014344,IZ014404,IZ014447,IZ014471,IZ014509,IZ014510,IZ014575,IZ014843 |
| biological regulation | 138 | 0.149471475 | 0.571095383 | IZ000131,IZ000171,IZ000218,IZ000452,IZ000596,IZ000616,IZ000632,IZ000638,IZ000832,IZ001275,IZ001762,IZ001784,IZ001899,IZ002233,IZ002289,IZ002328,IZ002353,IZ002439,IZ002491,IZ002585,IZ002637,IZ003472,IZ003631,IZ003737,IZ003858,IZ003902,IZ004194,IZ004491,IZ004500,IZ004558,IZ004601,IZ004643,IZ004678,IZ004730,IZ004755,IZ004758,IZ004887,IZ004898,IZ005046,IZ005353,IZ005458,IZ005471,IZ005641,IZ005700,IZ005704,IZ005716,IZ005748,IZ005782,IZ006009,IZ006524,IZ006619,IZ006709,IZ006718,IZ006773,IZ006814,IZ006918,IZ006924,IZ007013,IZ007081,IZ007092,IZ007094,IZ007098,IZ007210,IZ007424,IZ007569,IZ007617,IZ007880,IZ007970,IZ008124,IZ008403,IZ008515,IZ008555,IZ008600,IZ008982,IZ009005,IZ009102,IZ009140,IZ009265,IZ009360,IZ009384,IZ009413,IZ009420,IZ009522,IZ009581,IZ009594,IZ009736,IZ009744,IZ009774,IZ009811,IZ009927,IZ009936,IZ010039,IZ010158,IZ010186,IZ010191,IZ010218,IZ010221,IZ010256,IZ010361,IZ010629,IZ010813,IZ010983,IZ011112,IZ011415,IZ011738,IZ011929,IZ011981,IZ012000,IZ012018,IZ012166,IZ012171,IZ012220,IZ012953,IZ013169,IZ013362,IZ013426,IZ013466,IZ013537,IZ013600,IZ013662,IZ013670,IZ013719,IZ013806,IZ013946,IZ014053,IZ014147,IZ014171,IZ014202,IZ014220,IZ014344,IZ014404,IZ014447,IZ014471,IZ014509,IZ014510,IZ014575,IZ014670,IZ014843 |
| signal transducer activity | 7 | 0.183982708 | 0.604909354 | IZ006718,IZ007094,IZ010039,IZ010218,IZ010983,IZ013946,IZ014575 |
| membrane | 147 | 0.21309696 | 0.755104445 | IZ000049,IZ000318,IZ000385,IZ000570,IZ000608,IZ000673,IZ000702,IZ000832,IZ000974,IZ001108,IZ001146,IZ001241,IZ001260,IZ001382,IZ001437,IZ001455,IZ001524,IZ001540,IZ001687,IZ001762,IZ001784,IZ001898,IZ002002,IZ002328,IZ002614,IZ002627,IZ002764,IZ002988,IZ003047,IZ003053,IZ003192,IZ003308,IZ003339,IZ003472,IZ003486,IZ003631,IZ003737,IZ003954,IZ004112,IZ004146,IZ004194,IZ004491,IZ004558,IZ004609,IZ004643,IZ004678,IZ004758,IZ004978,IZ005316,IZ005353,IZ005520,IZ005545,IZ005641,IZ005704,IZ005705,IZ005782,IZ005797,IZ005942,IZ005991,IZ006197,IZ006335,IZ006467,IZ006524,IZ006619,IZ006644,IZ006656,IZ006814,IZ006924,IZ007013,IZ007081,IZ007094,IZ007204,IZ007245,IZ007382,IZ007424,IZ007442,IZ007569,IZ007677,IZ007965,IZ007970,IZ007974,IZ008194,IZ008327,IZ008440,IZ008606,IZ008730,IZ008982,IZ009005,IZ009012,IZ009063,IZ009098,IZ009265,IZ009360,IZ009581,IZ009594,IZ009736,IZ009744,IZ009760,IZ009799,IZ009825,IZ009927,IZ010123,IZ010182,IZ010186,IZ010191,IZ010221,IZ010315,IZ010515,IZ010629,IZ010701,IZ010813,IZ010831,IZ010836,IZ010910,IZ010983,IZ011154,IZ011415,IZ011509,IZ011595,IZ011826,IZ011920,IZ011929,IZ011981,IZ011990,IZ012011,IZ012171,IZ012529,IZ012648,IZ012862,IZ012926,IZ013117,IZ013206,IZ013281,IZ013556,IZ013600,IZ013722,IZ013946,IZ014045,IZ014147,IZ014220,IZ014404,IZ014447,IZ014515,IZ014575,IZ014623,IZ014670,IZ014896 |
| response to stimulus | 133 | 0.236586366 | 0.614469141 | IZ000049,IZ000131,IZ000171,IZ000218,IZ000616,IZ000702,IZ000832,IZ001146,IZ001275,IZ001762,IZ001784,IZ001899,IZ002125,IZ002233,IZ002251,IZ002289,IZ002353,IZ002439,IZ002491,IZ002585,IZ002614,IZ003413,IZ003631,IZ003737,IZ003858,IZ004112,IZ004194,IZ004491,IZ004500,IZ004558,IZ004643,IZ004755,IZ004758,IZ004898,IZ005315,IZ005353,IZ005471,IZ005641,IZ005700,IZ005704,IZ005705,IZ005716,IZ005748,IZ005782,IZ006009,IZ006197,IZ006335,IZ006467,IZ006524,IZ006709,IZ006718,IZ006814,IZ006918,IZ006924,IZ007013,IZ007081,IZ007092,IZ007094,IZ007424,IZ007569,IZ007677,IZ007880,IZ007970,IZ007974,IZ008124,IZ008403,IZ008511,IZ008600,IZ008730,IZ008982,IZ009005,IZ009102,IZ009140,IZ009265,IZ009360,IZ009413,IZ009581,IZ009644,IZ009736,IZ009744,IZ009810,IZ009825,IZ009927,IZ009936,IZ010039,IZ010158,IZ010186,IZ010191,IZ010218,IZ010221,IZ010235,IZ010256,IZ010361,IZ010535,IZ010629,IZ010813,IZ010836,IZ010983,IZ011001,IZ011045,IZ011595,IZ011738,IZ011929,IZ011981,IZ011990,IZ012000,IZ012018,IZ012171,IZ012220,IZ012363,IZ012529,IZ012953,IZ013117,IZ013169,IZ013362,IZ013426,IZ013466,IZ013537,IZ013600,IZ013719,IZ013722,IZ013806,IZ013946,IZ014053,IZ014147,IZ014202,IZ014344,IZ014404,IZ014447,IZ014509,IZ014575,IZ014623,IZ014843 |
| immune system process | 23 | 0.243242117 | 0.614469141 | IZ000832,IZ002289,IZ002353,IZ003631,IZ004500,IZ005471,IZ005716,IZ006009,IZ006709,IZ007013,IZ007424,IZ007880,IZ008600,IZ009927,IZ012018,IZ012171,IZ013362,IZ013719,IZ013806,IZ014053,IZ014344,IZ014404,IZ014843 |
| locomotion | 34 | 0.255232178 | 0.614469141 | IZ000218,IZ000596,IZ000616,IZ001762,IZ001784,IZ002353,IZ002491,IZ002585,IZ003472,IZ003631,IZ004112,IZ004194,IZ004491,IZ004558,IZ004643,IZ005353,IZ005748,IZ006335,IZ007081,IZ007424,IZ007569,IZ008555,IZ008982,IZ009005,IZ009102,IZ009140,IZ009384,IZ009774,IZ009927,IZ010039,IZ010813,IZ011981,IZ013537,IZ013662 |
| membrane part | 50 | 0.284413582 | 0.817172971 | IZ000608,IZ000702,IZ000832,IZ001687,IZ001784,IZ003047,IZ003053,IZ003192,IZ003631,IZ003737,IZ003954,IZ004146,IZ004194,IZ004491,IZ004558,IZ004643,IZ005353,IZ005545,IZ005704,IZ005705,IZ005942,IZ006656,IZ006814,IZ007013,IZ007081,IZ007094,IZ007382,IZ007965,IZ008982,IZ009005,IZ009063,IZ009098,IZ009744,IZ009927,IZ010123,IZ010191,IZ010813,IZ010831,IZ010910,IZ010983,IZ011509,IZ011826,IZ011981,IZ012011,IZ012171,IZ013946,IZ014045,IZ014220,IZ014404,IZ014447 |
| molecular transducer activity | 7 | 0.28724647 | 0.64568187 | IZ006718,IZ007094,IZ010039,IZ010218,IZ010983,IZ013946,IZ014575 |
| nucleic acid binding transcription factor activity | 9 | 0.287713165 | 0.64568187 | IZ000171,IZ000218,IZ000452,IZ007092,IZ007098,IZ007617,IZ012166,IZ012953,IZ013719 |
| structural molecule activity | 14 | 0.295423756 | 0.66042608 | IZ000570,IZ001384,IZ002491,IZ003631,IZ004500,IZ006009,IZ008511,IZ010158,IZ011752,IZ011981,IZ012162,IZ013117,IZ013169,IZ013806 |
| reproductive process | 61 | 0.355656574 | 0.670078839 | IZ000218,IZ000452,IZ000616,IZ000673,IZ000832,IZ001347,IZ001729,IZ001745,IZ001762,IZ002353,IZ002585,IZ003394,IZ003472,IZ003631,IZ003829,IZ004112,IZ004194,IZ004491,IZ004558,IZ004643,IZ004730,IZ005315,IZ005353,IZ005458,IZ006335,IZ006415,IZ006524,IZ006619,IZ006718,IZ006918,IZ007013,IZ007569,IZ007880,IZ007970,IZ008403,IZ008555,IZ008600,IZ009005,IZ009102,IZ009140,IZ009265,IZ009360,IZ009927,IZ010039,IZ010191,IZ010545,IZ010701,IZ010813,IZ010983,IZ012220,IZ012363,IZ012953,IZ013112,IZ013426,IZ013466,IZ013537,IZ014053,IZ014147,IZ014317,IZ014404,IZ014471 |
| catalytic activity | 154 | 0.399075953 | 0.718116616 | IZ000049,IZ000131,IZ000236,IZ000608,IZ000638,IZ000673,IZ000832,IZ000974,IZ001108,IZ001347,IZ001437,IZ001687,IZ001784,IZ001794,IZ001846,IZ001899,IZ002066,IZ002125,IZ002233,IZ002251,IZ002289,IZ002353,IZ002491,IZ002585,IZ002627,IZ002925,IZ002988,IZ003053,IZ003245,IZ003394,IZ003413,IZ003446,IZ003546,IZ003630,IZ003737,IZ003954,IZ004038,IZ004112,IZ004146,IZ004194,IZ004200,IZ004491,IZ004601,IZ004609,IZ004755,IZ004758,IZ004853,IZ004898,IZ004977,IZ005118,IZ005147,IZ005316,IZ005420,IZ005432,IZ005458,IZ005700,IZ005704,IZ005705,IZ005748,IZ006197,IZ006335,IZ006415,IZ006467,IZ006575,IZ006709,IZ006718,IZ006773,IZ006814,IZ006918,IZ006924,IZ007013,IZ007094,IZ007204,IZ007424,IZ007569,IZ007575,IZ007602,IZ007750,IZ007800,IZ007826,IZ007880,IZ007898,IZ007965,IZ007970,IZ007974,IZ007994,IZ008194,IZ008555,IZ008600,IZ008606,IZ008730,IZ008982,IZ009063,IZ009098,IZ009265,IZ009360,IZ009370,IZ009384,IZ009413,IZ009422,IZ009522,IZ009644,IZ009691,IZ009760,IZ009799,IZ009927,IZ009936,IZ010129,IZ010186,IZ010191,IZ010218,IZ010221,IZ010235,IZ010315,IZ010400,IZ010545,IZ010629,IZ010831,IZ010836,IZ010983,IZ011001,IZ011045,IZ011112,IZ011308,IZ011738,IZ011990,IZ012000,IZ012171,IZ012220,IZ012420,IZ012647,IZ012648,IZ012953,IZ012988,IZ013112,IZ013426,IZ013466,IZ013615,IZ013704,IZ013826,IZ013891,IZ013946,IZ014045,IZ014053,IZ014147,IZ014202,IZ014283,IZ014317,IZ014447,IZ014510,IZ014623,IZ014631,IZ014644,IZ014896 |
| single-organism process | 234 | 0.471165152 | 0.741644921 | IZ000049,IZ000131,IZ000171,IZ000218,IZ000318,IZ000452,IZ000570,IZ000596,IZ000608,IZ000616,IZ000638,IZ000673,IZ000702,IZ000832,IZ000974,IZ001057,IZ001275,IZ001347,IZ001390,IZ001437,IZ001687,IZ001729,IZ001745,IZ001762,IZ001784,IZ001794,IZ001898,IZ001899,IZ002125,IZ002145,IZ002233,IZ002251,IZ002328,IZ002353,IZ002439,IZ002491,IZ002585,IZ002614,IZ002627,IZ002637,IZ002764,IZ002988,IZ003047,IZ003053,IZ003394,IZ003413,IZ003446,IZ003472,IZ003630,IZ003631,IZ003737,IZ003829,IZ003858,IZ003902,IZ003954,IZ004112,IZ004146,IZ004194,IZ004491,IZ004500,IZ004558,IZ004601,IZ004609,IZ004643,IZ004678,IZ004730,IZ004755,IZ004758,IZ004898,IZ005044,IZ005315,IZ005316,IZ005353,IZ005420,IZ005432,IZ005458,IZ005471,IZ005520,IZ005545,IZ005641,IZ005655,IZ005700,IZ005704,IZ005705,IZ005716,IZ005748,IZ005782,IZ005942,IZ006009,IZ006197,IZ006335,IZ006415,IZ006524,IZ006575,IZ006619,IZ006644,IZ006656,IZ006709,IZ006718,IZ006773,IZ006814,IZ006918,IZ006924,IZ007013,IZ007015,IZ007081,IZ007094,IZ007382,IZ007424,IZ007569,IZ007575,IZ007602,IZ007617,IZ007750,IZ007780,IZ007800,IZ007826,IZ007880,IZ007898,IZ007925,IZ007945,IZ007965,IZ007970,IZ007974,IZ007994,IZ008124,IZ008199,IZ008403,IZ008511,IZ008515,IZ008555,IZ008600,IZ008606,IZ008730,IZ008982,IZ009005,IZ009063,IZ009098,IZ009102,IZ009140,IZ009265,IZ009360,IZ009370,IZ009384,IZ009413,IZ009420,IZ009522,IZ009581,IZ009594,IZ009644,IZ009691,IZ009744,IZ009760,IZ009774,IZ009799,IZ009810,IZ009927,IZ009936,IZ010039,IZ010123,IZ010158,IZ010177,IZ010182,IZ010186,IZ010191,IZ010218,IZ010221,IZ010235,IZ010256,IZ010315,IZ010400,IZ010545,IZ010629,IZ010701,IZ010813,IZ010831,IZ010983,IZ011001,IZ011112,IZ011287,IZ011308,IZ011415,IZ011509,IZ011738,IZ011752,IZ011826,IZ011929,IZ011981,IZ011990,IZ012000,IZ012011,IZ012018,IZ012171,IZ012220,IZ012363,IZ012647,IZ012862,IZ012953,IZ013112,IZ013117,IZ013169,IZ013281,IZ013362,IZ013426,IZ013466,IZ013537,IZ013600,IZ013615,IZ013662,IZ013670,IZ013719,IZ013722,IZ013806,IZ013946,IZ014045,IZ014053,IZ014147,IZ014171,IZ014202,IZ014220,IZ014283,IZ014317,IZ014344,IZ014404,IZ014447,IZ014471,IZ014509,IZ014575,IZ014623,IZ014644,IZ014670,IZ014719,IZ014843,IZ014896 |
| positive regulation of biological process | 60 | 0.48339786 | 0.744738013 | IZ000171,IZ000596,IZ000616,IZ000832,IZ001275,IZ001762,IZ001784,IZ002233,IZ002289,IZ002328,IZ002439,IZ002585,IZ002637,IZ003472,IZ003858,IZ004194,IZ004491,IZ004500,IZ004601,IZ004643,IZ004758,IZ004898,IZ005353,IZ005471,IZ005748,IZ006009,IZ006814,IZ007013,IZ007081,IZ007424,IZ007569,IZ007617,IZ008124,IZ008555,IZ008600,IZ009005,IZ009102,IZ009140,IZ009265,IZ009360,IZ009384,IZ009413,IZ009420,IZ009581,IZ009774,IZ009811,IZ009927,IZ009936,IZ010039,IZ010191,IZ010256,IZ010629,IZ010813,IZ011981,IZ012171,IZ013537,IZ013719,IZ013806,IZ014053,IZ014404 |
| negative regulation of biological process | 52 | 0.534858806 | 0.768859534 | IZ000171,IZ000616,IZ000632,IZ000638,IZ001762,IZ001784,IZ002233,IZ003472,IZ003631,IZ003902,IZ004194,IZ004491,IZ004601,IZ004730,IZ004758,IZ004898,IZ005471,IZ005716,IZ005748,IZ006619,IZ006709,IZ006918,IZ007013,IZ007424,IZ007569,IZ007617,IZ007880,IZ008124,IZ008403,IZ008600,IZ009102,IZ009140,IZ009360,IZ009413,IZ009581,IZ009811,IZ009927,IZ009936,IZ010039,IZ010158,IZ011415,IZ011929,IZ012018,IZ012171,IZ013362,IZ013426,IZ013537,IZ013719,IZ014053,IZ014171,IZ014344,IZ014843 |
| localization | 88 | 0.564951039 | 0.787821633 | IZ000171,IZ000318,IZ000570,IZ000616,IZ000702,IZ000832,IZ001762,IZ001784,IZ001899,IZ002233,IZ002353,IZ002491,IZ002585,IZ002614,IZ002627,IZ002764,IZ003047,IZ003053,IZ003472,IZ003631,IZ003737,IZ003954,IZ004146,IZ004194,IZ004491,IZ004500,IZ004558,IZ004758,IZ004898,IZ005471,IZ005545,IZ005641,IZ005704,IZ005705,IZ005748,IZ005942,IZ006009,IZ006619,IZ006718,IZ006924,IZ007013,IZ007015,IZ007081,IZ007569,IZ007880,IZ007945,IZ007965,IZ007970,IZ008124,IZ008403,IZ008515,IZ008730,IZ008982,IZ009063,IZ009098,IZ009140,IZ009360,IZ009413,IZ009420,IZ009522,IZ009581,IZ009744,IZ009927,IZ009936,IZ010123,IZ010158,IZ010186,IZ010191,IZ010629,IZ011308,IZ011509,IZ011738,IZ011826,IZ011929,IZ011981,IZ012862,IZ013281,IZ013537,IZ013662,IZ013670,IZ013719,IZ013806,IZ014045,IZ014147,IZ014220,IZ014447,IZ014575,IZ014644 |
| binding | 107 | 0.578105837 | 0.790861632 | IZ000131,IZ000171,IZ000570,IZ000616,IZ000638,IZ000673,IZ000832,IZ001108,IZ001275,IZ001347,IZ001437,IZ001729,IZ001745,IZ001762,IZ001784,IZ001899,IZ002125,IZ002233,IZ002353,IZ002439,IZ002585,IZ002637,IZ003053,IZ003394,IZ003631,IZ003737,IZ003829,IZ003858,IZ004146,IZ004194,IZ004200,IZ004491,IZ004643,IZ004758,IZ004887,IZ004898,IZ005044,IZ005315,IZ005316,IZ005353,IZ005471,IZ005509,IZ005641,IZ005782,IZ006415,IZ006718,IZ006814,IZ006924,IZ007013,IZ007081,IZ007424,IZ007617,IZ007750,IZ007880,IZ007898,IZ007970,IZ008194,IZ008403,IZ008511,IZ008600,IZ009005,IZ009102,IZ009140,IZ009360,IZ009370,IZ009413,IZ009420,IZ009422,IZ009581,IZ009760,IZ009811,IZ009927,IZ009936,IZ010039,IZ010123,IZ010129,IZ010158,IZ010177,IZ010191,IZ010221,IZ010256,IZ010361,IZ010545,IZ010629,IZ010813,IZ010831,IZ011506,IZ011752,IZ011981,IZ012011,IZ012238,IZ012648,IZ013112,IZ013169,IZ013466,IZ013600,IZ013615,IZ013662,IZ013719,IZ013891,IZ013946,IZ014045,IZ014053,IZ014171,IZ014317,IZ014404,IZ014719 |
| reproduction | 67 | 0.645247663 | 0.835266759 | IZ000218,IZ000452,IZ000596,IZ000616,IZ000673,IZ000832,IZ001347,IZ001390,IZ001729,IZ001745,IZ001762,IZ002353,IZ002585,IZ003192,IZ003394,IZ003472,IZ003631,IZ003829,IZ004112,IZ004194,IZ004491,IZ004558,IZ004643,IZ004730,IZ005315,IZ005353,IZ005458,IZ006335,IZ006415,IZ006524,IZ006619,IZ006718,IZ006918,IZ007013,IZ007569,IZ007880,IZ007970,IZ008403,IZ008555,IZ008600,IZ009005,IZ009102,IZ009140,IZ009265,IZ009360,IZ009581,IZ009774,IZ009927,IZ010039,IZ010158,IZ010191,IZ010545,IZ010701,IZ010813,IZ010983,IZ012220,IZ012363,IZ012953,IZ013112,IZ013426,IZ013466,IZ013537,IZ014053,IZ014147,IZ014317,IZ014404,IZ014471 |
| molecular function regulator | 11 | 0.676061105 | 0.864104591 | IZ001899,IZ003472,IZ004491,IZ004758,IZ007210,IZ009581,IZ009927,IZ010361,IZ012171,IZ014053,IZ014510 |
| multicellular organismal process | 108 | 0.781756422 | 0.897248999 | IZ000171,IZ000218,IZ000452,IZ000596,IZ000616,IZ000832,IZ001390,IZ001729,IZ001745,IZ001762,IZ002353,IZ002491,IZ002585,IZ003047,IZ003394,IZ003446,IZ003472,IZ003631,IZ003737,IZ003829,IZ003902,IZ004112,IZ004194,IZ004491,IZ004500,IZ004558,IZ004643,IZ004730,IZ004758,IZ004898,IZ005044,IZ005353,IZ005458,IZ005471,IZ005520,IZ005641,IZ005700,IZ005782,IZ006009,IZ006335,IZ006524,IZ006644,IZ006718,IZ006918,IZ006924,IZ007013,IZ007424,IZ007569,IZ007602,IZ007617,IZ007826,IZ007880,IZ007925,IZ007970,IZ007994,IZ008511,IZ008555,IZ008600,IZ008730,IZ008982,IZ009005,IZ009102,IZ009140,IZ009265,IZ009360,IZ009384,IZ009413,IZ009522,IZ009581,IZ009644,IZ009774,IZ009927,IZ010039,IZ010123,IZ010158,IZ010177,IZ010182,IZ010186,IZ010191,IZ010221,IZ010545,IZ010629,IZ010813,IZ010831,IZ011112,IZ011287,IZ011308,IZ011738,IZ011752,IZ011929,IZ011981,IZ012171,IZ012220,IZ012953,IZ013112,IZ013117,IZ013466,IZ013537,IZ013670,IZ013719,IZ013806,IZ014053,IZ014317,IZ014404,IZ014447,IZ014471,IZ014575,IZ014670 |
| metabolic process | 213 | 0.793152411 | 0.900756234 | IZ000049,IZ000131,IZ000171,IZ000218,IZ000236,IZ000452,IZ000608,IZ000616,IZ000632,IZ000638,IZ000673,IZ000832,IZ000974,IZ001057,IZ001108,IZ001275,IZ001347,IZ001437,IZ001687,IZ001729,IZ001762,IZ001784,IZ001794,IZ001802,IZ001846,IZ001899,IZ002066,IZ002125,IZ002233,IZ002251,IZ002289,IZ002328,IZ002353,IZ002439,IZ002491,IZ002585,IZ002627,IZ002925,IZ002988,IZ003053,IZ003245,IZ003394,IZ003413,IZ003446,IZ003472,IZ003546,IZ003630,IZ003737,IZ003858,IZ003902,IZ003954,IZ004038,IZ004112,IZ004146,IZ004194,IZ004200,IZ004491,IZ004601,IZ004609,IZ004678,IZ004730,IZ004755,IZ004758,IZ004853,IZ004887,IZ004898,IZ004977,IZ005044,IZ005046,IZ005118,IZ005147,IZ005315,IZ005316,IZ005420,IZ005432,IZ005458,IZ005471,IZ005655,IZ005700,IZ005704,IZ005705,IZ005716,IZ005748,IZ005942,IZ006197,IZ006335,IZ006415,IZ006467,IZ006575,IZ006619,IZ006709,IZ006718,IZ006773,IZ006814,IZ006918,IZ006924,IZ007013,IZ007092,IZ007094,IZ007098,IZ007204,IZ007210,IZ007382,IZ007424,IZ007569,IZ007575,IZ007602,IZ007617,IZ007750,IZ007780,IZ007800,IZ007826,IZ007880,IZ007898,IZ007965,IZ007970,IZ007974,IZ007994,IZ008194,IZ008403,IZ008555,IZ008600,IZ008606,IZ008730,IZ008982,IZ009063,IZ009098,IZ009102,IZ009140,IZ009265,IZ009360,IZ009370,IZ009384,IZ009413,IZ009422,IZ009522,IZ009581,IZ009594,IZ009644,IZ009691,IZ009760,IZ009799,IZ009811,IZ009927,IZ009936,IZ010039,IZ010129,IZ010186,IZ010191,IZ010218,IZ010221,IZ010235,IZ010256,IZ010315,IZ010361,IZ010400,IZ010545,IZ010629,IZ010831,IZ010836,IZ010983,IZ011001,IZ011045,IZ011112,IZ011308,IZ011415,IZ011738,IZ011929,IZ011981,IZ011990,IZ012000,IZ012018,IZ012166,IZ012171,IZ012220,IZ012363,IZ012420,IZ012647,IZ012648,IZ012953,IZ012988,IZ013112,IZ013169,IZ013362,IZ013426,IZ013466,IZ013615,IZ013662,IZ013704,IZ013719,IZ013722,IZ013826,IZ013891,IZ013946,IZ014045,IZ014053,IZ014147,IZ014171,IZ014202,IZ014283,IZ014317,IZ014344,IZ014404,IZ014447,IZ014471,IZ014510,IZ014575,IZ014623,IZ014631,IZ014644,IZ014719,IZ014843,IZ014896 |
| developmental process | 110 | 0.805406393 | 0.907410207 | IZ000171,IZ000218,IZ000452,IZ000596,IZ000616,IZ000832,IZ001390,IZ001729,IZ001745,IZ001762,IZ001794,IZ002145,IZ002353,IZ002491,IZ002585,IZ003047,IZ003446,IZ003472,IZ003631,IZ003737,IZ003829,IZ003902,IZ004112,IZ004194,IZ004491,IZ004500,IZ004643,IZ004730,IZ004758,IZ004898,IZ005315,IZ005353,IZ005458,IZ005471,IZ005520,IZ005641,IZ005700,IZ005782,IZ006009,IZ006335,IZ006524,IZ006619,IZ006644,IZ006718,IZ006773,IZ006924,IZ007013,IZ007424,IZ007569,IZ007602,IZ007617,IZ007826,IZ007880,IZ007925,IZ007970,IZ007994,IZ008403,IZ008511,IZ008555,IZ008600,IZ008730,IZ008982,IZ009005,IZ009102,IZ009140,IZ009265,IZ009360,IZ009384,IZ009413,IZ009522,IZ009581,IZ009644,IZ009774,IZ009927,IZ009936,IZ010039,IZ010123,IZ010158,IZ010177,IZ010182,IZ010191,IZ010629,IZ010701,IZ010813,IZ010831,IZ010983,IZ011112,IZ011287,IZ011308,IZ011738,IZ011752,IZ011929,IZ011981,IZ012171,IZ012220,IZ012953,IZ013112,IZ013117,IZ013466,IZ013537,IZ013662,IZ013670,IZ013719,IZ013806,IZ014053,IZ014317,IZ014447,IZ014471,IZ014575,IZ014670 |
| cellular component organization or biogenesis | 103 | 0.807002807 | 0.907410207 | IZ000171,IZ000570,IZ000608,IZ000616,IZ000638,IZ000673,IZ000832,IZ001057,IZ001347,IZ001762,IZ001794,IZ002233,IZ002353,IZ002439,IZ002491,IZ002585,IZ002627,IZ003053,IZ003394,IZ003472,IZ003631,IZ003737,IZ003858,IZ004194,IZ004491,IZ004500,IZ004601,IZ004609,IZ004643,IZ004758,IZ004898,IZ005315,IZ005316,IZ005353,IZ005471,IZ005520,IZ005748,IZ006009,IZ006415,IZ006575,IZ006619,IZ006656,IZ006718,IZ006773,IZ006918,IZ006924,IZ007013,IZ007081,IZ007424,IZ007569,IZ007750,IZ007880,IZ008199,IZ008403,IZ008515,IZ008600,IZ008730,IZ009005,IZ009102,IZ009140,IZ009265,IZ009360,IZ009370,IZ009420,IZ009522,IZ009581,IZ009644,IZ009760,IZ009811,IZ009927,IZ009936,IZ010039,IZ010123,IZ010158,IZ010182,IZ010186,IZ010218,IZ010256,IZ010545,IZ010629,IZ010701,IZ010813,IZ010983,IZ011308,IZ011752,IZ011929,IZ011981,IZ011990,IZ012011,IZ012363,IZ013169,IZ013426,IZ013662,IZ013719,IZ013806,IZ014045,IZ014053,IZ014147,IZ014171,IZ014404,IZ014447,IZ014471,IZ014719 |
| multi-organism process | 46 | 0.81892764 | 0.915034573 | IZ000171,IZ000218,IZ000616,IZ000832,IZ001729,IZ001762,IZ002233,IZ002289,IZ002491,IZ002585,IZ003394,IZ003472,IZ003631,IZ004500,IZ004558,IZ004898,IZ005315,IZ005458,IZ005748,IZ006009,IZ006619,IZ006709,IZ006718,IZ006918,IZ007013,IZ007569,IZ007677,IZ007880,IZ009102,IZ009140,IZ009360,IZ009736,IZ009927,IZ010039,IZ010545,IZ010701,IZ010983,IZ011990,IZ012171,IZ012529,IZ013112,IZ013426,IZ013537,IZ013806,IZ014317,IZ014404 |
| transporter activity | 22 | 0.880045755 | 0.973299118 | IZ000702,IZ001784,IZ002614,IZ002764,IZ003047,IZ003954,IZ004146,IZ005545,IZ005704,IZ005705,IZ005942,IZ006619,IZ007015,IZ007945,IZ007965,IZ009063,IZ009098,IZ010191,IZ011509,IZ013281,IZ014220,IZ014575 |
| cell junction | 12 | 0.908298427 | 1 | IZ000638,IZ000832,IZ003631,IZ005704,IZ007204,IZ007424,IZ009140,IZ009581,IZ011981,IZ011990,IZ013117,IZ013600 |
| macromolecular complex | 76 | 0.950236999 | 1 | IZ000049,IZ000171,IZ000452,IZ000608,IZ000638,IZ000832,IZ001057,IZ001347,IZ001729,IZ001745,IZ001762,IZ002353,IZ002439,IZ002491,IZ002585,IZ003472,IZ003631,IZ003737,IZ003829,IZ003858,IZ003902,IZ004146,IZ004194,IZ004491,IZ004601,IZ004758,IZ004898,IZ005147,IZ005458,IZ005471,IZ005700,IZ005748,IZ006415,IZ006524,IZ006718,IZ006918,IZ006924,IZ007013,IZ007081,IZ007424,IZ007569,IZ007750,IZ007880,IZ008403,IZ008515,IZ008600,IZ009102,IZ009140,IZ009265,IZ009360,IZ009370,IZ009413,IZ009522,IZ009581,IZ009927,IZ009936,IZ010158,IZ010191,IZ010256,IZ010361,IZ010629,IZ010831,IZ011308,IZ011509,IZ012011,IZ012171,IZ013112,IZ013169,IZ013537,IZ013662,IZ013946,IZ014147,IZ014171,IZ014220,IZ014447,IZ014719 |
| growth | 38 | 0.958103319 | 0.987273465 | IZ000049,IZ000171,IZ000596,IZ000616,IZ000673,IZ002585,IZ003339,IZ003446,IZ003472,IZ003631,IZ004194,IZ004491,IZ004563,IZ004643,IZ005316,IZ005353,IZ006619,IZ007013,IZ007602,IZ007826,IZ008555,IZ009005,IZ009140,IZ009360,IZ009760,IZ009774,IZ009927,IZ009936,IZ010039,IZ010123,IZ010158,IZ010813,IZ011738,IZ011929,IZ013117,IZ013426,IZ013537,IZ014053 |
| membrane-enclosed lumen | 45 | 0.993108722 | 1 | IZ000131,IZ000171,IZ000452,IZ000638,IZ001275,IZ001916,IZ002233,IZ002328,IZ002439,IZ002585,IZ003394,IZ003858,IZ003902,IZ004194,IZ004491,IZ004601,IZ004758,IZ004898,IZ005471,IZ005655,IZ006524,IZ006918,IZ007013,IZ007081,IZ007750,IZ007994,IZ008403,IZ008555,IZ008600,IZ009102,IZ009370,IZ009522,IZ009927,IZ009936,IZ010191,IZ010256,IZ010545,IZ011415,IZ012171,IZ013117,IZ013662,IZ013719,IZ014171,IZ014623,IZ014719 |

**Table S18** KEGG enrichment of the specific families genes identified in *I. galbana*

| KEGG_A_class | KEGG_B_class | Pathway | number | p-value | q-value | Genes_id |
| --- | --- | --- | --- | --- | --- | --- |
| Metabolism | Global and overview maps | Metabolic pathways | 63 | 0.9998778 | 1.00E+00 | IZ005147;IZ007539;IZ007575;IZ007622;IZ008927;IZ008982;IZ010315;IZ012758;IZ013573;IZ014317;IZ014471;IZ001789;IZ002251;IZ004088;IZ004678;IZ005420;IZ005443;IZ005700;IZ005951;IZ006575;IZ007780;IZ007800;IZ007898;IZ008110;IZ008284;IZ008555;IZ008606;IZ009413;IZ009594;IZ009691;IZ009799;IZ009825;IZ010019;IZ010181;IZ010186;IZ010221;IZ010235;IZ010983;IZ011112;IZ011154;IZ011170;IZ011407;IZ011960;IZ012420;IZ013826;IZ013869;IZ013953;IZ014703;IZ014896;IZ000896;IZ000900;IZ001123;IZ002066;IZ002615;IZ002689;IZ002971;IZ003559;IZ003903;IZ004853;IZ004854;IZ004856;IZ006079;IZ007033 |
| Cellular Processes | Cellular community - eukaryotes | Signaling pathways regulating pluripotency of stem cells | 36 | 6.79E-14 | 1.83E-11 | IZ000011;IZ004985;IZ007620;IZ011666;IZ014246;IZ001169;IZ001176;IZ001306;IZ003637;IZ004490;IZ005472;IZ008873;IZ009782;IZ010039;IZ010124;IZ010858;IZ011307;IZ012917;IZ012975;IZ013230;IZ013947;IZ014236;IZ014893;IZ001639;IZ001670;IZ002190;IZ002205;IZ002435;IZ002620;IZ002630;IZ003049;IZ003391;IZ003825;IZ004547;IZ004851;IZ005514 |
| Metabolism | Global and overview maps | Biosynthesis of secondary metabolites | 23 | 0.9978796 | 1.00E+00 | IZ005147;IZ007622;IZ014317;IZ001789;IZ002251;IZ005443;IZ005951;IZ007800;IZ008110;IZ009594;IZ009691;IZ010181;IZ010186;IZ010221;IZ011154;IZ011170;IZ011407;IZ011960;IZ013869;IZ014071;IZ000900;IZ001123;IZ003903 |
| Environmental Information Processing | Signal transduction | Calcium signaling pathway | 14 | 0.000131245 | 3.94E-03 | IZ001241;IZ005750;IZ007210;IZ008049;IZ009360;IZ009363;IZ011287;IZ011475;IZ012000;IZ013891;IZ014703;IZ003103;IZ005782;IZ006858 |
| Environmental Information Processing | Signal transduction | cGMP - PKG signaling pathway | 13 | 0.000503943 | 9.74E-03 | IZ001241;IZ005750;IZ006899;IZ007210;IZ007827;IZ011475;IZ014163;IZ002996;IZ003103;IZ005782;IZ006493;IZ006494;IZ006858 |
| Environmental Information Processing | Signal transduction | Apelin signaling pathway | 13 | 0.005861459 | 4.65E-02 | IZ000184;IZ014447;IZ001241;IZ004491;IZ005750;IZ007013;IZ008049;IZ009360;IZ010039;IZ011475;IZ003103;IZ005782;IZ006858 |
| Cellular Processes | Cell growth and death | Cellular senescence | 12 | 0.005050398 | 4.26E-02 | IZ001241;IZ002233;IZ004310;IZ005409;IZ005750;IZ007013;IZ007210;IZ009265;IZ011475;IZ003103;IZ005782;IZ006858 |
| Environmental Information Processing | Signal transduction | PI3K-Akt signaling pathway | 12 | 0.01206058 | 7.75E-02 | IZ000184;IZ014447;IZ004310;IZ004491;IZ005407;IZ007013;IZ007941;IZ008274;IZ008479;IZ009265;IZ013719;IZ014151 |
| Cellular Processes | Cell growth and death | Oocyte meiosis | 11 | 0.01937009 | 1.11E-01 | IZ001241;IZ004310;IZ005750;IZ007210;IZ008049;IZ009265;IZ009360;IZ011475;IZ003103;IZ005782;IZ006858 |
| Environmental Information Processing | Signal transduction | Ras signaling pathway | 10 | 0.00092285 | 1.31E-02 | IZ014447;IZ001241;IZ005750;IZ008049;IZ009360;IZ011475;IZ014053;IZ003103;IZ005782;IZ006858 |
| Genetic Information Processing | Folding, sorting and degradation | Ubiquitin mediated proteolysis | 10 | 0.07507997 | 2.57E-01 | IZ005186;IZ011738;IZ001899;IZ002289;IZ004038;IZ006170;IZ010928;IZ014510;IZ014631;IZ003790 |
| Metabolism | Global and overview maps | Microbial metabolism in diverse environments | 10 | 0.9688042 | 1.00E+00 | IZ005147;IZ005443;IZ009691;IZ010221;IZ011112;IZ011154;IZ011170;IZ011407;IZ003903;IZ004856 |
| Cellular Processes | Transport and catabolism | Lysosome | 9 | 0.07617341 | 2.57E-01 | IZ001762;IZ001802;IZ007826;IZ008284;IZ013953;IZ002689;IZ002971;IZ003446;IZ003559 |
| Cellular Processes | Transport and catabolism | Autophagy - animal | 9 | 0.1193933 | 3.79E-01 | IZ000184;IZ013466;IZ013600;IZ002289;IZ004491;IZ007013;IZ008049;IZ009360;IZ012127 |
| Cellular Processes | Transport and catabolism | Endocytosis | 9 | 0.1276832 | 3.92E-01 | IZ000251;IZ008981;IZ010315;IZ013600;IZ002289;IZ009420;IZ010983;IZ013152;IZ004194 |
| Cellular Processes | Cellular community - eukaryotes | Gap junction | 8 | 0.00521216 | 4.26E-02 | IZ006899;IZ007827;IZ008049;IZ009360;IZ014163;IZ002996;IZ006493;IZ006494 |
| Environmental Information Processing | Signal transduction | MAPK signaling pathway | 8 | 0.0147888 | 9.07E-02 | IZ002289;IZ004755;IZ007210;IZ008049;IZ009360;IZ009384;IZ014053;IZ014202 |
| Environmental Information Processing | Signal transduction | cAMP signaling pathway | 8 | 0.02273147 | 1.23E-01 | IZ001241;IZ005750;IZ008049;IZ009360;IZ011475;IZ003103;IZ005782;IZ006858 |
| Environmental Information Processing | Signal transduction | Phosphatidylinositol signaling system | 8 | 0.02945939 | 1.45E-01 | IZ001241;IZ005750;IZ010983;IZ011475;IZ014703;IZ003103;IZ005782;IZ006858 |
| Environmental Information Processing | Signal transduction | FoxO signaling pathway | 8 | 0.06970324 | 2.51E-01 | IZ000184;IZ004310;IZ004491;IZ005407;IZ005409;IZ009265;IZ010039;IZ014053 |
| Genetic Information Processing | Translation | Ribosome biogenesis in eukaryotes | 8 | 0.1837875 | 4.90E-01 | IZ007994;IZ008403;IZ014147;IZ014719;IZ001437;IZ002939;IZ005526;IZ006484 |
| Metabolism | Global and overview maps | Biosynthesis of amino acids | 8 | 0.8826951 | 9.75E-01 | IZ014317;IZ005443;IZ009691;IZ010221;IZ011154;IZ011407;IZ000900;IZ001123 |
| Environmental Information Processing | Signal transduction | Rap1 signaling pathway | 7 | 0.03339436 | 1.61E-01 | IZ001241;IZ005750;IZ011475;IZ013946;IZ003103;IZ005782;IZ006858 |
| Environmental Information Processing | Membrane transport | ABC transporters | 7 | 0.03807654 | 1.68E-01 | IZ009098;IZ001784;IZ003954;IZ005704;IZ007965;IZ012529;IZ004146 |
| Cellular Processes | Cellular community - eukaryotes | Tight junction | 7 | 0.06101152 | 2.32E-01 | IZ000184;IZ005242;IZ013600;IZ004491;IZ008049;IZ009360;IZ000941 |
| Genetic Information Processing | Folding, sorting and degradation | Protein processing in endoplasmic reticulum | 7 | 0.7877422 | 9.51E-01 | IZ013466;IZ014404;IZ001275;IZ006656;IZ008626;IZ012127;IZ014623 |
| Metabolism | Global and overview maps | Carbon metabolism | 7 | 0.9821946 | 1.00E+00 | IZ009691;IZ010221;IZ011112;IZ011154;IZ011170;IZ003903;IZ004856 |
| Environmental Information Processing | Signal transduction | MAPK signaling pathway - plant | 6 | 0.008309882 | 6.02E-02 | IZ001241;IZ005750;IZ011475;IZ003103;IZ005782;IZ006858 |
| Metabolism | Carbohydrate metabolism | Starch and sucrose metabolism | 6 | 0.1664142 | 4.68E-01 | IZ008927;IZ001789;IZ002251;IZ007800;IZ011960;IZ007033 |
| Genetic Information Processing | Transcription | Spliceosome | 6 | 0.9430094 | 9.96E-01 | IZ013662;IZ001269;IZ005458;IZ013089;IZ000784;IZ001608 |
| Cellular Processes | Cell growth and death | Apoptosis - fly | 5 | 0.04041389 | 1.70E-01 | IZ002289;IZ004755;IZ005409;IZ009384;IZ014202 |
| Metabolism | Glycan biosynthesis and metabolism | Other glycan degradation | 5 | 0.07358762 | 2.55E-01 | IZ008284;IZ013953;IZ002689;IZ002971;IZ003559 |
| Metabolism | Lipid metabolism | Sphingolipid metabolism | 5 | 0.1306145 | 3.96E-01 | IZ008284;IZ013953;IZ002689;IZ002971;IZ003559 |
| Cellular Processes | Cellular community - prokaryotes | Quorum sensing | 5 | 0.186962 | 4.90E-01 | IZ008327;IZ008372;IZ014045;IZ001123;IZ003053 |
| Environmental Information Processing | Signal transduction | AMPK signaling pathway | 5 | 0.5248771 | 8.75E-01 | IZ000184;IZ013600;IZ004491;IZ004678;IZ007013 |
| Environmental Information Processing | Signal transduction | Hedgehog signaling pathway | 4 | 0.0503047 | 2.03E-01 | IZ007569;IZ008049;IZ009360;IZ001678 |
| Environmental Information Processing | Signal transduction | Wnt signaling pathway | 4 | 0.0717238 | 2.51E-01 | IZ007210;IZ008049;IZ009360;IZ010039 |
| Environmental Information Processing | Signal transduction | Hippo signaling pathway | 4 | 0.0717238 | 2.51E-01 | IZ004755;IZ009384;IZ010039;IZ014202 |
| Cellular Processes | Cell growth and death | p53 signaling pathway | 4 | 0.08391837 | 2.73E-01 | IZ004310;IZ005409;IZ005700;IZ009265 |
| Cellular Processes | Cell growth and death | Apoptosis | 4 | 0.3874336 | 7.69E-01 | IZ005409;IZ007826;IZ012127;IZ003446 |
| Metabolism | Lipid metabolism | Glycerolipid metabolism | 4 | 0.4271822 | 7.71E-01 | IZ000684;IZ008606;IZ009799;IZ014896 |
| Metabolism | Carbohydrate metabolism | Pyruvate metabolism | 4 | 0.4271822 | 7.71E-01 | IZ007898;IZ011112;IZ013826;IZ003903 |
| Metabolism | Energy metabolism | Methane metabolism | 4 | 0.4271822 | 7.71E-01 | IZ009691;IZ011154;IZ011170;IZ004856 |
| Genetic Information Processing | Folding, sorting and degradation | Protein export | 4 | 0.4468225 | 7.89E-01 | IZ008327;IZ014045;IZ003053;IZ004609 |
| Cellular Processes | Transport and catabolism | Autophagy - yeast | 4 | 0.5043422 | 8.51E-01 | IZ013466;IZ007013;IZ008049;IZ009360 |
| Cellular Processes | Transport and catabolism | Peroxisome | 4 | 0.5411983 | 8.86E-01 | IZ010547;IZ008256;IZ010629;IZ004146 |
| Metabolism | Amino acid metabolism | Glycine, serine and threonine metabolism | 4 | 0.6104529 | 9.37E-01 | IZ005443;IZ009691;IZ011154;IZ011407 |
| Cellular Processes | Cell growth and death | Cell cycle | 4 | 0.7975988 | 9.51E-01 | IZ004310;IZ005409;IZ009265;IZ010039 |
| Metabolism | Carbohydrate metabolism | Glycolysis / Gluconeogenesis | 4 | 0.8076613 | 9.54E-01 | IZ009691;IZ010221;IZ011154;IZ011170 |
| Genetic Information Processing | Translation | Ribosome | 4 | 0.9938363 | 1.00E+00 | IZ000370;IZ012710;IZ001057;IZ006524 |

**Table S19** Differentially expressed genes in the comparison of C7d vs. T7d by transcriptome data

| gene_id | log2FoldChange | lfcSE | stat | pvalue | padj |
| --- | --- | --- | --- | --- | --- |
| IZ000921 | -0.618228045 | 0.28330285 | -2.182216116 | 0.029093586 | 0.128349004 |
| IZ000920 | -0.862878253 | 0.395637297 | -2.180983083 | 0.029184668 | 0.128651816 |
| IZ010869 | -0.019082564 | 0.513613945 | -0.037153517 | 0.970362601 | 0.987349924 |
| IZ011827 | -0.011840673 | 0.34486052 | -0.034334672 | 0.972610277 | 0.98799762 |
| IZ010564 | -0.772164862 | 0.382123114 | -2.02072273 | 0.043308475 | 0.165837741 |
| IZ003505 | -0.300845842 | 0.657648064 | -0.4574572 | 0.647342459 | 0.817137655 |
| IZ008009 | -0.46717575 | 0.49530133 | -0.943215215 | 0.345570833 | 0.581678454 |
| IZ010260 | -0.654490375 | 0.30151432 | -2.170677585 | 0.029955553 | 0.130988806 |
| IZ006832 | 0.358276546 | 0.41590108 | 0.861446541 | 0.388992153 | 0.620149672 |
| IZ012322 | -1.041756606 | 0.374929781 | -2.7785379 | 0.005460414 | 0.042540685 |
| IZ000195 | -1.812689526 | 1.357545286 | -1.335270023 | 0.181787986 | 0.402025948 |
| IZ002967 | 0.095797365 | 0.520894459 | 0.183909357 | 0.854084564 | 0.935664555 |
| IZ007183 | 0.129312115 | 0.50400017 | 0.256571569 | 0.797509525 | 0.907113004 |
| IZ009688 | 1.78319659 | 0.298305616 | 5.97775065 | 2.26E-09 | 3.38E-07 |
| IZ001744 | -1.692288335 | 0.47018672 | -3.599183607 | 0.000319218 | 0.005373199 |
| IZ003288 | -0.726128197 | 0.452481446 | -1.604768998 | 0.108544651 | 0.294645988 |
| IZ000291 | 0.254309903 | 0.288428783 | 0.881707784 | 0.377934851 | 0.610612329 |
| IZ013688 | 1.393393923 | 0.378958137 | 3.67690725 | 0.000236079 | 0.004268348 |
| IZ013190 | -0.215916559 | 0.283158015 | -0.762530274 | 0.445743582 | 0.667805133 |
| IZ002979 | 2.110430225 | 0.414719263 | 5.088816496 | 3.60E-07 | 2.74E-05 |
| IZ005918 | -2.151611413 | 0.423190094 | -5.084266964 | 3.69E-07 | 2.76E-05 |
| IZ001188 | 0.163039038 | 0.373436404 | 0.436591174 | 0.662407864 | 0.827300281 |
| IZ008332 | 1.281168966 | 0.461675097 | 2.775044558 | 0.005519415 | 0.0427694 |
| IZ007195 | -1.15478264 | 0.575781124 | -2.005593084 | 0.04489968 | 0.169885644 |
| IZ006149 | 1.375698459 | 0.302553454 | 4.546960017 | 5.44E-06 | 0.000239136 |
| IZ007042 | -0.637016402 | 0.29061089 | -2.191990816 | 0.02838017 | 0.126337578 |
| IZ007738 | 0.905071091 | 0.322767895 | 2.804092678 | 0.00504584 | 0.040526097 |
| IZ012603 | 1.494212381 | 0.284789023 | 5.246734463 | 1.55E-07 | 1.38E-05 |
| IZ007325 | -0.076755196 | 0.329380162 | -0.233029202 | 0.81573873 | 0.916852587 |
| IZ011107 | -0.545539022 | 0.307710975 | -1.772894262 | 0.07624623 | 0.237586114 |
| IZ005291 | -0.3532401 | 0.314010222 | -1.124931851 | 0.260617914 | 0.493797145 |
| IZ003473 | -0.090417255 | 0.376020193 | -0.240458509 | 0.809974825 | 0.913979253 |
| IZ004535 | 1.205586609 | 0.407258941 | 2.960245895 | 0.003073936 | 0.028450653 |
| IZ008735 | 0.616690163 | 0.304361504 | 2.026176621 | 0.042746689 | 0.16426549 |
| IZ009303 | 0.99199501 | 0.317468966 | 3.124699155 | 0.00177987 | 0.019397025 |
| IZ007092 | -1.657513516 | 0.276271983 | -5.999571508 | 1.99E-09 | 1.32E-07 |
| IZ013676 | 1.152823461 | 0.319632713 | 3.606713001 | 0.0003101 | 0.005271705 |

**Table S20 Raw data of carotenoids compound content detected by HPLC**

| Compound | 7d-W1 | 7d-W2 | 7d-W3 | 7d-G1 | 7d-G2 | 7d-G3 | Pvalue | FoldChange | Log2FC |
| --- | --- | --- | --- | --- | --- | --- | --- | --- | --- |
| β-Carotene | 202.74 | 251.97 | 296.51 | 439.63 | 393.78 | 397.05 | 0.0129 | 1.6379 | 0.7119 |
| ε-Carotene | 0.32 | 0.45 | 0.64 | 1.18 | 1.07 | 0.99 | 0.0088 | 2.2854 | 1.1925 |
| Lutein myristate | 0.07 | 0.08 | 0.07 | 0.19 | 0.08 | 0.07 | 0.3977 | 1.5650 | 0.6461 |
| Violaxanthin laurate | 0.02 | 0.03 | 0.04 | 0.08 | 0.10 | 0.11 | 0.0031 | 3.5034 | 1.8087 |
| Violaxanthin myristate | 13.05 | 11.83 | 11.55 | 43.48 | 24.11 | 26.54 | 0.0865 | 2.5839 | 1.3695 |
| Zeaxanthin myristoleate | 0.04 | 0.05 | 0.05 | N/A | N/A | N/A | N/A | N/A | N/A |
| Zeaxanthin palmitate | 3.50 | 3.11 | 2.34 | 0.85 | 1.18 | 1.31 | 0.0201 | 0.3740 | -1.4189 |
| Antheraxanthin | 0.26 | 0.43 | 0.50 | 0.98 | 1.19 | 1.24 | 0.0026 | 2.8554 | 1.5137 |
| Apocarotenal | 0.05 | 0.03 | 0.04 | 0.08 | 0.06 | 0.06 | 0.0641 | 1.5979 | 0.6762 |
| Canthaxanthin | 0.18 | 0.11 | 0.11 | 0.26 | 0.25 | 0.21 | 0.0254 | 1.8273 | 0.8697 |
| Capsanthin | 0.10 | 0.11 | 0.10 | 0.43 | 0.19 | 0.18 | 0.1822 | 2.5528 | 1.3521 |
| Echinenone | 35.18 | 24.02 | 21.73 | 45.79 | 32.51 | 33.32 | 0.1622 | 1.3792 | 0.4639 |
| Zeaxanthin | 8.35 | 7.30 | 12.04 | 28.16 | 22.62 | 23.14 | 0.0029 | 2.6692 | 1.4164 |
| β-Cryptoxanthin | 12.38 | 7.28 | 5.99 | 14.47 | 9.33 | 9.52 | 0.3791 | 1.2983 | 0.3766 |
| Fucoxanthin | 2.83 | 2.12 | 2.75 | 5.28 | 5.49 | 6.01 | 0.0091 | 2.1476 | 1.1028 |

**Table S21** Differentially accumulated carotenoids compounds in the comparison of 7d-W vs. 7d-G by targeted metabolomics (n=3)

| Compound | Class | Pvalue | FoldChange | Log2FoldChange |  |
| --- | --- | --- | --- | --- | --- |
| ε-Carotene | carotenes | 0.0088 | 2.2854 | 1.1925 | up |
| violaxanthin laurate | carotenoid esters | 0.0031 | 3.5034 | 1.8087 | up |
| violaxanthin myristate | carotenoid esters | 0.0087 | 2.5839 | 1.3695 | up |
| zeaxanthin palmitate | carotenoid esters | 0.0201 | 0.3740 | -1.4189 | down |
| antheraxanthin | xanthophylls | 0.0026 | 2.8554 | 1.5137 | up |
| capsanthin | xanthophylls | 0.0018 | 2.5528 | 1.3521 | up |
| zeaxanthin | xanthophylls | 0.0029 | 2.6692 | 1.4164 | up |
| fucoxanthin | xanthophylls | 0.0091 | 2.1476 | 1.1028 | up |

**Table S22** Statistical analysis of transcriptome data

| Sample | Repeat | Total_reads | Total_bases | GC_content | Q20 | Q30 |
| --- | --- | --- | --- | --- | --- | --- |
| C9d | C_9_r1 | 67,821,102 | 10,173,165,300 | 60.12% | 96.11% | 90.83% |
|  | C_9_r2 | 74,208,330 | 11,131,249,500 | 60.33% | 96.10% | 90.82% |
|  | C_9_r3 | 83,696,476 | 12,554,471,400 | 60.12% | 96.14% | 90.85% |
| C7d | C_7_r1 | 108,882,342 | 16,332,351,300 | 60.70% | 95.50% | 89.52% |
|  | C_7_r2 | 90,029,112 | 13,504,366,800 | 60.14% | 95.83% | 90.21% |
|  | C_7_r3 | 75,430,952 | 11,314,642,800 | 60.14% | 95.97% | 90.52% |
| C5d | C_5_r1 | 109,665,534 | 16,449,830,100 | 60.49% | 96.67% | 92.29% |
|  | C_5_r2 | 93,155,890 | 13,973,383,500 | 60.26% | 95.61% | 89.76% |
|  | C_5_r3 | 83,308,448 | 12,496,267,200 | 59.98% | 95.96% | 90.48% |
| C3d | C_3_r1 | 82,271,478 | 12,340,721,700 | 60.11% | 96.11% | 90.82% |
|  | C_3_r2 | 85,417,812 | 12,812,671,800 | 59.83% | 96.66% | 92.01% |
|  | C_3_r3 | 92,750,238 | 13,912,535,700 | 60.13% | 97.01% | 93.02% |
| T9d | T-9_r1 | 82,128,488 | 12,319,273,200 | 60.07% | 96.26% | 91.01% |
|  | T-9_r2 | 76,520,248 | 11,478,037,200 | 60.30% | 94.51% | 87.56% |
|  | T-9_r3 | 91,676,842 | 13,751,526,300 | 56.83% | 96.18% | 90.83% |
| T7d | T-7_r1 | 75,864,106 | 11,379,615,900 | 60.06% | 96.38% | 91.33% |
|  | T-7_r2 | 77,804,744 | 11,670,711,600 | 60.01% | 96.19% | 90.93% |
|  | T-7_r3 | 66,379,212 | 9,956,881,800 | 60.42% | 96.82% | 92.32% |
| T5d | T-5_r1 | 82,685,654 | 12,402,848,100 | 60.07% | 96.27% | 91.14% |
|  | T-5_r2 | 83,617,610 | 12,542,641,500 | 60.47% | 97.05% | 92.83% |
|  | T-5_r3 | 65,585,916 | 9,837,887,400 | 60.15% | 95.98% | 90.44% |
| T3d | T-3_r1 | 118,223,716 | 17,733,557,400 | 59.75% | 96.96% | 92.55% |
|  | T-3_r2 | 109,416,042 | 16,412,406,300 | 60.52% | 96.38% | 91.34% |
|  | T-3_r3 | 75,262,158 | 11,289,323,700 | 60.02% | 95.86% | 90.21% |

**Table S23** Mapping data of each transcriptome sample to the generated genome assembly

| Sample | Repeat | Total_reads | Unique_mapped | Multiple_mapped | Unmapped |
| --- | --- | --- | --- | --- | --- |
| C9d | C_9_r1 | 33910551 | 93.62 | 1.54 | 4.84 |
|  | C_9_r2 | 37104165 | 93.45 | 1.52 | 5.03 |
|  | C_9_r3 | 41848238 | 93.41 | 1.5 | 5.09 |
| C7d | C_7_r1 | 54441171 | 92.25 | 1.79 | 5.96 |
|  | C_7_r2 | 45014556 | 92.68 | 1.5 | 5.82 |
|  | C_7_r3 | 37715476 | 93.38 | 1.77 | 4.85 |
| C5d | C_5_r1 | 54832767 | 92.76 | 1.7 | 5.54 |
|  | C_5_r2 | 46577945 | 93.34 | 1.68 | 4.98 |
|  | C_5_r3 | 41654224 | 92.6 | 1.93 | 5.47 |
| C3d | C_3_r1 | 41135739 | 93.56 | 1.42 | 5.02 |
|  | C_3_r2 | 42708906 | 93.48 | 1.45 | 5.07 |
|  | C_3_r3 | 46375119 | 93.63 | 1.61 | 4.76 |
| T9d | T_9_r1 | 41064244 | 93.68 | 1.64 | 4.68 |
|  | T_9_r2 | 38260124 | 92.53 | 1.65 | 5.82 |
|  | T_9_r3 | 45838421 | 91.11 | 1.84 | 7.05 |
| T7d | T_7_r1 | 37932053 | 93.96 | 1.61 | 4.43 |
|  | T_7_r2 | 38902372 | 93.37 | 1.59 | 5.04 |
|  | T_7_r3 | 33189606 | 93.24 | 1.81 | 4.95 |
| T5d | T_5_r1 | 41342827 | 93.83 | 1.52 | 4.65 |
|  | T_5_r2 | 41808805 | 93.61 | 1.68 | 4.71 |
|  | T_5_r3 | 32792958 | 93.35 | 1.6 | 5.05 |
| T3d | T_3_r1 | 59111858 | 93.65 | 1.72 | 4.63 |
|  | T_3_r2 | 54708021 | 93.16 | 1.65 | 5.19 |
|  | T_3_r3 | 37631079 | 93.47 | 1.45 | 5.08 |
