## Supplementary figures and images for "Multi-omics Analyses Provide Insight into the Biosynthesis Pathways of Fucoxanthin in *Isochrysis galbana*"

### Graphical abstract.jpeg

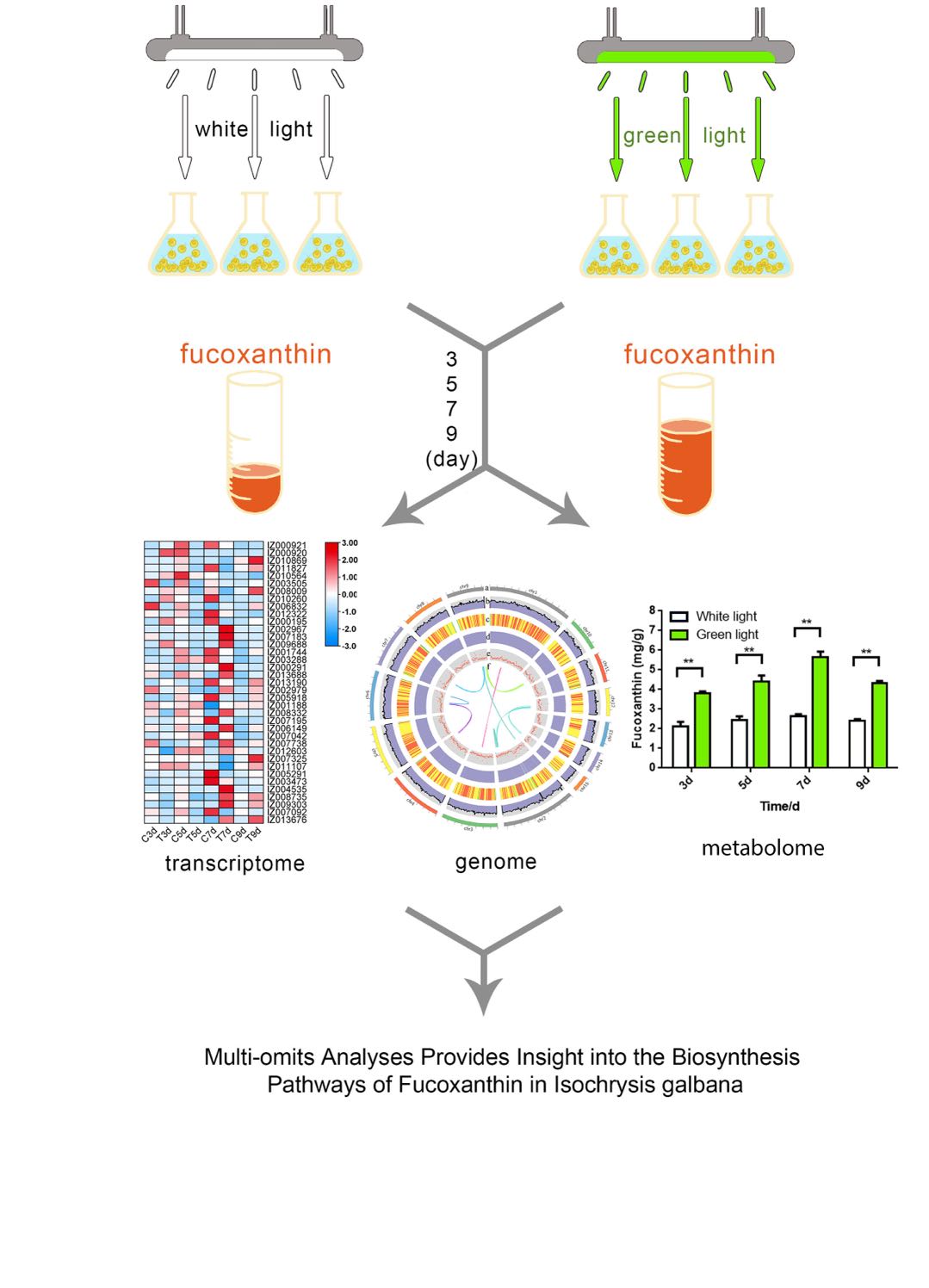
